# Orthogonal Chemical Proteomic Strategies Reveal the Cholesteryl Ester Interactome in Mammalian Cells

**DOI:** 10.64898/2026.09.21.753363

**Authors:** Aakash Chandramouli, Kavita Sharma, Adithya Kallattu, Chaitanya Katkar, Manish Deshmukh, Pooja Thakral, Harinath Chakrapani, Siddhesh S. Kamat

## Abstract

Cholesteryl esters (CEs) are widely regarded as inert storage forms of cholesterol, yet their potential to directly engage cellular proteins remains largely unexplored. Here, we establish an integrated chemical proteomic platform to systematically define CE–protein interactions in mammalian cells. We developed and comparatively deployed three orthogonal photoaffinity labeling strategies: (i) metabolic assembly of a bifunctional CE probe through endogenous acyl-CoA:cholesterol acyltransferase activity, (ii) direct delivery of a structurally defined diazirine–alkyne CE analog, and (iii) fragment-assisted subtraction using sterol and fatty acyl control probes to resolve interactions dependent on the intact esterified scaffold. Each probe system was rigorously validated by lipidomics and UV-dependent crosslinking prior to quantitative proteomic analysis. Integration of these complementary modalities identified 495 CE-associated proteins spanning enzymes, transporters, scaffolding proteins, and canonical sterol-binding families. The limited overlap across strategies reveals that lipid–protein engagement is strongly conditioned by biosynthetic origin, probe topology, and intracellular routing. Functional and database annotations further demonstrate enrichment of druggable and disease-linked proteins, connecting CE interactions to metabolic, neurological, and cardiovascular pathways. Collectively, this work provides the first systems-level map of CE–protein interactions and establishes a generalizable, multimodal framework for chemically resolving lipid–protein interactomes.

## INTRODUCTION

Cells are composed of a diverse repertoire of biomolecules that collectively orchestrate highly regulated and interconnected biochemical processes essential for life. Among these, lipids represent a major class of cellular macromolecules that extend far beyond their traditional roles as structural components of membranes and reservoirs of metabolic energy^1,2^. Lipids actively participate in signal transduction, organelle identity, and protein regulation, thereby influencing fundamental cellular decisions^3^. Disruption of lipid homeostasis is now recognized as a driving force in numerous pathophysiological states, including metabolic disorders, neurodegenerative diseases, and cancer^4^. Consequently, defining lipid-protein interactions has emerged as a critical objective in chemical biology, as such interactions underpin lipid-mediated signaling pathways and offer opportunities for therapeutic intervention.

Cholesterol is an important lipid in mammalian physiology, where it regulates membrane fluidity and organization, serves as a biosynthetic precursor for steroid hormones, bile acids, and vitamin D, and functions as a signaling molecule on its own^5,6^. Cellular cholesterol levels are tightly controlled to prevent toxicity arising from excess free cholesterol. A major mechanism for this regulation is the enzymatic esterification of cholesterol with long-chain fatty acids to form cholesteryl esters (CEs). This process is catalyzed intracellularly by acyl-CoA:cholesterol acyltransferase (ACAT)^7,8^ and in plasma (or blood) by lecithin:cholesterol acyltransferase (LCAT)^9^, while CE hydrolases mediate the reverse reaction to liberate free cholesterol^10^. Through this dynamic interconversion, CEs act as a buffering pool that governs cholesterol storage, transport, and mobilization in cells and tissues.

Perturbations in cholesterol flux and CE metabolism have been strongly implicated in disease^6,11–13^. Accumulation of CEs has been reported in multiple neurodegenerative disorders, including Alzheimer’s disease^14^, Huntington’s disease^15^, and amyotrophic lateral sclerosis^16^, where altered lipid storage is linked to neuronal dysfunction^17,18^. In cancer, enhanced CE accumulation within lipid droplets is increasingly recognized as a metabolic hallmark that supports rapid proliferation and oncogenic signaling^19,20^. Despite their clear association with disease, CEs have long been considered metabolically inert entities, relegated primarily to passive roles in cholesterol storage within lipid droplets and transport within lipoproteins. This perception has begun to shift with emerging evidence that CEs and their oxidized variants can directly engage protein targets and modulate signaling pathways, particularly in contexts like atherosclerosis and inflammation^21^. Nevertheless, in contrast to cholesterol itself, whose protein interactome and signaling functions have been extensively studied, the protein interaction landscape of CEs remains largely unexplored.

Chemical proteomics approaches have proven to be indispensable for mapping lipid-protein interactions in native biological systems^22^. Among these, photoaffinity labeling (PAL) has emerged as a powerful strategy, enabling the covalent capture of transient or low-affinity lipid-protein interactions *in situ*^23–26^. In a conventional PAL workflow, a functionalized lipid probe is designed to closely mimic the native lipid while incorporating a photoreactive crosslinking moiety and a bioorthogonal handle for downstream enrichment and identification (**Figure 1**). Upon UV irradiation, the activated probe forms a covalent bond with proximal protein binding partners, which can then be conjugated to reporter tags and analyzed by various techniques such as SDS-PAGE gel electrophoresis, fluorescence microscopy and quantitative mass spectrometry (**Figure 1**). This strategy has been successfully applied to a wide range of lipid classes, including fatty acids and its signaling derivatives^27–30^, phospholipids^31–35^, sphingolipids^36–38^, and cholesterol^39–41^, yielding valuable insights into lipid signaling networks in cells.

**Figure 1.**
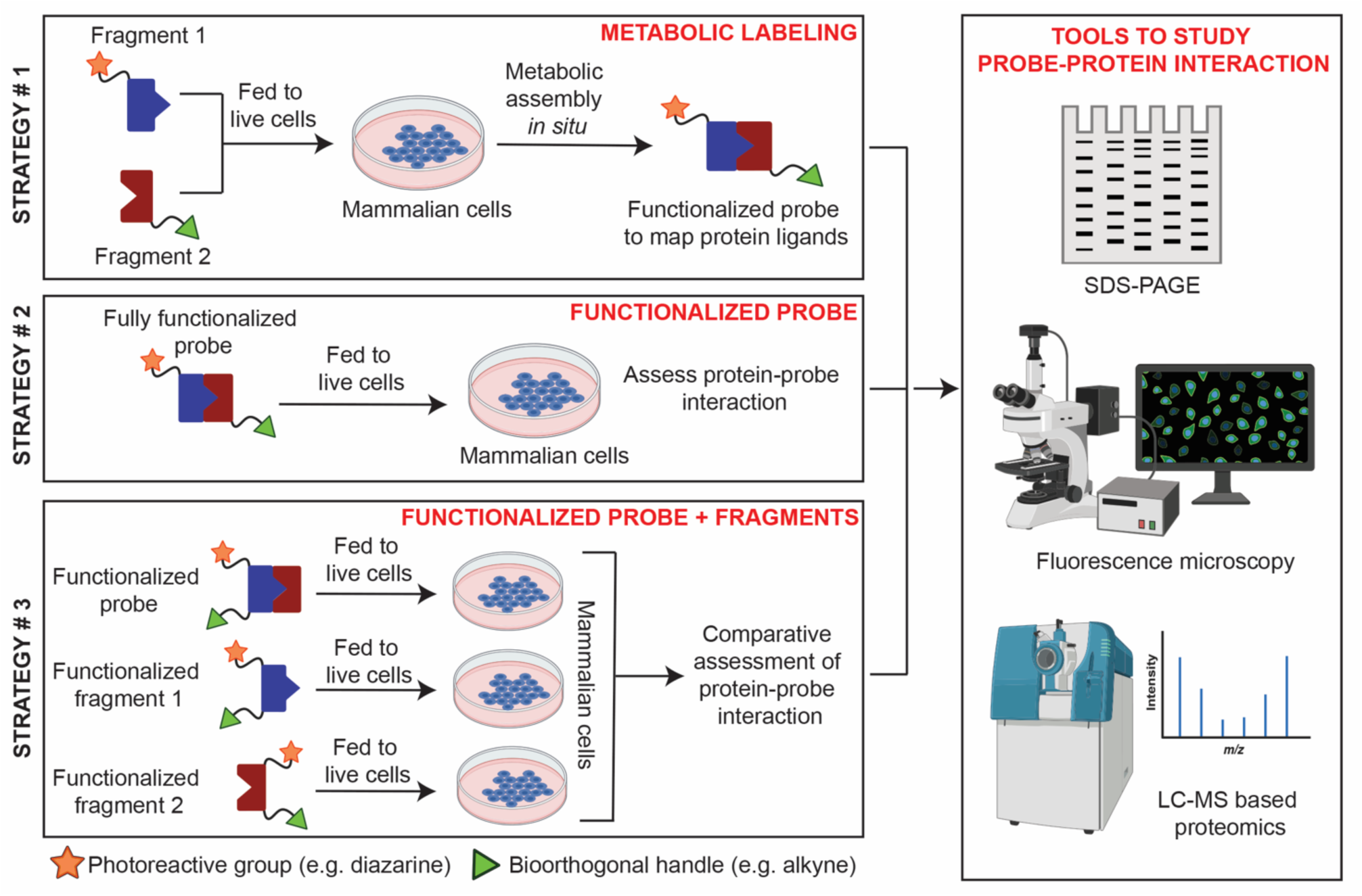
The diverse PAL strategies used to map CE interacting proteins. A schematic representing the three strategies, namely metabolic labeling (strategy # 1), functionalized probe (strategy # 2), and functionalized probe & fragments (strategy # 3), along with the analytical tools to study probe-protein interactions towards identifying protein ligands of CEs in mammalian cells.

Despite its utility, the PAL approach presents notable challenges. The synthesis of functionalized lipid probes is often laborious and challenging, particularly for complex lipid classes, and subtle structural perturbations may compromise biological fidelity. Moreover, the introduction of exogenous probes frequently necessitates extensive control experiments to distinguish bonafide protein interactors from non-specific background binding^42^. To address such limitations, recent efforts have turned toward metabolic labeling strategies^31,35,43–45^, wherein minimally modified lipid precursors are supplied to cells, and enzymatically assembled into endogenous lipid species using the cell’s own biosynthetic machinery (**Figure 1**). This approach preserves native lipid architecture and cellular distribution while enabling chemical proteomic interrogation. Although relatively new, metabolic labeling has been successfully employed to study phospholipid interactions, highlighting its potential as a complementary alternative to functionalized lipid probes.

Here, we complementarily apply both metabolic labeling and tailored probes based PAL strategies to systematically interrogate the protein interactome of CEs (**Figure 1**). We report the development of two distinct types of functionalized CE probes, alongside metabolic labeling probes, targeting both the cholesterol and fatty acid components of CE biosynthesis. These probes are rigorously characterized in a mammalian cell line using in-gel fluorescence, fluorescence microscopy, and lipidomics analysis to confirm cellular uptake, metabolic incorporation, and structural integrity. Leveraging LC-MS/MS based quantitative chemical proteomics workflows, we identify and classify CE interacting proteins across multiple experimental modalities, enabling direct comparison of metabolic labeling versus functionalized probes in the diverse PAL approaches used here. Collectively, this study provides the first comprehensive map of CE interacting proteins and establishes a versatile framework for elucidating the biological functions of this understudied yet disease-relevant lipid class.

## RESULTS

### Metabolic labeling strategy

To systematically map the proteome interacting with CEs, we first adopted the metabolic labeling strategy that leverages endogenous cellular biosynthetic machinery to generate a functionalized CE probe *in situ*. In mammalian cells, CE biosynthesis is catalyzed predominantly by the enzyme ACAT (also known as sterol O-acyltransferase, SOAT)^7,8^, which esterifies excess free cholesterol using medium- to long-chain fatty acyl-CoA substrates. Publicly available transcriptomic^46^ and proteomic^47^ datasets indicate that relative to other cell types, macrophages highly express ACAT. This prompted us to select the RAW264.7 murine macrophages as a candidate mammalian cell line for validating this metabolic labeling strategy.

For metabolic assembly of the functionalized CE probe, we synthesized two probes: (i) an alkyne-functionalized cholesterol analog (Chol-alk, Probe 1), bearing a clickable handle at the iso-octyl side chain (**Figure 2A, Supplementary Scheme 1A, Supplementary Synthetic Note**); and (ii) a diazirine-containing stearic acid analog (18-diaz-FA, Probe 2) (**Figure 2A, Supplementary Scheme 2, Supplementary Synthetic Note**). Synthesis of Probe 1 was carried out in four steps starting from cholenic acid. Protection of the alcohol group of cholenic acid with tetrahydropyran (THP) gave the derivative **1**, which was then reduced using lithium aluminium hydride (LiAlH4) to afford the alcohol, **2** in 74% yield. Treatment of **2** with propargyl bromide gave the THP-protected alkyne **3**, in 72% yield. The final step was cleavage of the THP group, which was done using *para*-toluene sulfonic acid (PTSA) in excellent yield (81%) (**Supplementary Scheme 1A**). On the other hand, Probe 2 was prepared using reported procedures^31^.

**Figure 2.**
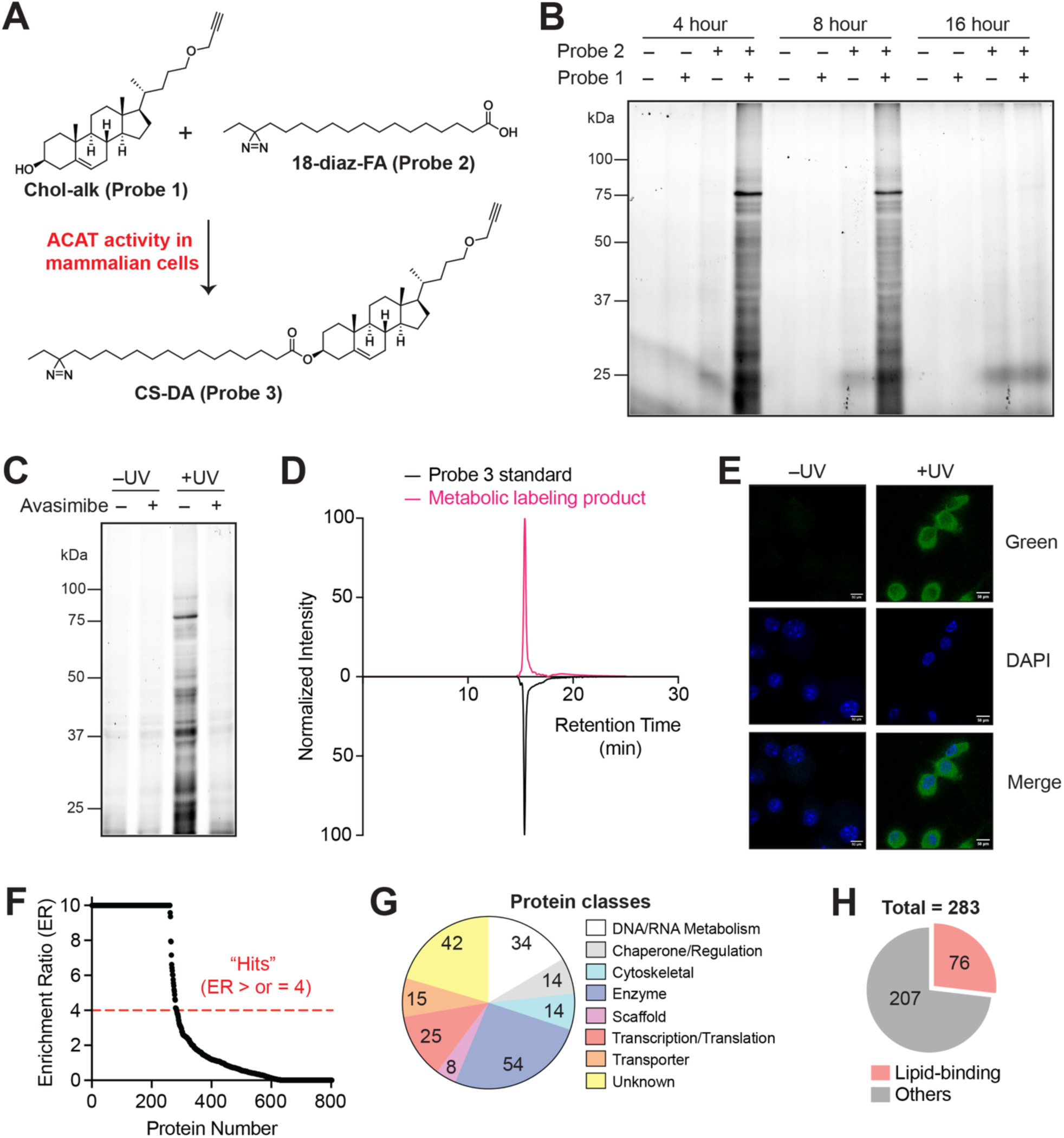
Validation of a metabolic labeling strategy for profiling CE interacting proteins. (**A**) Chemical structures of metabolic precursors, Chol-alk (*<u>Probe 1</u>*) and 18-diaz-FA (*<u>Probe 2</u>*), and the ACAT dependent activity in mammalian cells that converts them to a fully functionalized CE probe, CS-DA (*<u>Probe 3</u>*). (**B**) A representative fluorescence gel from a time-course experiment showing maximum protein labeling via Probe 3 formation when both Probe 1 and Probe 2 (200 μM each), but not the individual probes, are added to RAW264.7 cells, at 4 h relative to 8 h (reduced labeling) and 16 h (no labeling). (**C**) A representative fluorescence gel showing the ACAT dependent protein labeling from metabolically synthesized Probe 3, where RAW264.7 cells treated with vehicle (DMSO) showing intense protein labeling compared to cells treated with the ACAT inhibitor Avasimibe^48^ (25 μM, 4 h) showing reduced labeling similar to no UV controls. (**D**) A LC-MS experiment showing the elution profile of Probe 3 formed from the metabolic labeling strategy in cells co-eluting with the chemically synthesized Probe 3 at the same retention time. (**E**) Representative fluorescence microscopy images showing cellular fluorescence for Probe 3 (green channel) obtained from metabolic labeling strategy when both Probe 1 and Probe 2 (200 μM each, 4 h) are added to RAW264.7 cells only in the presence of UV. In the same experiment, no fluorescence (green channel) was observed in the non-UV irradiated controls. DAPI (blue channel) was used in these fluorescence microscopy experiments to mark the nucleus in individual cells. The scale bars on each panel are 50 μm in length. (**F**) A LC-MS/MS based quantitative proteomics experiment showing the enrichment ratio (ER) (+UV enriched proteins/–UV enriched proteins) of all the proteins identified from the UV-dependent photocrosslinking by metabolically assembled Probe 3, when both Probe 1 and Probe 2 (200 μM each, 4 h) were added from RAW264.7 cells. Each data point represents the average ER for the respective protein from at least three biological replicates. The horizontal dotted line denotes an ER ≥ 4, and proteins having an ER above this threshold were considered as “hits” i.e. enriched by Probe 3, and considered for subsequent analysis. (**G,F**) Categorization of “hits” from the metabolically synthesized Probe 3 into: (**G**) protein classes based on the Panther database classification^53,54^, and (**H**) lipid-binding proteins based on the UniProt annotation^82^. For (**B,C**): The loading control for this gel (Coomassie staining) can be found in the *<u>Supplementary Information</u>*. For (**B-E**): This experiment was done three times with reproducible results each time. For (**F-H**): Complete details for all the “hits” (283 proteins) and categorization can be found in *Supplementary Table 1*.

We hypothesized that upon simultaneously feeding these two probes to RAW264.7 cells, ACAT-mediated enzymatic esterification of these precursors would yield a fully functionalized CE probe [cholesteryl stearate diazirine alkyne (CS-DA), Probe 3] bearing both a photocrosslinkable diazirine and a bioorthogonal alkyne (**Figure 2A**). Critically, this design minimizes non-specific protein labeling, as it ensures that only the fully assembled CE probe (Probe 3), but not the individual fragments (Probe 1 or Probe 2), would result in photocrosslinking of CE interacting proteins and their downstream enrichment via click chemistry.

We first optimized the incubation time required for efficient intracellular synthesis of Probe 3, and its subsequent protein labeling. For this, RAW264.7 cells were treated with Probe 1, Probe 2 or both (200 μM each), under conditions known to stimulate ACAT dependent CE formation.

Subsequently, the treated cells were harvested at different time points (4, 8 or 16 h), UV-irradiated, and analyzed by established gel-based chemical proteomics. In-gel fluorescence analysis revealed maximal protein labeling at 4 h, slightly reduced labeling at 8 h, and complete loss of signal at 16 h (**Figure 2B**). The diminished labeling at extended time points likely reflects cytotoxic effects associated with prolonged exposure to excess free cholesterol or perhaps the dynamic cellular turnover of CEs in RAW264.7 cells. Accordingly, a 4 h incubation was selected as the time point for all subsequent experiments in this study. Importantly, treatment with either Probe 1 or Probe 2 alone, or vehicle control, did not produce any detectable protein labeling, confirming that probe activation requires formation of Probe 3 (**Figure 2B**). Moreover, UV-dependent labeling was strictly required, as demonstrated by the absence of fluorescence in non-UV irradiated samples even when both Probe 1 and Probe 2 were added (**Supplementary Figure 1A**).

To directly establish ACAT dependence, RAW 264.7 cells were pre-treated with a reversible ACAT specific inhibitor Avasimibe^48^ (25 μM, 4 h), following which Probe 1 and Probe 2 (200 μM each, 4 h) were added to these cells to metabolically assemble Probe 3. Inhibition of ACAT markedly reduced probe dependent labeling to levels comparable to no UV controls, whereas robust labeling was observed in vehicle treated cells (**Figure 2C**). To corroborate this, we also performed the same metabolic labeling study in Neuro2A cells, which have low ACAT expression and activity^46,47^. Here, we found negligible fluorescence from gel based chemical proteomics experiments (**Supplementary Figure 1B**), confirming the reduced formation of Probe 3 in cells that have low ACAT activity. Together, these data confirm that intracellular formation of Probe 3 is ACAT activity mediated, and RAW264.7 cells are indeed a suitable mammalian cell line for subsequent studies.

In an effort to benchmark the metabolic labeling strategy, we chemically synthesized Probe 3 to serve as an exogenous reference in our lipidomics analysis. Probe 3, which has both the alkyne handle and the diazirine moiety, was synthesized by esterification of Probe 2 with Probe 1 (**Supplementary Scheme 1B**, **Supplementary Synthetic Note**). LC-MS analysis^10^ of cellular lipid extracts [treated with Probe 1 and Probe 2 (200 µM each, 4 h)] demonstrated that the expected product of metabolic labeling strategy (i.e. Probe 3) exactly co-eluted during chromatographic separation, and matched the retention time of the chemically synthesized Probe 3 standard (**Figure 2D**). Further, only RAW264.7 cells treated with both Probe 1 and Probe 2 (200 μM each, 4 h) exhibited accumulation of the expected functionalized CE probe, whereas cells treated with either precursor alone or vehicle showed no detectable formation of Probe 3 (**Supplementary Figure 2**). This semi-quantitative lipidomics experiment also confirmed efficient cellular uptake of both precursors and their consumption during CE synthesis as part of the metabolic labeling strategy (**Supplementary Figure 2**).

Next, to visualize the probe-derived protein labeling *in situ*, we performed fluorescence microscopy following UV-induced crosslinking and click conjugation to a fluorophore (Alexa Fluor-488). Robust intracellular fluorescence was observed exclusively in RAW264.7 cells treated with both metabolic precursors (200 μM each, 4 h) and exposed to UV light, while no signal was detected in non-UV irradiated cells or in cells treated with individual probes (**Figure 2E, Supplementary Figure 3**). These observations provide additional confirmation of efficient intracellular synthesis of Probe 3 and UV-dependent protein labeling throughout the cell.

Having established optimal protein-labeling conditions via our metabolic labeling strategy, next, we performed LC-MS/MS based quantitative proteomics^29^ to identify CE-interacting proteins. Here, probe-labeled proteomes [after addition of both precursors (200 μM each, 4 h)], with and without UV irradiation, were enriched via click chemistry^49^, processed to generate tryptic peptides, and analyzed using SWATH-MS, an advanced LC-MS/MS based data independent acquisition technique^50–52^. In our analysis, a protein was considered a “hit” (enriched by Probe 3) if it fulfilled the following three criteria: (i) detected in ≥ 3 (out of 6) biological replicates; (ii) had ≥ 3 quantifiable peptides per replicate it was detected in; and (iii) had an enrichment ratio (ER) of ≥ 4 in all replicates it was detected in [ER is a protein’s abundance in UV irradiated samples versus non-irradiated samples, with a maximum value capped at 10). Applying these stringent criteria, we identified a total 283 proteins enriched by the metabolically synthesized Probe 3 from RAW264.7 cells (**Figure 2F, Supplementary Table 1**). Functional classification of these 283 proteins using the PANTHER database^53,54^ revealed broad representation across various protein classes (**Figure 2G, Supplementary Table 1**), and biological activities/processes (**Supplementary Figure 4, Supplementary Table 1**). Enriched proteins spanned diverse processes including lipid homeostasis, metabolic regulation, membrane trafficking, and cellular signaling (**Supplementary Figure 4, Supplementary Table 1)**. Notably, multiple established lipid-binding protein classes (**Figure 2H, Supplementary Table 1**), such as transporters, storage proteins, and vesicular trafficking components, including known sterol binding proteins, were prominently represented in the 283 enriched proteins. Together, our proteomics analysis supports the conclusion that our metabolic labeling strategy to produce the functionalized CE probe (Probe 3), faithfully mimics endogenous CEs, and establishes a robust platform for systematic discovery of CE-protein interactions in mammalian cells having heightened ACAT enzymatic activity.

### Functionalized probe strategy

Having established the metabolic labeling strategy, we next deployed a chemically synthesized functionalized CE probe, Probe 3, to map CE-protein interactions in RAW264.7 macrophages. To define optimal conditions for Probe 3 labeling, RAW264.7 cells were incubated with Probe 3 (200 μM, 4 h), followed by UV irradiation and gel-based chemical proteomic analysis.

In-gel fluorescence profiling revealed robust, strictly UV-dependent protein labeling after 4 h of treatment (**Figure 3A**), establishing this time point for all subsequent Probe 3 cellular experiments.

**Figure 3.**
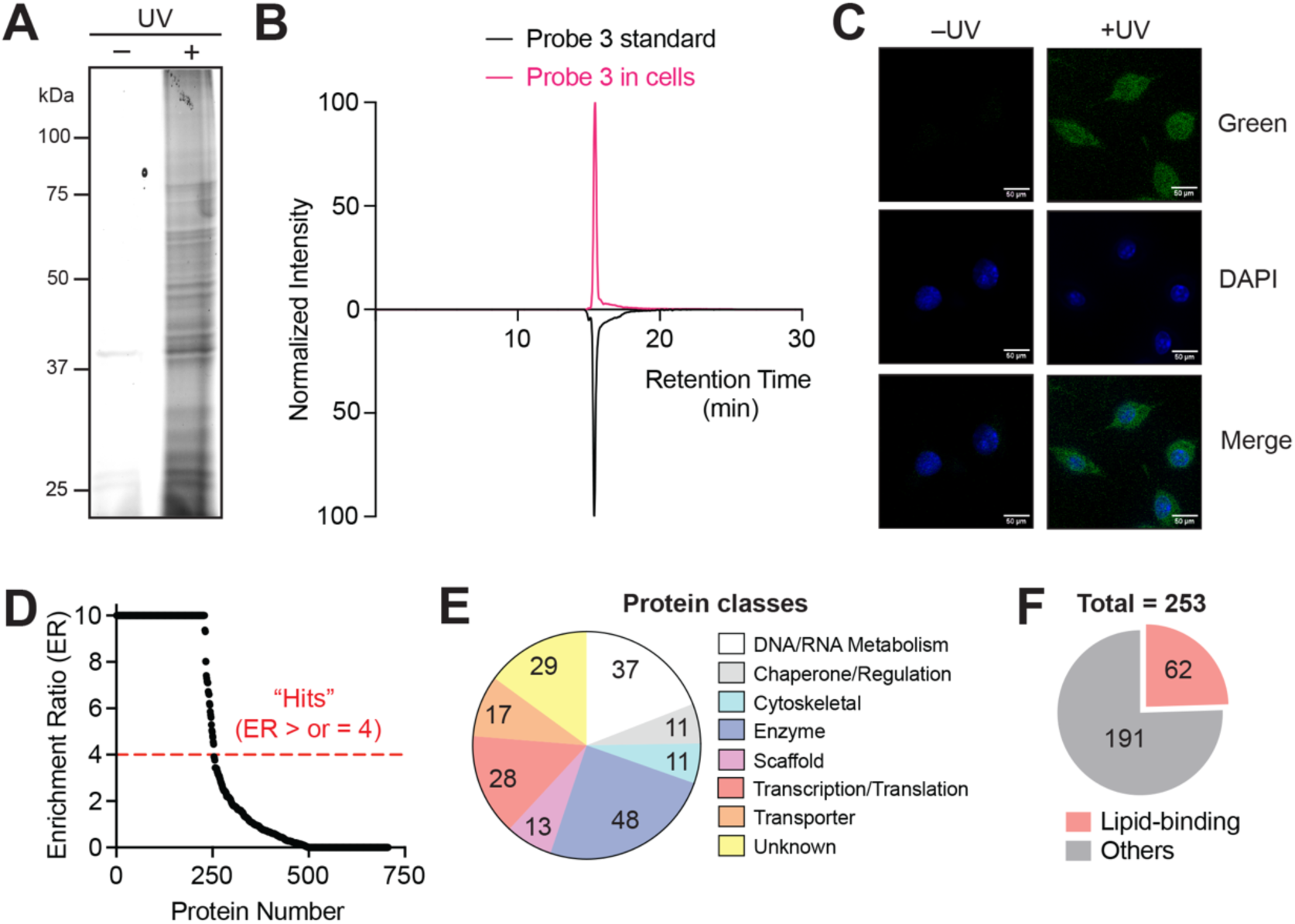
Validation of a functionalized probe for identifying CE interacting proteins. (**A**) A representative fluorescence gel from a gel based chemical proteomics experiment showing the UV-dependent photocrosslinking of the CS-DA probe (*<u>Probe 3</u>*) (200 μM, 4h treatment). (**B**) A LC-MS experiment showing the elution profile of Probe 3 (50 μM, 4 h treatment) extracted from treated RAW264.7 cells co-eluting with the Probe 3 synthesized standard at the same retention time. (**C**) Representative fluorescence microscopy images showing cellular fluorescence for Probe 3 (green channel) fed to RAW264.7 cells (200 μM, 4 h) only in the presence of UV. In the same experiment, no fluorescence (green channel) was observed in the non-UV irradiated controls. DAPI (blue channel) was used in these fluorescence microscopy experiments to mark the nucleus in individual cells. The scale bars on each panel are 50 μm in length. (**D**) A LC-MS/MS based quantitative proteomics experiment showing the enrichment ratio (ER) (+UV enriched proteins/–UV enriched proteins) of all the proteins identified from the UV-dependent photocrosslinking by Probe 3 (50 µM, 4 h treatment), when fed to RAW264.7 cells. Each data point represents the average ER for the respective protein from at least three biological replicates. The horizontal dotted line denotes an ER ≥ 4, and proteins having an ER above this threshold were considered as “hits” i.e. enriched by Probe 3, and considered for subsequent analysis. (**E,F**) Categorization of “hits” from Probe 3 when fed to RAW264.7 cells into: (**E**) protein classes based on the Panther database classification^53,54^, and (**F**) lipid-binding proteins based on the UniProt annotation^82^. For (**A**): The loading control for this gel (Coomassie staining) can be found in the *<u>Supplementary Information</u>*. For (**A-C**): This experiment was done three times with reproducible results each time. For (**D-F**): Complete details for all the “hits” (253 proteins) and categorization can be found in *Supplementary Table 2*.

To confirm cellular uptake and integrity of the functionalized probe, we performed a semi-quantitative LC-MS lipidomics on lipid extracts from Probe 3-treated (50 μM, 4 h) RAW264.7 cells. Probe 3 was efficiently internalized, co-eluted with the authentic chemical standard, and matched its chromatographic retention time (**Figure 3B, Supplementary Figure 5**). No Probe 3 signal was detected in vehicle controls, and hydrolytic degradation products (i.e., Probe 1 or Probe 2) were absent, demonstrating that Probe 3 remains intact under the defined experimental conditions (**Supplementary Figure 5**). Consistent with these findings, fluorescence microscopy following UV photocrosslinking and click conjugation to a fluorophore (Alexa Fluor-488) revealed strong intracellular signal exclusively in Probe 3-treated (200 μM, 4 h), UV-irradiated RAW264.7 cells, with negligible signal in non-irradiated controls (**Figure 3C**). These data collectively confirm efficient uptake, metabolic stability, and UV-dependent proteome labeling by Probe 3.

With optimal labeling conditions established, we applied the same LC–MS/MS-based quantitative proteomics pipeline to identify Probe 3-enriched (50 μM, 4 h) CE-interacting proteins. RAW264.7 proteomes labeled in the presence or absence of UV were enriched by click chemistry^49^, digested to generate tryptic peptides, and interrogated by SWATH-MS^50–52^. Proteins were designated as Probe 3 “hits” if they met three rigorous criteria: (i) detection in ≥ 3 of 4 biological replicates; (ii) have ≥ 3 quantifiable peptides per replicate; and (iii) an enrichment ratio (ER) ≥ 4 in all detected replicates (ER defined as abundance in UV vs non-UV samples, capped at 10, as mentioned earlier). Using these filters, we identified 253 proteins significantly enriched by Probe 3 (**Figure 3D, Supplementary Table 2**). Functional classification via the PANTHER database^53,54^ revealed broad representation across major protein classes (**Figure 3E**), biological activities (**Supplementary Figure 6A**), and physiological processes (**Supplementary Figure 6B**), including lipid homeostasis, metabolic regulation, membrane trafficking, and signaling. Notably, established lipid-binding families, such as lipid transporters, storage proteins, vesicular trafficking factors, and known sterol-binding proteins, were prominently enriched (**Figure 3F, Supplementary Table 2**). These results demonstrate that the functionalized CE probe, Probe 3, serves as an excellent surrogate for endogenous CE behavior, and enables robust chemical proteomic profiling of CE–protein interactions in mammalian cells.

Further, quite interestingly, we find that of the 253 proteins enriched by Probe 3 in this experimental paradigm, only ∼ 50% of them (128 out of 253) are common to this functionalized probe feeding and metabolic labeling strategy (**Supplementary Figure 6C**). This result strongly suggests that metabolically synthesizing the probe in cells versus feeding the same functionalized probe to cells engages different protein pools, presumably due to their differential partitioning or apparent localized concentrations in different cellular compartments. It also emphasizes that multiple strategies are needed to fully sample the complete repertoire of CE-interacting proteins in mammalian cells.

### Functionalized probe with fragments strategy

Using the metabolic labeling and functionalized probe strategy, we identified a broad repertoire of CE-interacting proteins. However, because this study represents the first systematic effort to globally profile CE-interacting proteins, we sought to maximize coverage of potential interactors while minimizing structural perturbations to the native lipid scaffold. CEs can engage proteins through two chemically and spatially distinct regions: the sterol moiety, which includes the iso-octyl side chain, and the esterified fatty acyl chain. In the previous strategies, the iso-octyl arm of cholesterol was modified via an ether linkage to introduce an alkyne handle for click chemistry, potentially biasing protein interactions arising from the sterol region. To complement this strategy and selectively capture CE-specific interactions without modifying the sterol core, we designed another tailored bifunctional CE probe in which the cholesterol moiety remained unaltered.

To this end, we employed a previously reported bifunctional fatty acid probe, palmitic acid diazirine alkyne (PA-DA, Probe 4)^27,29,39^ (**Supplementary Scheme 3, Supplementary Synthetic Note**), and esterified it to cholesterol to generate the cholesteryl palmitate diazirine alkyne (CP-DA, Probe 5) (**Supplementary Scheme 1C, Supplementary Synthetic Note**). In this design, the diazirine and alkyne functionalities are positioned on the fatty acyl chain, enabling UV-induced crosslinking and subsequent click-chemistry based enrichment while preserving the native sterol architecture (**Figure 4A**). We reasoned that Probe 5 would also mimic endogenous CEs and thus potentially expand the repertoire of CE-interacting proteins beyond those accessible by metabolic labeling strategy and the functionalized probe mentioned earlier. Although Probe 5 would potentially yield an expanded interactome, our overarching goal was to define proteins that interact specifically with CEs rather than cholesterol or fatty acids alone. We therefore hypothesized that the increased number of enriched proteins could arise from two other sources: (i) the unmodified iso-octyl side chain of the sterol moiety, which may recruit cholesterol-binding proteins not specific to CEs, and (ii) non-specific interactions originating from the proximity of the diazirine and alkyne functionalities on the fatty acyl chain.

**Figure 4.**
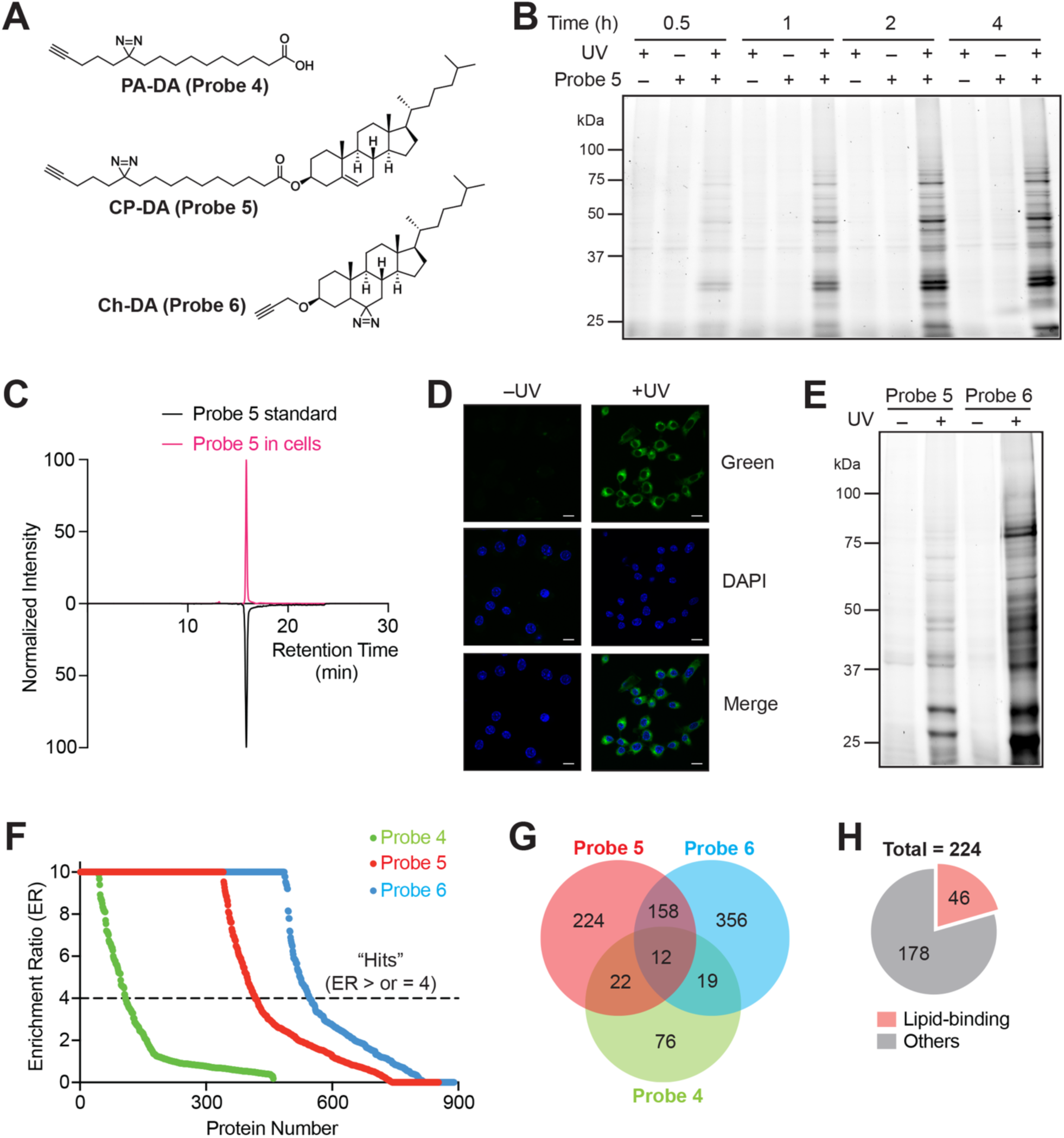
**Validation of a functionalized probe and fragments strategy for identification of CE-interacting proteins**. (**A**) Chemical structures of functionalized probe and fragments, PA-DA (fragment, *<u>Probe 4</u>*), CP-DA (functionalized CE probe, *<u>Probe 5</u>*), and Ch-DA (fragment, *<u>Probe 6</u>*). (**B**) A representative fluorescence gel from a time-course experiment showing linear time-dependent increase in protein labeling when Probe 5 (50 μM) is fed to RAW264.7 cells from 30 min to 4 h, with maximum labeling observed at 4 h timepoint. (**C**) A LC-MS experiment showing the elution profile of Probe 5 (50 μM, 4 h treatment) extracted from treated RAW264.7 cells co-eluting with the Probe 5 synthesized standard at the same retention time. (**D**) Representative fluorescence microscopy images showing cellular fluorescence for Probe 5 (green channel) fed to RAW264.7 cells (50 μM, 4 h) only in the presence of UV. In the same experiment, no fluorescence (green channel) was observed in the non-UV irradiated controls. DAPI (blue channel) was used in these fluorescence microscopy experiments to mark the nucleus in individual cells. The scale bars on each panel are 50 μm in length. (**E**) A representative fluorescence gel from a gel based chemical proteomics experiment showing the UV-dependent photocrosslinking of Probe 5 relative to Probe 6 with Probe 6 showing overall higher labeling intensity (50 μM, 4 h treatment). (**F**) A LC-MS/MS based quantitative proteomics experiment showing the enrichment ratio (ER) (+UV enriched proteins/–UV enriched proteins) of all the proteins identified from the UV-dependent photocrosslinking by Probe 4 (50 μM, 30 min treatment), Probe 5 (50 μM, 4 h treatment), and Probe 6 (50 μM, 4 h treatment) when fed to RAW264.7 cells. Each data point represents the average ER for the respective protein from at least three biological replicates. The horizontal dotted line denotes an ER ≥ 4, and proteins having an ER above this threshold were considered as “hits” i.e. enriched by Probe 5 and Probe 6, and considered for subsequent analysis. For Probe 4, proteins above the threshold ER ≥ 3 were considered as “hits” i.e. enriched by Probe 4 and considered for subsequent analysis. (**G**) Venn diagram depicting shared and unique protein “hits” of Probe 4, Probe 5, and Probe 6 respectively. (**H**) Categorization of unique “hits” from Probe 5 after comparative analysis with Probe 4 and Probe 6 into lipid-binding proteins based on UniProt annotation^82^. For (**B,E**): The loading control for this gel (Coomassie staining) can be found in the *<u>Supplementary Information</u>*. For (**B-E**): This experiment was done three times with reproducible results each time. For (**F-H**): Complete details for all the “hits” from Probe 4 (129 proteins), Probe 5 (416 Proteins), and Probe 6 (545 Proteins) and their categorization can be found in *Supplementary Table 3*.

To negate these contributions, we employed two complementary bifunctional probe fragments as controls in all our assays in this strategy: (i) cholesterol diazirine alkyne (Ch-DA, Probe 6) to capture sterol-derived background interactors (**Figure 4A, Scheme 4**), and (ii) Probe 4 itself, to account for any fatty acid derived interactions (**Figure 4A**). Probe 6, which has a diazirine functional group embedded in the core cycle of cholesterol along with a propargyl group appended was synthesized in 4 steps from cholesterol (**Supplementary Scheme 1D, Supplementary Synthetic Note**). First, cholesterol was reacted with propargyl bromide under basic conditions to produce **4**. To install the diazirine functional group, we first need to prepare a ketone. We envisaged the conversion of the olefin of cholesterol to the ketone through a nitration-reduction sequence. Nitration of the olefin **4** gave the nitrovinyl derivative **5** in 61% yield. Reduction of the nitro group with zinc and acetic acid gave the ketone **6** in 43% yield, presumably through the enamine as the intermediate. The ketone was converted to the desired Probe 6 using the standard diazirination conditions in 27% yield (**Supplementary Scheme 1D**).

Previous studies have validated the uptake and functionality of Probe 4 in various mammalian cells^27,39^. Hence, to confirm its cellular uptake in RAW264.7 cells, we fed these cells Probe 4 (50 μM) over a time period from 30 min to 4h. In-gel fluorescence analysis confirmed optimal uptake of Probe 4 by RAW264.7 cells at all time points via only UV-irradiation dependent protein labeling readout at all these time points (**Supplementary Figure 7**). To additionally confirm probe uptake and its intactness in RAW264.7 cells, we also performed a fluorescence microscopy experiment under similar experimental conditions (50 μM, 30 min). Here, we found exclusive UV-dependent labeling throughout the cell, and no fluorescence in the absence of UV-irradiation (**Supplementary Figure 8**). These complementary experiments together confirmed that Probe 4 was taken up by RAW264.7 cells, and set a benchmark of concentration (50 μM) and treatment time (30 min) for all subsequent experiments using this strategy.

Next, to optimize conditions for protein labeling by Probe 5 in live cells using the PAL strategy, RAW264.7 macrophages were incubated with this lipid probe (50 μM) for time periods ranging from 30 min to 4 h (**Figure 4B**). In-gel fluorescence analysis revealed a time-dependent, linear increase in protein labeling, with maximal signal observed at 4 h. Labeling was strictly UV-dependent, as no fluorescence was detected in vehicle or no-UV controls. Probe uptake and chemical integrity were assessed by LC-MS/MS analysis, which demonstrated that intracellular Probe 5 co-eluted with an authentic synthesized Probe 5 standard (**Figure 4C**) and remained intact after 4 h, with no detectable hydrolysis to any probe fragments. Consistent with these findings, fluorescence microscopy following click conjugation to a fluorophore (Alexa Fluor 488) showed robust, UV-dependent labeling distributed throughout the cell, whereas no signal was observed in the absence of UV irradiation (**Figure 4D**).

We also performed similar validation studies for Probe 6 in RAW264.7 cells. When RAW264.7 cells were incubated with Probe 6 (50 μM) for 30 min to 4 h, in-gel fluorescence analysis showed a robust, UV-dependent protein labeling with maximal intensity at 4 h (**Supplementary Figure 9**), comparable to Probe 5. A LC-MS analysis confirmed cellular uptake and intactness of Probe 6, as evidenced by co-elution with an authentic chemically synthesized standard (**Supplementary Figure 10**), and fluorescence microscopy further demonstrated widespread, UV-dependent intracellular distribution of the probe 6 (50 μM) after 4 h (**Supplementary Figure 11**). A direct comparison of Probe 5 and Probe 6 from in-gel fluorescence labeling profiles revealed that Probe 6 produced a higher overall labeling intensity and a greater number of protein bands, consistent with higher cellular uptake of cholesterol relative to CEs by RAW264.7 cells (**Figure 4E**). Of note, some banding patterns were highly similar between Probe 5 and Probe 6, consistent with a potential overlap in sterol-driven interactions.

Having established optimal labeling conditions, we next performed the same quantitative proteomics analysis (SWATH-MS analysis^50–52^ for Probe 5 and Probe 6; and post-tryptic reductive dimethylation labeling^29^ for Probe 4) to identify proteins enriched specifically by Probe 5. Here, RAW264.7 cells were treated with each probe (50 μM probe, 4 h treatment for Probe 5 and Probe 6, and 30 min for Probe 4), followed by UV crosslinking, enrichment, and proteomic analysis in direct comparison to non-UV controls. Proteins were classified as probe-enriched if they were identified in at least two of three or four biological replicates, quantified with ≥ 3 peptides per replicate they were identified in, and exhibited an enrichment ratio (ER) ≥ 4 for Probe 5 and Probe 6 (or ≥ 3 for Probe 4) in all replicates (ER defined as abundance in UV vs non-UV samples, as mentioned earlier for Probe 5 and Probe 6, while for Probe 4 ER is heavy label to light label (H:L) for proteins in UV (Heavy) vs non-UV (Light) samples, where ER is capped at 10 in all cases).

Application of these stringent criteria yielded 416 proteins significantly enriched by Probe 5 upon UV treatment, nearly double the number identified using metabolic labeling probes or feeding Probe 3 to cells (**Figure 4F, Supplementary Table 3**). Using identical filtering criteria to those applied for Probe 5, identified 545 proteins were enriched by Probe 6 (**Figure 4F, Supplementary Table 3**), while only 129 proteins were identified as Probe 4 enriched upon UV crosslinking **Figure 4F, Supplementary Table 3**). A comparative analysis of the proteins enriched by all the probes (Probe 4, Probe 5 and Probe 6) revealed that only 224 proteins uniquely enriched by Probe 5, while, 12 proteins common to all three probes, and 158 and 22 proteins shared with Probe 6 and Probe 4, respectively (**Figure 4G, Supplementary Table 3**). Functional annotation of the 224 and 356 proteins uniquely enriched by Probe 5 and Probe 6 respectively, using the PANTHER database^53,54^ revealed a diverse distribution of protein classes, including adaptor, transporter, defence, and storage proteins, spanning multiple biological pathways and physiological functions (**Supplementary Figures 12-13, Supplementary Table 3**). These results define a refined subset of CE-specific protein interactors and highlight the utility of Probe 5, in combination with appropriate control probe fragments (Probe 4 and Probe 6), for dissecting lipid–protein interaction landscapes with high specificity. Notably, amongst the 224 proteins exclusively enriched by Probe 5, several known lipid-binding and sterol-interacting proteins were present, supporting the conclusion that Probe 5 effectively mimics endogenous CEs in mammalian cells (**Figure 4H, Supplementary Table 3**).

### Comparing the three strategies

To comprehensively map the cellular CE interactome, we performed an integrated comparative analysis of protein “hits” identified across the three orthogonal strategies described in this study. Collectively, these approaches yielded 495 putative CE-interacting proteins (**Figure 5A, Supplementary Table 4**). Notably, each strategy contributed a substantial and comparable subset of uniquely enriched proteins (**Figure 5A**), underscoring the complementary nature of these methodologies. Specifically, 101 proteins were uniquely identified through the metabolic labeling strategy, 95 proteins were uniquely enriched using Probe 3 in the functionalized probe approach, and 87 proteins were uniquely captured using Probe 5 in the fragment-assisted functionalized probe strategy (**Figure 5A**). In contrast, only 53 proteins were commonly enriched across all three platforms (**Figure 5A**), highlighting the limited overlap between individual methods and emphasizing the necessity for multi-modal chemical proteomic strategies to achieve a more complete and unbiased mapping of the CE–protein interaction landscape in mammalian systems.

**Figure 5.**
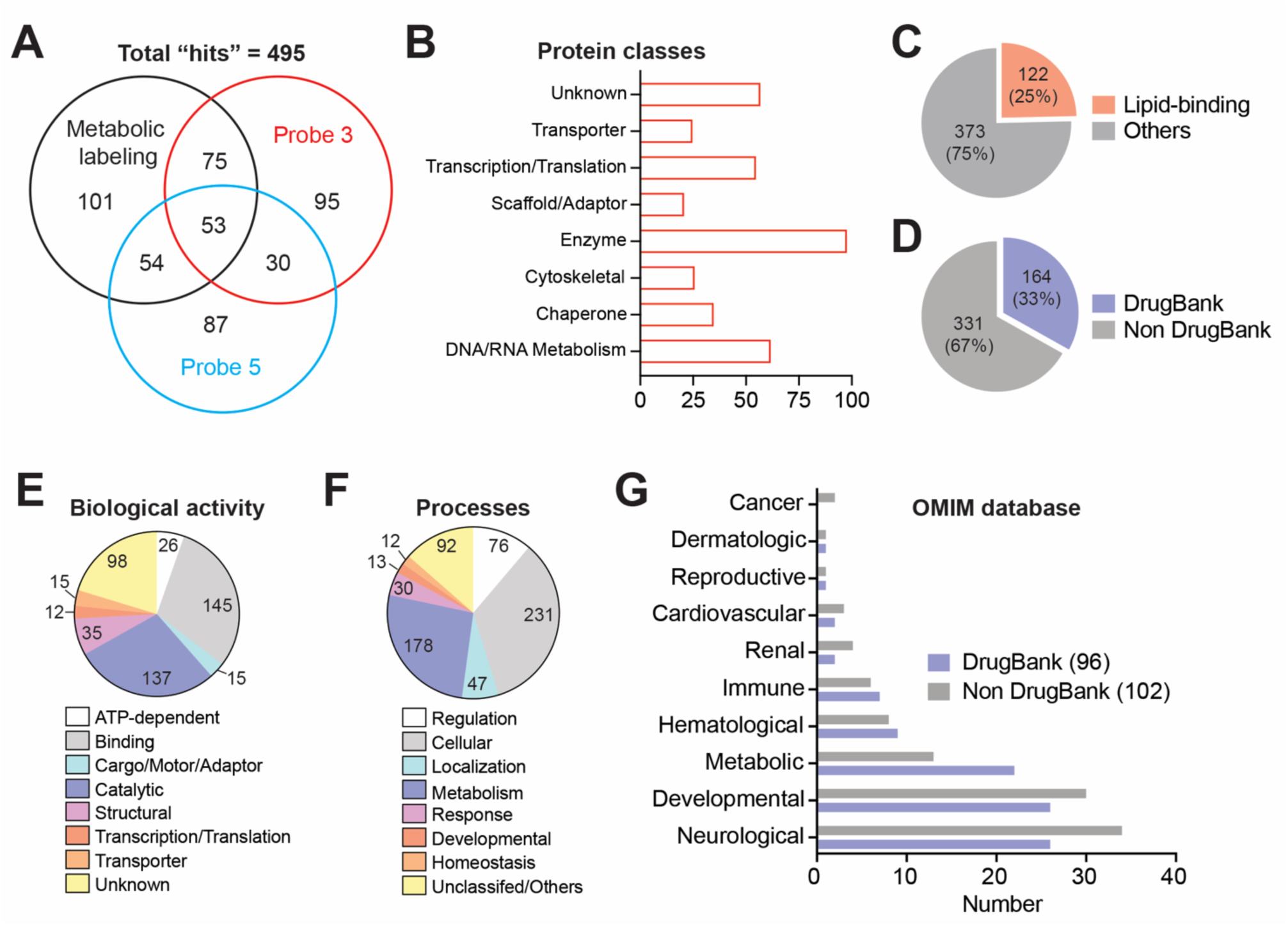
A comparative analysis of protein “hits” identified across the three orthogonal strategies. (**A**) Venn diagram depicting common, overlapping and unique protein “hits” (495 total proteins) from the metabolic labeling strategy, Probe 3 (functionalized probe strategy), and Probe 5 (fragment assisted functionalized probe strategy) respectively. (**B-D**) Categorization of total “hits” (495 proteins in total) obtained from all three orthogonal strategies into: (**B**) protein classes based on the Panther database classification^53,54^, (**C**) lipid-binding proteins based on the Uniprot annotation^82^, and (**D**) known drug targets based on the distribution in DrugBank database^55^. (**E-F**) Mapping of total “hits” (495 proteins in total) using the Panther database classification into: (**E**) known biological activities; and (**F**) physiological processes they are involved in. (**G**) CE-protein targets from the analysis performed in (**D**) that have been genetically linked to human diseases based on searches of the OMIM database^56^. For (**A-G**): Complete details for all the “hits” (495 proteins in total) from all three strategies and their categorization can be found in *Supplementary Table 4*.

Functional annotation of these 495 protein hits using the PANTHER classification^53,54^ again revealed broad representation across diverse protein classes (**Figure 5B, Supplementary Table 4**), including enzymes, cytoskeletal proteins, transporters, and adaptor or scaffolding proteins.

Importantly, several established lipid-interacting and sterol-associated protein families were prominently enriched, including lipid transporters, transmembrane proteins, lipid storage proteins, and canonical sterol-binding proteins, which together accounted for ∼25% (122 of 495 proteins) of the total dataset (**Figure 5C**). To further evaluate the pharmacological relevance of the CE interactome, we mapped these proteins to the DrugBank database^55^. This analysis revealed a marked enrichment of known drug targets within the CE-interacting proteome, with ∼33% (164 proteins) corresponding to annotated DrugBank entries (**Figure 5D, Supplementary Table 4**), compared to an estimated ∼18% representation of druggable targets in the overall human proteome. The remaining 331 proteins, which lacked DrugBank annotation, constitute a substantial pool of previously underexplored or non-canonical targets, and may represent a valuable resource for the identification of novel ligandable proteins and future therapeutic intervention points.

We next examined the functional distribution of the CE-associated proteome in terms of molecular function and biological processes using PANTHER ontology mapping^53,54^. At the level of biological activity, CE-interacting proteins were predominantly associated with binding functions, catalytic activity, structural roles, and transporter activity (**Figure 5E, Supplementary Table 4**). In parallel, physiological process analysis revealed extensive involvement in pathways central to lipid homeostasis, metabolic regulation, intracellular signaling, and developmental programs (**Figure 5F, Supplementary Table 4**) with diverse subcellular localisations (**Supplementary Figure 14**), collectively suggesting that CE interactions are embedded within core cellular regulatory networks rather than restricted to canonical lipid-handling pathways alone.

Finally, to explore potential disease associations of the CE interactome, we interrogated both DrugBank-annotated and non-annotated protein subsets against the OMIM (Online Mendelian Inheritance in Man) database^56^. Despite the well-established involvement of CEs in pathological conditions such as atherosclerosis, neurodegeneration, and inflammation, their direct protein interaction landscape has remained unexplored in a disease context. OMIM analysis of the DrugBank associated subset identified 96 proteins with documented links to human diseases, with strong representation in neurological, developmental, and metabolic disorders (**Figure 5G, Supplementary Table 4**). Strikingly, a parallel analysis of the non-DrugBank subset revealed 102 additional disease-associated proteins spanning neurological, developmental, metabolic, cardiovascular, and renal disease categories (**Figure 5G, Supplementary Table 4**). Taken together, these results provide the first systems-level view of the CE-interacting proteome in cells, revealing both its extensive functional diversity and its significant convergence with disease-relevant and pharmacologically actionable protein networks.

## DISCUSSION

CEs have traditionally been viewed as inert storage forms of excess cholesterol, largely confined to lipid droplets and circulating lipoproteins with limited regulatory function^5,6,21^. In contrast, free cholesterol and phospholipids have received far greater attention as modulators of membrane organization and signaling^5,6^. The present study revisits this long-standing perception by systematically interrogating CE-protein interactions in mammalian cells and revealing that CEs engage a wide and functionally heterogeneous proteome^57–59^. The collective datasets generated here demonstrate that CE associations extend well beyond canonical lipid transport or storage pathways and intersect with proteins involved in metabolism, membrane trafficking, structural organization, and intracellular signaling. Such breadth suggests that CEs may influence cellular physiology through distributed and context-dependent interactions rather than through a small set of dedicated high-affinity receptors, thereby positioning this lipid class as a more active participant in cellular regulatory networks than previously appreciated.

A defining feature of this study is the deliberate integration of three orthogonal PAL based chemical proteomic strategies (**Figure 1**), metabolic assembly of functionalized CEs, direct feeding of tailored bifunctional CE probes, and fragment-assisted functional probe analysis, to construct a composite interaction atlas for the CE-protein interactome. Each strategy samples a distinct dimension of lipid–protein engagement. Metabolic labeling leverages endogenous cellular enzymatic machinery to assemble the probe within native lipid biosynthetic circuits, thereby approximating physiological routing and intracellular distribution. Direct probe feeding offers chemical precision and experimental control, enabling defined delivery of structurally characterized CE analogs. The fragment-assisted strategy, in turn, provides a subtractive framework to distinguish interactions unique to the intact CE scaffold from those attributable to sterol or fatty acyl fragments alone. The limited overlap observed across these modalities underscores that CE-protein interactions are strongly influenced by lipid origin, compartmentalization, and perhaps even, local concentration gradients in cells. Rather than reflecting methodological inconsistency, this divergence highlights the intrinsic spatial and metabolic heterogeneity of lipid biology, where organelle localization and lipid flux dictate which subsets of proteins are accessible to a given lipid species at a given time in mammalian cells.

The fragment-guided experiments provide additional insight into lipid recognition principles by demonstrating that a subset of proteins is uniquely enriched when the intact esterified architecture is preserved in CEs. This observation implies that the combined sterol-acyl topology, and not merely its individual components, confers distinct physicochemical features recognizable by cellular proteins. Such features may arise from altered membrane insertion depth, conformational flexibility, or hydrophobic surface presentation imparted by the ester linkage. These findings align with an emerging view in lipid chemical biology that composite lipid structures and its overall three-dimensional organization, rather than isolated functional groups, frequently govern molecular recognition^57–59^. In this framework, CEs may modulate protein function not only through direct binding but also through effects on membrane microenvironments, lipid droplet interfaces, or transient contact zones that influence protein localization and activity.

Functionally, the CE-associated proteome uncovered here spans enzymes, transporters, adaptor and scaffolding proteins, cytoskeletal components, and transmembrane factors, reinforcing the idea that CE engagement intersects with core cellular regulatory circuits. The enrichment of established lipid and sterol-binding protein families provides internal validation of probe fidelity, while the concurrent identification of numerous previously unannotated or non-canonical targets expands the conceptual boundaries of CE biology. Importantly, mapping these proteins to pharmacological and disease-associated databases reveals substantial convergence with known drug targets as well as a large subset of understudied proteins lacking prior ligand annotation^55,56^. In light of accumulating evidence linking CE dysregulation to neurodegenerative disorders^60–68^, oncogenesis^13,19,20,69–71^, and cardiovascular disease^12,68,71–74^, this interaction atlas provides a foundational resource for connecting CE metabolism to disease-relevant protein networks and for identifying potentially ligandable nodes within these pathways.

Despite the breadth and internal consistency of these datasets, several limitations merit consideration. First, PAL captures proximity-dependent events and therefore encompasses indirect associations, membrane co-residency, and transient encounters in addition to direct binding^42,75^.

The interactome described here should thus be interpreted as a landscape of potential engagement rather than a definitive catalogue of high-affinity ligands. Second, although probe design was guided by structural fidelity and supported by extensive controls, any chemical modification of a lipid, whether introduced synthetically or via metabolic incorporation, may subtly influence membrane partitioning, intracellular trafficking, or protein affinity. Third, quantitative proteomic thresholds, while intentionally stringent, inherently favor detection of abundant or readily ionizable proteins and may underrepresent low-abundance, highly hydrophobic, or transiently expressed targets^42^. A further limitation is intrinsic to the metabolic labeling paradigm itself: this strategy is contingent on the presence and activity of the requisite biosynthetic enzymes within a given cell type. Cells that lack, or express low levels of, enzymes such as ACAT/SOAT are unlikely to efficiently assemble the CE probe *in situ*, thereby restricting the applicability of metabolic labeling across diverse biological systems and necessitating complementary exogenous probe-based strategies. Finally, the majority of experiments were performed in a single macrophage-derived cell line under defined culture conditions, and CE-protein interactions are expected to vary across tissues, developmental stages, and metabolic or inflammatory states as well.

These considerations delineate several productive avenues for future investigation. Extending this chemical proteomic framework to additional cell types, primary cells, and *in vivo* models will be essential for defining tissue-specific and disease-contextual CE interaction networks. Orthogonal validation approaches, including competitive displacement with native lipids, mutagenesis of candidate interaction sites, structural biology, and quantitative biophysical assays, will further help distinguish direct ligands from indirect associations and refine mechanistic understanding of CE-binding proteins. Spatially restricted or organelle-targeted probe designs could enable higher-resolution mapping of CE interactions at lipid droplets, endoplasmic reticulum membranes, or plasma membrane microdomains, while temporal profiling under metabolic perturbations or pharmacological inhibition of lipid enzymes may illuminate dynamic CE-dependent signaling axes. Integration with computational docking and motif discovery efforts may further reveal recurring structural determinants of CE recognition analogous to established sterol-binding motifs. Collectively, the multi-modal strategy and interaction resource presented here reposition CEs from passive storage entities to active components of cellular protein networks and establish a versatile conceptual and methodological platform for future exploration of this understudied yet disease-relevant lipid class.

## CONCLUSIONS

In conclusion, we establish CEs as functionally engaged components of the cellular proteome rather than inert depots of excess cholesterol. By integrating three orthogonal chemical proteomic strategies namely metabolic probe assembly, direct delivery of structurally defined CE analogs, and fragment-assisted subtraction, we generated a systems-level and internally validated atlas of 495 CE-associated proteins in mammalian cells. The limited overlap across platforms underscores the context dependence of lipid–protein engagement, shaped by biosynthetic origin, intracellular routing, and scaffold topology. Importantly, convergence on established sterol-binding families, together with enrichment of drug-annotated and disease-linked proteins, supports both probe fidelity and translational relevance. Beyond expanding the molecular landscape of CE biology, this work provides a generalizable blueprint for resolving complex lipid interactomes.

Collectively, these findings reposition CEs within mainstream chemical biology and motivate future structure-guided and in vivo investigations into CE-mediated regulatory mechanisms and therapeutic opportunities.

## MATERIALS AND METHODS

### Materials

Unless otherwise mentioned: all chemicals, buffers, and reagents were purchased from Sigma-Aldrich (now Merck); all tissue culture media and supplies were purchased from HiMedia; and all mass spectrometry solvents used were LC-MS grade and were purchased from JT Baker.

### Mammalian cell culture and probe treatments

All mammalian cell lines used in this study (RAW264.7 and Neuro2A) were obtained from ATCC and maintained according to the supplier’s recommended protocols. Cells were cultured in Dulbecco’s Modified Eagle’s Medium (DMEM) supplemented with 10% (v/v) heat-inactivated fetal bovine serum (FBS) and 1% (v/v) penicillin– streptomycin solution (MP Biomedicals). Cultures were maintained at 37 °C in a humidified incubator with 5% (v/v) CO2. Cell morphology and confluency were routinely monitored using phase-contrast microscopy. As a quality control measure, all cell lines were periodically screened for mycoplasma contamination using 4′,6-diamidino-2-phenylindole (DAPI) staining followed by fluorescence microscopy. Only mycoplasma-free cultures were used for downstream experiments. For probe-labeling experiments, cells were seeded in 6 cm or 10 cm tissue culture dishes at an initial confluency of 50–60% and allowed to adhere and recover for 16 to 24 h prior to treatment.

For live-cell labeling experiments, growth media was aspirated and adherent cells were gently washed twice with sterile Dulbecco’s phosphate-buffered saline (DPBS) to remove residual serum components. Cells were then incubated in serum-free DMEM for the duration of probe treatment. The lipid probes were prepared as 10 mM stock solutions in molecular grade DMSO (Sigma-Aldrich, Catalog No: D8418) and diluted to the desired working concentrations directly in pre-warmed culture medium under low ambient light conditions to minimize photodegradation. The final DMSO concentration in all treatments was maintained below 1% (v/v). For gel-based profiling, proteomics, and LC–MS/MS–based lipidomics experiments, cells were incubated with probes at 37 °C for the indicated durations (specified in corresponding figure legends) in the absence of light. Following incubation, cells were washed with ice-cold sterile DPBS (2x) to remove excess probe. Photocrosslinking was performed by irradiating the adherent cells in cold DPBS using a 365 nm UV source in a CL-1000L UVP crosslinker (Analytik Jena) for 10 min. Cells were subsequently harvested by scraping, pelleted by centrifugation, flash-frozen in liquid nitrogen, and stored at −80 °C until further processing.

### In-gel fluorescence analysis

In-gel fluorescence experiments were conducted with minor modifications to previously reported protocols^29^. Briefly, probe-labeled cell pellets were thawed on ice, resuspended in cold DPBS, and lysed by probe sonication using short pulses while maintaining samples on ice to prevent overheating. Total protein concentrations were determined using the Bradford assay with bovine serum albumin as a standard. For click chemistry conjugation, 100 µL of clarified proteome (2 mg/mL total protein) was reacted with a freshly prepared click reagent mixture (11 µL) consisting of Tris-(benzyl-triazolylmethyl)-amine (TBTA, 6 µL of 1.7 mM stock in 4:1 DMSO: *tert*-butanol), copper sulfate (CuSO₄, 2 µL of 50 mM stock), Tris- (2-carboxyethyl)-phosphine (TCEP, 2 µL of 50 mM stock), and rhodamine azide (Sigma-Aldrich, Catalog No: 760765) (1 µL of 10 mM stock). Reactions were incubated for 1 h at 25 °C with gentle agitation. The reactions were quenched by the addition of 4xSDS–PAGE loading buffer without prior boiling to preserve fluorescence signal. Proteins were resolved on 10% SDS–polyacrylamide gels under standard electrophoretic conditions. Fluorescently labeled proteins were visualized using an iBright 1500 imaging system (Invitrogen). Following fluorescence acquisition, gels were stained with Coomassie Brilliant Blue-R to verify equal protein loading and imaged using a Syngene Chemi-XRQ gel documentation system.

### Imaging lipid probe in mammalian cells

Microscopy slides were prepared based on previously reported methods with minor modifications^27,76,77^. After UV irradiation, adherent cells were washed with sterile DPBS (2x) and fixed with ice-cold methanol for 10 min at −20 °C. Fixed cells were sequentially washed with isopropanol for 2 min and then treated with a chloroform/methanol/acetic acid mixture (10:55:0.75, v/v/v) for 2 min to remove unbound probe molecules. Cells were subsequently washed with sterile DPBS (3x) for 2 min each. The click conjugation was performed directly on coverslips using a 100 µL click reaction mixture containing TCEP (1 mM), TBTA (100 µM), CuSO₄ (1 mM), and Alexa Fluor 488 azide (Sigma-Aldrich, Catalog No: 760765) (2 µM). Reactions were carried out in a humidified chamber at room temperature for 1 h, protected from light. After completion, cells were washed thoroughly with DPBS and counterstained with 1x DAPI solution to visualize nuclei. Coverslips were mounted onto glass slides using Fluoromount mounting medium (Sigma–Aldrich, Catalog No: F4680) and allowed to cure before imaging. Confocal fluorescence images were acquired using a Zeiss LSM 770 confocal microscope equipped with a 63x oil immersion objective. Image processing and quantitative analyses were performed using Fiji (ImageJ) software with identical acquisition parameters applied across all samples^78,79^.

### LC-MS/MS based lipid probe measurement in mammalian cells

Intracellular lipid probe levels were quantified using a modified Folch extraction procedure^10,80^ without UV crosslinking. Frozen cell pellets were thawed on ice, resuspended in 1 mL ice-cold DPBS, and transferred to glass extraction vials. A volume of 3 mL chloroform:methanol (2:1, v/v) containing 1 nmol of C17:0 cholesteryl ester internal standard (Sigma–Aldrich, Catalog No: 700186M) was added to each sample. Mixtures were vortexed vigorously and centrifuged at 1500g for 15 min at room temperature to achieve phase separation. The lower organic phase was carefully collected without disturbing the aqueous layer and transferred to a fresh glass vial. Solvents were evaporated under a gentle stream of nitrogen gas. To remove residual aqueous contaminants, dried extracts were reconstituted in 1 mL chloroform, vortexed, and dried again under nitrogen. Final lipid extracts were dissolved in 200 µL chloroform:methanol (2:1, v/v), and 10 µL aliquots were injected for LC– MS/MS analysis. Mass spectrometric analyses were performed on an Agilent 6545 Quadrupole Time-of-Flight (QTOF) mass spectrometer equipped with an electrospray ionization source. Instrument parameters, chromatographic conditions, and acquisition settings were consistent with previously validated laboratory protocols to ensure reproducibility and sensitivity^10^.

### Proteomics sample preparation and SWATH-MS analysis

Proteomic sample preparation was performed following established protocols with minor optimizations^29^. For each experiment, 2 mg of probe-labeled proteome in 1 mL sterile DPBS was subjected to click chemistry using biotin azide as the reporter tag. The click reaction mixture (110 µL in total per sample) consisted of TBTA (100 µM), CuSO_4_ (1 mM), TCEP (1 mM), and biotin azide (Sigma-Aldrich, Catalog No: 762024) (100 µM). Samples were incubated at 25 °C for 1 h with continuous mixing. Following conjugation, proteins were denatured, reduced, and alkylated using iodoacetamide. Proteolytic digestion was carried out overnight at 37 °C using sequencing-grade trypsin (Promega, Catalog No: V5111) at an enzyme-to-substrate ratio of 1:50 (w/w). Resulting peptides were desalted using C18 StageTips^81^ and dried under vacuum prior to LC–MS/MS analysis.

All proteomic LC–MS/MS analyses were conducted on a Sciex TripleTOF 6600 mass spectrometer coupled to an Eksigent nanoLC 425 system. Peptides were initially loaded onto a C18 trap column and subsequently separated on a C18 analytical column (15 cm x 75 µm internal diameter) using a linear acetonitrile gradient at a constant flow rate of 300 nL/min. Solvent A consisted of water with 0.1% formic acid, and solvent B consisted of acetonitrile with 0.1% formic acid. A typical gradient program included 5% solvent B for 1 min, a linear increase to 30% solvent B over 330 min, 90% solvent B for 20 min, followed by re-equilibration at 5% solvent B for 10 min. For ion library generation, data were acquired in information-dependent acquisition mode across an m/z range of 200–2,000. Each full MS scan was followed by MS/MS acquisition of the 15 most intense precursor ions. Dynamic exclusion parameters were set to a repeat count of 2 and an exclusion duration of 6 sec. Peptide identification was performed using ProteinPilot (v2.0.1, Sciex) with the Paragon and ProGroup algorithms against the *Mus musculus* RefSeq protein database (Release 109). Carbamidomethylation of cysteine was set as a fixed modification, whereas methionine oxidation and N-terminal acetylation were considered variable modifications. Precursor and fragment mass tolerances were set to 20 ppm and 50 ppm, respectively. A decoy database strategy was employed to control false discovery rates (FDR), and only identifications with FDR < 1% were retained.

For label-free quantitative analysis (for all the probes except Probe 4, PA-DA), Sequential Window Acquisition of All Theoretical Fragment Ion Spectra (SWATH–MS)^50–52^ was performed in data-independent acquisition mode across an m/z range of 400–1,500. Variable isolation windows were defined as 5 Da (400–600 m/z), 15 Da (600–800 m/z), and 50 Da (800–1,200 m/z). Protein identification and quantification were carried out using PeakView software (v2.2, Sciex).

Quantitative criteria included a minimum of three peptides per protein, six transitions per peptide, peptide confidence ≥ 95%, and FDR ≤ 1%. Protein peak areas were used for enrichment analysis, and only proteins consistently identified in at least two biological replicates were considered for downstream interpretation. For comparative analyses between UV-treated and non-UV controls, a maximal enrichment ratio cutoff of 10 was applied to minimize outlier-driven bias.

For quantitative analysis of Probe 4-labelled proteins, established reductive demethylation (ReDiMe) peptide labeling strategy^29^ was used where the tryptic peptides obtained from the UV-crosslinking group were labeled with heavy formaldehyde (CD2O) (Cambridge Isotope Laboratories Inc.; Catalog No: DLM-805-20), while those from the non-UV treated controls were labeled with light formaldehyde (CH2O) (Sigma-Aldrich; Catalog No: 252549). Post ReDiMe labeling, the heavy and light labeled peptides were mixed together and desalted using the StageTip protocol^81^. Data acquisition and peptide identification was performed using the same acquisition and analysis parameters described for the ion library generation. The ReDiMe algorithm was selected within ProteinPilot software for quantification of identified proteins and the enrichment ratio (heavy label to light label) was used for enrichment-based analysis. Proteins consistently identified in at least two biological replicates were only considered for downstream interpretation. For comparative analyses between UV-treated and non-UV controls, a maximal enrichment ratio cutoff of 10 was applied to minimize outlier-driven bias.

## Supporting information

Synthetic note

## SUPPLEMENTARY INFORMATION

The supplementary files include: (a) Supplementary synthetic note (PDF); (b) Supplementary Information containing Supplementary Schemes 1 – 3, Supplementary Figures 1 – 14 and Loading control gels; and (c) Supplementary Tables 1 – 4 (XLSX).

## AUTHOR CONTRIBUTIONS

A.C. performed all the biological studies with assistance from A.K., C. K. and M.D.; K.S. developed the methodology and synthesized all the new probes (probes 1, 2, 3, 5, 6) reported in this study.

P.T. provided PA-DA (probe 4) that was used in this study. H.C. and S.S.K. conceived, supervised and acquired funding for this project. A.C., K.S., H.C., and S.S.K. wrote the manuscript with inputs from all authors.

## ACKNOWLEDGEMENTS

We thank Saddam Shekh for maintenance of the biological mass spectrometry facility at IISER Pune, and for technical assistance. We also thank the IISER Pune microscopy facility for providing access to and technical expertise for microscopes needed for the lipid probe imaging experiments. Members of the S.S.K. and H.C. labs at IISER Pune are thanked for providing critical comments and inputs throughout the course of this study.

## FUNDING

We acknowledge generous financial support from the Anusudhan National Research Foundation (ANRF), Government of India (Grants: SwarnaJayanti Fellowship SB/SJF/2021-22/01 to S.S.K.), and an EMBO Young Investigator Award (to S.S.K.). A.C. and A.K. are supported by graduate student fellowships from IISER Pune.

## DATA AVAILABILITY

All data that supports the findings of this study are available in this paper and its Supplementary Information or are available with the corresponding author (S.S.K.) upon reasonable request.

All the raw proteomics data have been deposited with the ProteomeXchange Consortium via the PRIDE partner repository with the identifiers: PXD072445 (SWATH library generation), PXD072490 (metabolic labeling), PXD072501 (Probe 3), PXD072423 (Probe 4), PXD072454 (Probe 5), and PXD072459 (Probe 6).

## Notes

### Competing Interest Statement

The authors have declared no competing interest.

