## Supplementary material for "Orthogonal Chemical Proteomic Strategies Reveal the Cholesteryl Ester Interactome in Mammalian Cells": Synthetic note

**SUPPLEMENTARY SYNTHETIC NOTE**

#### General information:

Unless otherwise specified, all reagents, starting materials, and dry solvents were purchased from commercial suppliers and used as received without further purification. Glassware items were oven-dried prior to use. All reactions were conducted under a nitrogen atmosphere. Analytical thin-layer chromatography (TLC) was performed using Silica Gel 60 F254 pre-coated plates (0.25 mm thickness, Merck), and visualized with staining using phosphomolybdic acid (PMA) solution. All work up and purification were carried out with reagent grade solvents in air. Column chromatography was performed on Rankem silica gel (100-200 mesh) using reagent grade solvents. $^{1}$H (400 MHz) and $^{13}$C (100 MHz) nuclear magnetic resonance (NMR) were recorded on BRUKER 400 spectrometer using tetramethylsilane (TMS, $\delta_H$ = 0.00 ppm, $\delta_C$ = 0.0 ppm) as an internal standard or residual solvent (Chloroform, $\delta_H$ = 7.26 ppm, $\delta_C$ = 77.2 ppm) signals as reference. Chemical shifts ($\delta$) are reported in parts per million (ppm) and coupling constants ($J$) in Hz. The following notations are used to indicate the multiplicity of the signals: s (singlet), broad singlet (bs), d (doublet), dd (doublet of doublet), t (triplet), td (triplet of doublet), q (quartet), qd (quartet of doublet), quint (quintet) and m (multiplet). Liquid Chromatography-High Resolution Mass Spectra (LC-HRMS) were obtained using a Agilent 6545 Quadrapole Time of Flight (QTOF) using established protocols for cholesterol, cholesteryl esters, and fatty acids[1,2]. Infrared (IR) spectra were recorded using BRUKER ALPHA FT-IR spectrometer.

#### Synthesis of cholesterol alkyne, Probe 1

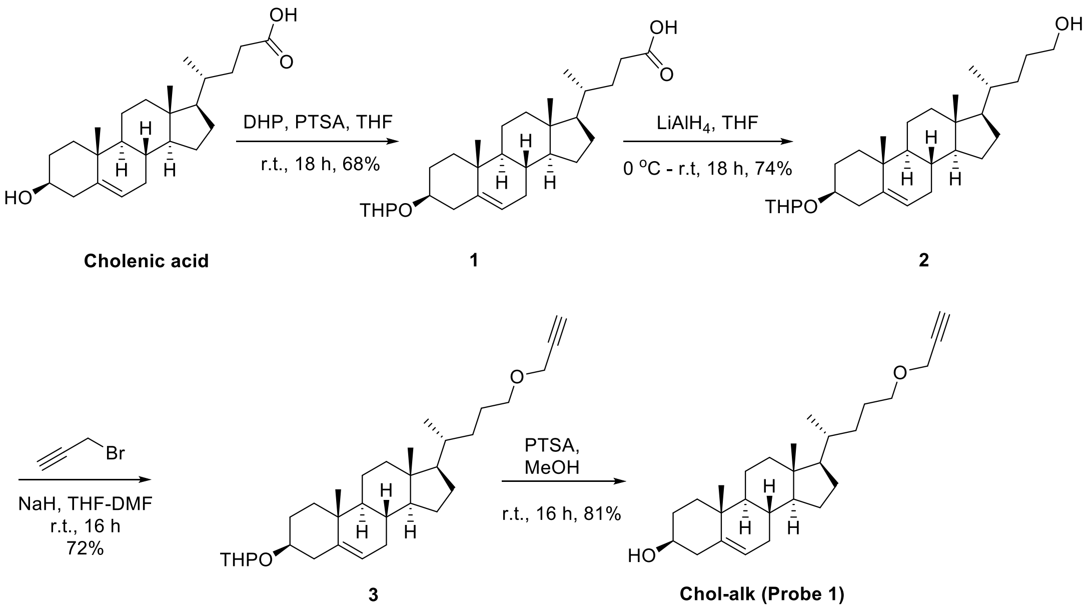

**(4*R*)-4-((3*S*,8*S*,9*S*,10*R*,13*R*,14*S*,17*R*)-10,13-Dimethyl-3-((tetrahydro-2*H*-pyran-2-yl)oxy)-2,3,4,7,8,9,10,11,12,13,14,15,16,17-tetradecahydro-1*H*-cyclopenta[a]phenanthren-17-yl)pentanoic acid, 1**

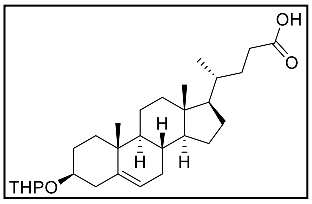


Compound **1** was synthesized using a reported protocol[3]. Briefly, to a solution of cholenic acid (1 g, 2.67 mmol) in dry THF (100 mL) under nitrogen, 3,4-dihydro-2*H*-pyran (DHP, 1.46 mL, 16.02 mmol) and *para*-toluene sulfonic acid monohydrate (PTSA, 0.101 g, 0.53 mmol) were added. The reaction mixture was stirred at room temperature for 18 h and then progress of reaction was monitored using TLC. The reaction mixture was diluted with 300 mL water, and the solution was extracted using dichloromethane (3x50 mL). The combined organic layer was washed with brine, dried over anhydrous sodium sulphate and then filtered. The filtrate was concentrated and the residue was purified using column chromatography, silica gel 100-200 mesh. The product eluted at 10% ethyl acetate-pet ether and upon concentration and drying the desired product was obtained as white solid (1.05 g, 86%): $^1$H NMR (400 MHz, $CDCl_3$): $\delta$ 5.36-5.33 (m, 1H), 4.72-4.71 (m, 1H), 3.94-3.90 (m, 1H), 3.56-3.47 (m, 2H), 2.43-2.15 (m, 4H), 2.01-1.95 (m, 2H), 1.90-1.56 (m, 10H), 1.53-1.03 (m, 16H), 1.00 (s, 3H), 0.93 (d, $J$ = 6.4 Hz, 3H), 0.68 (s, 3H).

**(4*R*)-4-((3*S*,8*S*,9*S*,10*R*,13*R*,14*S*,17*R*)-10,13-Dimethyl-3-((tetrahydro-2*H*-pyran-2-yl)oxy)-2,3,4,7,8,9,10,11,12,13,14,15,16,17-tetradecahydro-1*H*-cyclopenta[a]phenanthren-17-yl)pentan-1-ol, 2**

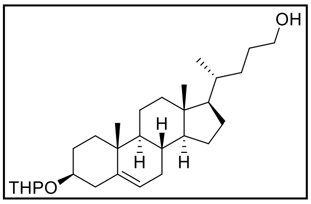


Compound **2** was synthesized using a reported protocol[3]. Briefly, to an ice-cold solution of lithium aluminium hydride ($LiAlH_4$, 0.52 g, 13.69 mmol) in dry THF (60 mL) under nitrogen, the solution of compound **1** (1.047 g, 2.28 mmol) in dry THF (40 mL) was dropwise added. The reaction mixture was warmed to room temperature and stirred for 18 h. Completion of reaction was monitored using TLC and the reaction was then quenched with saturated ammonium chloride solution. The aqueous solution was then extracted with dichloromethane (3x50 mL). The combined organic layer was washed with brine, dried over anhydrous sodium sulphate and then filtered. The filtrate was concentrated and the residue was purified using column chromatography, silica gel 100-200 mesh. The product eluted at 10% ethyl acetate-pet ether and upon concentration and drying the desired product was obtained as white solid (0.938 g, 92%): $^1$H NMR (400 MHz, $CDCl_3$): $\delta$ 5.34 (t, $J$ = 6.0 Hz, 1H), 4.71-4.70 (m, 1H), 3.93-3.89 (m, 1H), 3.63-3.46 (m, 4H), 2.36-2.16 (m, 2H), 2.01-1.62 (m, 9H), 1.57-1.02 (m, 21H), 1.00 (s, 3H), 0.93 (d, $J$ = 6.5 Hz, 3H), 0.67 (s, 3H).

**2-(((3*S*,8*S*,9*S*,10*R*,13*R*,14*S*,17*R*)-10,13-Dimethyl-17-((*R*)-5-(prop-2-yn-1-yloxy) pentan-2-yl)-2,3,4,7,8,9,10,11,12,13,14,15,16,17-tetradecahydro-1*H*-cyclopenta[a]phenanthren-3-yl)oxy) tetrahydro-2*H*-pyran, 3**

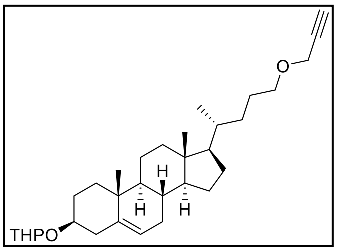


A solution of sodium hydride (NaH, 0.178 g, 4.45 mmol, 60% dispersion in oil) in dry THF (8 mL) and DMF (2 mL) was stirred for 15 min under nitrogen atmosphere. Compound **2** (0.330 g, 0.742 mmol) was added to the solution and stirred at room temperature for 30 min. To this solution, propargyl bromide (0.25 mL, 2.22 mmol) was added dropwise. The reaction mixture was then stirred at room temperature for 16 h. The reaction was quenched using water (100 mL) and extracted using ethyl acetate (3x15 mL). The combined organic layer was again washed with water, brine and dried over anhydrous sodium sulphate and then filtered. The filtrate was concentrated and the residue was purified using column chromatography, silica gel 100-200 mesh. The product eluted at 5-8% ethyl acetate-pet ether and upon concentration and drying the desired product was obtained as white solid (0.26 g, 72%): $^1$H NMR (400 MHz, $CDCl_3$): $\delta$ 5.34 (t, $J$ = 5.8 Hz, 1H), 4.71-4.70 (m, 1H), 4.13 (d, $J$ = 2.3 Hz, 2H), 3.93-3.89 (m, 1H), 3.56-3.44 (m, 4H), 2.41 (t, $J$ = 2.3 Hz, 1H), 2.36-2.17 (m, 2H), 2.04-1.94 (m, 2H), 1.90-1.78 (m, 4H), 1.74-1.28 (m, 22H), 1.19-1.04 (m, 6H), 1.00 (s, 3H), 0.92 (d, $J$ = 6.5 Hz, 3H), 0.67 (s, 3H) The presence of grease from solvent was noted; $^{13}$C NMR (100 MHz, $CDCl_3$): $\delta$ 141.2, 141.1, 121.7, 121.6, 97.1, 97.0, 80.2, 76.2, 74.2, 71.0, 63.1, 63.0, 58.1, 56.9, 56.1, 50.3, 42.5, 40.4, 39.9, 38.9, 37.6, 37.3, 36.9, 35.7, 32.3, 32.1, 32.0, 31.4, 29.8, 28.4, 28.1, 26.2, 25.6, 24.4, 21.2, 21.2, 20.2, 20.2, 19.5, 18.8, 12.0; FT-IR ($\nu_{max}$, $cm^{-1}$): 3304 (alkyne C-H), 1105 (ether C-O-C); LC-HRMS for $C_{32}H_{50}O_3$ $[M+NH_4]^+$: Calculated: 500.4104, Found: 500.4062.

**(3*S*,8*S*,9*S*,10*R*,13*R*,14*S*,17*R*)-10,13-Dimethyl-17-((*R*)-5-(prop-2-yn-1-yloxy) pentan-2-yl)-2,3,4,7,8,9,10,11,12,13,14,15,16,17-tetradecahydro-1*H*-cyclopenta[*a*]phenanthren-3-ol, Chol-Alk (Probe 1)**

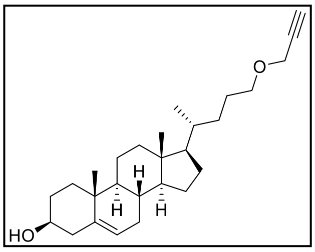


To a solution of compound **3** (0.3 g, 0.62 mmol) in THF (8 mL) and methanol (8 mL) at room temperature, PTSA (0.024 g, 0.12 mmol) was added, and the reaction mixture was stirred for 2 h. Upon completion of reaction (TLC monitored), the reaction mixture was concentrated in vacuo and then dissolved in diethyl ether (50 ml). The organic layer was washed with saturated sodium bicarbonate solution, brine and dried over anhydrous sodium sulphate and then filtered. The filtrate was concentrated in vacuo and the residue was purified using column chromatography, silica gel 100-200 mesh. The product eluted at 15-20% ethyl acetate-pet ether and upon concentration and drying the desired product was obtained as white solid (0.20 g, 81%): $^1$H

NMR (400 MHz, $CDCl_3$): $\delta$ 5.35-5.34 (m, 1H), 4.13 (d, $J$ = 2.3 Hz, 2H), 3.56-3.46 (m, 3H), 2.41 (t, $J$ = 2.3 Hz, 1H), 2.32-2.19 (m, 2H), 2.04-1.93 (m, 2H), 1.88-1.79 (m, 3H), 1.72-1.61 (m, 2H), 1.55-1.03 (m, 17H), 1.00 (s, 3H), 0.93 (d, $J$ = 6.5 Hz, 3H), 0.68 (s, 3H); $^{13}C$ NMR (100 MHz, $CDCl_3$): $\delta$ 140.9, 121.9, 80.2, 74.2, 72.0, 71.0, 58.1, 56.9, 56.1, 50.2, 42.5, 42.4, 39.9, 37.4, 36.6, 35.7, 32.3, 32.0, 31.8, 28.4, 26.2, 24.4, 21.2, 19.5, 18.8, 12.0; FT-IR ($\nu_{max}$, $cm^{-1}$): 3412 (O-H), 3305 (alkyne C-H), 1099 (ether C-O-C); LC-HRMS for $C_{27}H_{42}O_2$ $[M+NH_4]^+$: Calculated: 416.3529, Found: 416.3506.

**Synthesis of fatty acid, Probe 2**

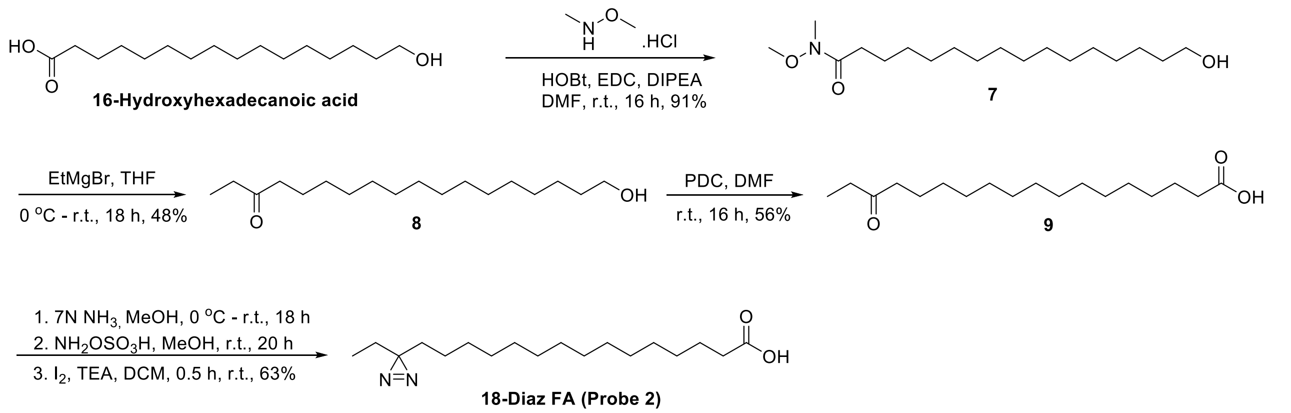


**16-Hydroxy-*N*-methoxy-*N*-methylhexadecanamide, 7**

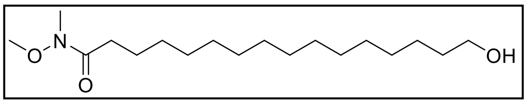


Compound **7** was synthesized using a reported protocol[4]. Briefly, to an ice-cold solution of 16-hydroxyhexadecanoic acid (0.56 g, 2.05 mmol), *N*, *O*-dimethyl hydroxylamine hydrochloride (0.31 g, 1.54 mmol), 1-hydroxybenzotriazole (HOBt, 0.416g, 3.08 mmol) and *N,N*-diisopropylethylamine (DIPEA, 1.07 mL, 6.16 mmol) in dry DMF (10 mL) under nitrogen, EDC.HCl (0.591 g, 3.08 mmol) was added. The reaction mixture was warmed to room temperature and then stirring continued for 16 h. The reaction mixture was diluted with water (100 mL) and then extracted with dichloromethane (3x30 mL). The combined organic layer was given water wash (50 mL), brine wash, dried over anhydrous sodium sulphate and then filtered. The filtrate was concentrated in vacuo, and the residue was purified using column chromatography, silica gel 100-200 mesh. The product eluted at 25-30% ethyl acetate-pet ether and upon concentration and drying the desired product was obtained as white solid (0.59 g, 91%): $^1H$ NMR (400 MHz, $CDCl_3$): $\delta$ 3.64 (s, 3H), 3.57 (t, $J$ = 6.6 Hz, 2H), 3.13(s, 3H), 2.36 (t, $J$ = 7.4 Hz, 2H), 2.30 (bs, 1H), 1.61–1.48 (m, 4H), 1.26 –1.21 (m, 23H). Presence of minor grease impurities were noted.

**18-Hydroxyoctadecan-3-one, 8**

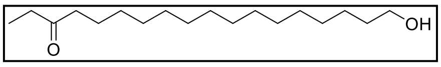

Compound **8** was synthesized using a reported protocol[4]. Briefly, to an ice-cold solution of compound **5** (0.66 g, 2.09 mmol) in dry THF (20 mL) under nitrogen atmosphere, ethyl magnesium bromide (12.55 mL, 12.55 mmol, 1 M in THF) was dropwise added over a period of 10 min. The reaction mixture was warmed to room temperature and stirring was continued for 16 h. The reaction mixture was quenched with saturated ammonium chloride solution (50 mL) and extracted with ethyl acetate (3×30 mL). The combined organic layer was washed with brine, dried over anhydrous sodium sulphate and filtered. The filtrate was concentrated *in vacuo* and the residue was purified using column chromatography, silica gel 100-200 mesh. The product eluted at 10% ethyl acetate-pet ether. The desired product was obtained as white solid (0.3 g, 48%): $^1$H NMR (400 MHz, $CDCl_3$): $\delta$ 3.64 (t, $J$ = 6.6 Hz, 2H), 2.44-2.37 (m, 4H), 1.60-1.53 (m, 5H), 1.31–1.25 (m, 24H), 1.05 (t, $J$ = 7.3 Hz, 3H). Presence of minor grease impurities were noted.

**16-Oxooctadecanoic acid, 9**

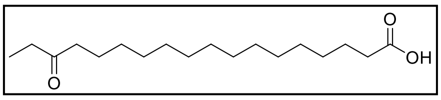

Compound **9** was a reported compound[4]. However, we used another protocol to synthesize it. To a solution of compound **6** (0.3 g, 1.0 mmol) in dry DMF under nitrogen, pyridinium dichromate (PDC, 1.32 g, 3.52 mmol) was added and the reaction mixture was stirred at room temperature for 16 h. The reaction mixture was quenched with water (100 mL), ethyl acetate (20 mL) was added, and the solution was filtered through celite bed. The aqueous layer was extracted with ethyl acetate (3x25 mL). The combined organic layer was given brine wash, dried over anhydrous sodium sulphate and then filtered. The filtrate was concentrated in vacuo, and the residue was purified using column chromatography, silica gel 100-200 mesh. The product eluted at 20-30% ethyl acetate-pet ether and upon concentration and drying the desired product was obtained as white solid (0.176 g, 56%): $^1$H NMR (400 MHz, $CDCl_3$): $\delta$ 2.44-2.33 (m, 6H), 1.67-1.54 (m, 4H), 1.34-1.25 (m, 20H), 1.05 (t, $J$ = 7.3 Hz, 3H).

**15-(3-Ethyl-3*H*-diazirin-3-yl) pentadecanoic acid, 18-diaz-FA (Probe 2)**

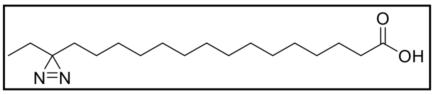

Compound **8** was synthesized using a reported protocol[4]. Briefly, a solution of compound **7** (0.17 g, 0.57 mmol) in dry methanol (1 mL) under nitrogen was cooled to 0 ºC in a 25 ml round bottom flask. 7 N $NH_3$ in methanol (10 mL) was added to the flask, tightly capped with a glass stopper and the resulting reaction mixture was warmed to room temperature and stirring continued for 18 h. The reaction mixture was again cooled to 0 ºC and a solution of hydroxylamine-*O*-sulfonic acid ($NH_2OSO_3H$, 0.074 g, 0.65 mmol) in dry methanol (1 mL) was added dropwise and the round bottom flask was covered with aluminium foil and the reaction mixture was stirred at room temperature for 20 h. Ammonia gas was evaporated under a stream of nitrogen gas and to the remaining solution, diethyl ether (10 mL) was added, leading to the formation of a suspension containing insoluble salts, which were subsequently filtered off. The filtrate was concentrated in vacuo and the residue obtained was re-dissolved in dry DCM (5 mL). To this solution iodine (0.13 g, 0.51 mmol) and triethylamine (TEA, 0.12 mL, 0.85 mmol) were added at room temperature and stirred for 30 min. The reaction solution was concentrated and then purified using column chromatography, silica 100-200 mesh. without doing a workup. The product eluted at 15-20% ethyl acetate-pet ether and upon concentration and drying the desired product was obtained as white solid (0.10 g, 63%): $^1$H NMR (400 MHz, $CDCl_3$): $\delta$ 2.35 (t, $J$ = 7.5 Hz, 2H), 1.64 (quint, $J$ = 7.4 Hz, 2H), 1.41 (q, $J$ = 7.7 Hz, 2H), 1.36-1.24 (m, 22H), 1.09-1.05 (m, 2H), 0.67 (t, $J$ = 7.6 Hz, 3H); LC-HRMS for $C_{18}H_{34}N_2O_2$ $[M-H]^-$: Calculated: 309.2548, Found: 309.2539.

**Synthesis of cholesteryl ester probe, Probe 3 (CS-DA)**

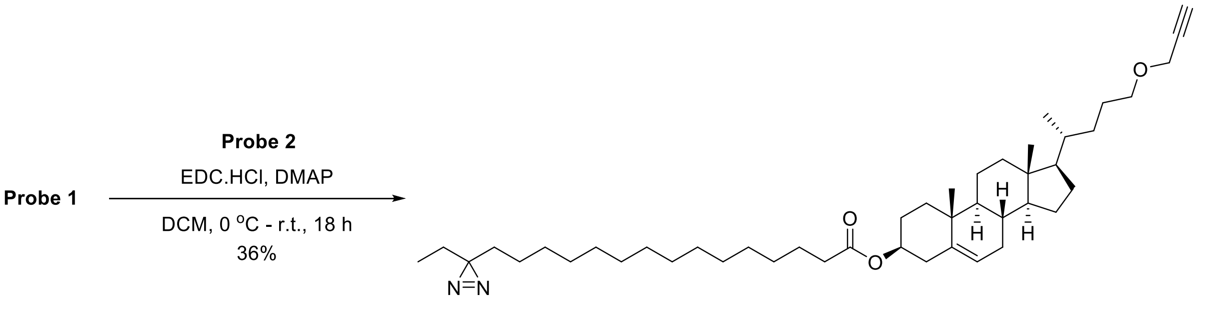


**(3*S*,8*S*,9*S*,10*R*,13*R*,14*S*,17*R*)-10,13-Dimethyl-17-((*R*)-5-(prop-2-yn-1-yloxy) pentan-2-yl)-2,3,4,7,8,9,10,11,12,13,14,15,16,17-tetradecahydro-1*H*-cyclopenta[a]phenanthren-3-yl 15-(3-ethyl-3*H*-diazirin-3-yl) pentadecanoate, CS-DA (Probe 3)**

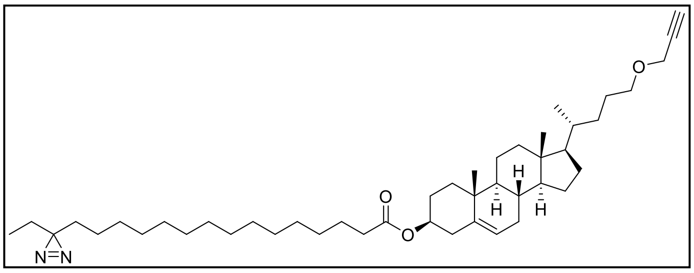

To an ice-cold solution of compound **Probe 1** (0.035 g, 0.089 mmol) and **Probe 2** (0.025 g, 0.081 mmol) in dry dichloromethane (5 mL), under nitrogen atmosphere, 1-ethyl-3-(3- dimethylaminopropyl) carbodiimide hydrochloride (EDC.HCl, 0.023 g, 0.121 mmol) and 4-(dimethylamino) pyridine (DMAP, 0.9 mg, 0.008 mmol) were added. The reaction mixture was then warmed to room temperature and stirring continued for 16 h. The progress of reaction was monitored using TLC. The reaction solution was concentrated and then purified using column chromatography, silica 100-200 mesh, without doing a workup. The product eluted at 2% ethyl acetate-pet ether and upon concentration and drying the desired product was obtained as white solid (0.019 g, 43%): $^{1}$H NMR (400 MHz, $CDCl_3$): $\delta$ 5.37 (d, $J$ = 4.2 Hz, 1H), 4.65-4.57 (m, 1H), 4.13 (d, $J$ = 2.3 Hz, 2H), 3.50-3.47 (m, 2H), 2.41 (t, $J$ = 2.3 Hz, 1H), 2.32-2.24 (m, 4H), 2.02-1.93 (m, 2H), 1.87-1.79 (m, 3H), 1.72-1.58 (m ,4H), 1.50-1.04 (m, 42H), 1.02 (s, 3H), 0.93 (d, $J$ = 6.5 Hz, 3H), 0.70-0.65 (m, 6 H); $^{13}$C NMR (100 MHz, $CDCl_3$): $\delta$ 173.5, 139.9, 122.7, 80.2, 74.2, 73.8, 71.0, 58.1, 56.8, 56.1, 50.1, 42.5, 39.9, 38.3, 37.1, 36.7, 35.7, 34.9, 32.7, 32.3, 32.0, 29.8, 29.7, 29.6, 29.4, 29.3, 28.3, 28.0, 26.3, 26.1, 25.2, 24.4, 24.0, 21.2, 19.5, 18.8, 12.0, 8.4; FT-IR ($\nu_{max}$, $cm^{-1}$): 3305 (alkyne C-H), 1733 (ester C=O), 1577 (N=N), 1100 (ether C-O-C); LC-HRMS for $C_{45}H_{74}N_2O_3$ $[M+NH_4]^+$: Calculated: 708.6043, Found: 708.6010.

##### Synthesis of fatty acid probe, Probe 4 (PA-DA)

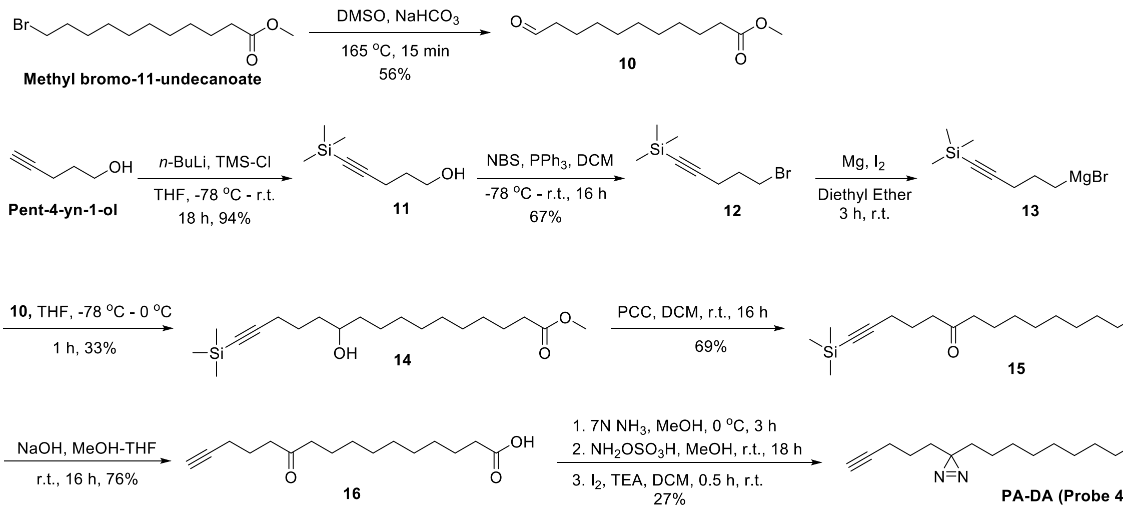


##### Methyl 11-oxoundecanoate, 10

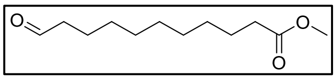

Compound **10** was synthesized using a reported protocol[5,6]. Briefly, to a solution of methyl 11-bromoundecanoate (0.8 g, 2.81 mmol) in DMSO (8 mL), sodium bicarbonate (0.52 g, 6.18 mmol) was added, and the reaction mixture was refluxed at 165 ºC for 15 min. The reaction mixture was cooled to room temperature, diluted with ice cold water (50 mL), and extracted with ethyl acetate (3x40 mL). The combined organic layer was washed with water (50 mL), saturated sodium bicarbonate solution (30 mL) and brine. The resulting organic layer was dried over anhydrous sodium sulphate, filtered and the filtrate was concentrated in vacuo. The crude product was purified using column chromatography, silica 100-200 mesh. The product eluted at 10% ethyl acetate-pet ether and upon concentration and drying the desired product was obtained as colourless oil (0.345 g, 56%): $^1$H NMR (400 MHz, $CDCl_3$): $\delta$ 9.76 (t, $J$ = 1.8 Hz, 1H), 3.66 (s, 3H), 2.41(td, $J$ = 7.3, 1.8 Hz, 2H), 2.30 (t, $J$ = 7.5 Hz, 2H), 1.62–1.59 (m, 4H), 1.29 (m, 10H).

**5-(Trimethylsilyl) pent-4-yn-1-ol, 11**

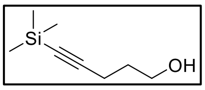


Compound **11** was synthesized using a reported protocol[5,6]. Briefly, in a two necked flame-dried 250 ml round-bottom flask, 4-pentyn-1-ol (5.0 g, 59.44 mmol) was dissolved in anhydrous THF (100 mL) and then cooled to -78 ºC. To this, *n*-butyl-lithium (*n*-BuLi, 82 mL, 1.6 M in hexanes, 130.76 mmol) was dropwise added over a period of 30 min. After one hour stirring at -78 ºC, chlorotrimethylsilane (TMS-Cl, 22.63 mL, 178.32 mmol) was added dropwise to the reaction mixture and then gradually warmed to room temperature. The reaction was stirred for an additional 16 hours at room temperature. Consumption of starting material was monitored with TLC. The reaction mixture was subsequently cooled to 0 ºC and acidified with 1 M HCl (150 mL), followed by stirring for 1 h. The resulting mixture was then extracted with ethyl acetate (3×100 mL). The combined organic layer was washed with water (250 mL), saturated sodium bicarbonate solution (200 mL), brine, dried over anhydrous sodium sulphate and then filtered. The filtrate was concentrated in vacuo, and the obtained residue was purified using column chromatography, silica 100-200 mesh. The product eluted at 2% ethyl acetate-pet ether and upon concentration and drying the desired product was obtained as colourless oil (9 g, 94%): $^1$H NMR (400 MHz, $CDCl_3$): $\delta$ 3.75 (t, $J$ = 6.0 Hz, 2H), 2.34 (t, $J$ = 6.9 Hz, 2H), 1.76 (quint, $J$ = 6.6 Hz, 2H), 0.14 (s, 9H).

**(5-Bromopent-1-yn-1-yl) trimethylsilane, 12**

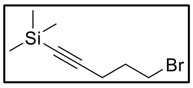


Compound **12** was synthesized using a reported protocol[5,6]. Briefly, a solution of compound **11** (9.0 g, 57.58 mmol) in anhydrous dichloromethane (150 mL) under nitrogen atmosphere, was cooled to -78 ºC. Sequentially triphenylphosphine ($PPh_3$, 22.6 g, 86.37 mmol) and portion wise *N*-bromosuccinimide (NBS, 15.37 g, 86.37 mmol) were added. The reaction mixture was gradually warmed to room temperature and stirred for 6 h. The progress of reaction was monitored using TLC and upon completion, the reaction mixture was diluted with ethyl acetate (500 mL) and washed with sat. $NaHCO_3$ (2 x 300 mL). The organic layer was then washed with brine, dried over $Na_2SO_4$, filtered and the filtrate was concentrated in vacuo. The resulting crude was purified by silica gel column chromatography (elution with 100% hexanes) to afford the **2** as a colourless oil. This bromide was further distilled under vacuum (5-10 mm Hg (Torr), 90-100 ºC) to obtain colorless oil (70-80% yield): $^1$H NMR (400 MHz, $CDCl_3$): $\delta$ 3.51 (t, $J$ = 6.5 Hz, 2H), 2.41 (t, $J$ = 6.8 Hz, 2H), 2.04 (quint, $J$ = 6.6 Hz, 2H), 0.15 (s, 9H).

**Methyl 11-hydroxy-16-(trimethylsilyl) hexadec-15-ynoate, 14**

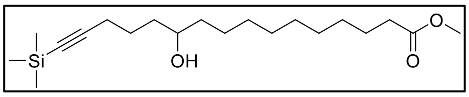

Compound **14** was synthesized using a reported protocol[6] in steps.

1. *Preparation of Grignard reagent*[6]: Reaction flasks, take offs, spatulas, syringes, needles and magnetic stir bars were dried at 110 °C in an oven for 2 h. Magnesium turnings were activated by washing with a 10% aq. HCl solution (3x), followed by subsequent rinsing with water and acetone. The magnesium turnings were then dried in the oven at 110 ºC for 8 h. All items in the oven were gradually cooled to room temperature in a desiccator. Diethyl ether used for the preparation of Grignard reagent was dried over sodium metal and then distilled. Dry tetrahydrofuran available from commercial sources was used for the reaction. The reaction was performed using a Schlenk line.
2. *(5-(Trimethylsilyl)pent-4-yn-1-yl)magnesium bromide*, **13**[6]: A 100 mL round-bottom flask, charged with the magnesium turnings (0.114 g, 4.69 mmol), was heated with a heat gun for 10 min under vacuum, and then cooled to room temperature under nitrogen atmosphere. A magnetic stir bar and catalytic amount of iodine (0.006 g, 0.022 mmol) were added into the flask followed by addition of anhydrous diethyl ether (10 mL). The mixture was stirred at room temperature for 10 min. A solution of **12** (0.822 g, 3.75 mmol) in anhydrous diethyl ether (10 mL) was prepared and a small portion, approximately 40%, was added dropwise, and then the reaction mixture was stirred at room temperature for 10 min until the colour of the solution transitioned from brown/red to greyish, signifying the initiation of the Grignard reagent. The remaining solution of **12** was then added dropwise over 10 min and the reaction mixture was stirred at room temperature for another 3 h. The Grignard reagent **13** was used immediately for the next step without characterization.
3. *Grignard reaction*[6]*:* To a chilled solution of methyl 11-oxoundecanoate **10** (0.670 g, 3.13 mmol) in dry THF (10 mL) at -78 ºC, under nitrogen atmosphere, was added Grignard solution **13** dropwise over 10 min. The reaction mixture was then gently warmed to 0 ºC and stirred for 1 h. The reaction mixture was quenched with saturated ammonium chloride solution (150 mL) and the product was extracted with ethyl acetate (3x70 mL). The combined organic layer was dried with anhydrous sodium sulphate, filtered and the filtrate was concentrated in vacuo. The crude product was purified using column chromatography, silica 100-200 mesh. The product eluted at 15% ethyl acetate-pet ether and upon concentration and drying the desired product was obtained as colorless oil (0.373 g, 33%): $^{1}$H NMR (400 MHz, $CDCl_3$): $\delta$ 3.65 (s, 3H), 3.63–3.59 (m, 1H), 2.31-2.23 (m, 4H), 1.69-1.38 (m, 10H), 1.27 (m, 10H), 0.13 (s, 9H).

**Methyl 11-oxo-16-(trimethylsilyl) hexadec-15-ynoate, 15**

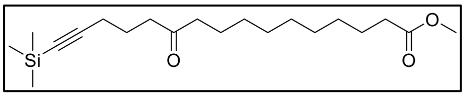

Compound **15** was a reported compound[6]. However, we used another protocol to synthesize it. To an ice-cold solution of compound **14** (0.373 g, 1.05 mmol) in dry dichloromethane, under nitrogen, was added pyridinium chlorochromate (PCC, 0.34 g, 1.58 mmol) and then the reaction mixture was warmed to room temperature. Stirring was continued at room temperature for 16 h. The reaction mixture was filtered through celite bed, and the filtrate was then concentrated in vacuo. The crude product was purified using column chromatography, silica 100-200 mesh. The product eluted at 5% ethyl acetate-pet ether and upon concentration and drying the desired product was obtained as white solid (0.250 g, 67%): $^{1}$H NMR (400 MHz, $CDCl_3$): $\delta$ 3.65 (s, 3H), 2.52 (t, $J$ = 7.2 Hz, 2H), 2.39 (t, $J$ = 7.4 Hz, 2H), 2.29 (t, $J$ = 7.5 Hz, 2H), 2.24 (t, $J$ = 7.0 Hz, 2H), 1.76 (quint, $J$ = 7.0 Hz, 2H), 1.62-1.54 (m, 4H), 1.27 (m, 10H), 0.13 (s, 9H).

**11-Oxohexadec-15-ynoic acid, 16**

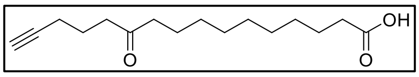

Compound **16** was synthesized using a reported protocol[6]. Briefly, to a solution of **15** (0.248 g, 0.70 mmol) in MeOH (4 mL), a solution of sodium hydroxide (NaOH, 0.14 g, 3.52 mmol) in water (4 mL) was dropwise added and the reaction mixture was stirred at room temperature for 16 h. The reaction mixture was acidified to pH 2-3 by treating with 10% aq. HCl at 0 ºC and extracted with ethyl acetate (3×30 mL). The combined organic layer was washed with brine (50 mL), dried over anhydrous sodium sulphate, filtered and the filtrate was concentrated in vacuo The crude product was purified using column chromatography, silica 100-200 mesh. The product eluted at 30-40% ethyl acetate-pet ether and upon concentration and drying the desired product was obtained as white solid (0.14 g, 76%): $^{1}$H NMR (400 MHz, $CDCl_3$): $\delta$ 2.55 (t, $J$ = 7.2 Hz, 2H), 2.40 (t, $J$ = 7.4 Hz, 2H), 2.34 (t, $J$ = 7.4 Hz, 2H), 2.22 (td, $J$ = 6.9, 2.6 Hz, 2H), 1.95 (t, $J$ = 2.6 Hz, 1H), 1.78 (quint, $J$ = 7.0 Hz, 2H), 1.66-1.54 (m, 4H), 1.34-1.27 (m, 10H).

**10-(3-(pent-4-yn-1-yl)-3*H*-diazirin-3-yl) decanoic acid, PA-DA (Probe 4)**

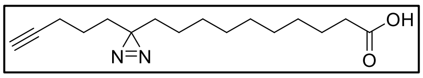

**PA-DA** (**Probe 4**) was synthesized using a reported protocol[5,6]. Briefly, in a 25 ml round bottom flask, compound **16** (0.108 g, 0.41 mmol) was added and cooled to 0 ºC. 7 N $NH_3$ in MeOH (10 mL) was added, tightly capped with a glass stopper and the resulting reaction mixture was stirred at 0 ºC for 3 h. To this cold solution, a solution of $NH_2OSO_3H$ (0.053 g, 0.466 mmol) in MeOH (1 mL) was dropwise added and the round bottom flask was covered with aluminium foil and the reaction mixture was stirred at room temperature for 16 h. The solvent was evaporated under a stream of nitrogen gas and to the remaining residue, diethyl ether (10 mL) was added, leading to the formation of a suspension containing insoluble salts, which were subsequently filtered off. The filtrate was concentrated in vacuo and the residue obtained was re-dissolved in dry DCM (2 mL). To this solution iodine (0.148 g, 0.58 mmol) and triethylamine (0.041 mL, 0.61 mmol) were added at room temperature and stirred for 30 min. The reaction solution was concentrated and then purified using column chromatography, silica 100-200 mesh. without doing a workup. The product eluted at 25-30% ethyl acetate-pet ether and upon concentration and drying the desired product was obtained as pale-yellow oil (30 mg, 27%): $^{1}$H NMR (400 MHz, $CDCl_3$): $\delta$ 2.34 (t, $J$ = 7.1 Hz, 2H), 2.15 (td, $J$ = 6.8, 2.3 Hz, 2H), 1.94 (t, $J$ = 2.6 Hz, 1H), 1.61 (quint, $J$ = 6.3 Hz, 2H), 1.50–1.46 (m, 2H), 1.38–1.23 (m, 14H), 1.10-1.06 (m, 2H); LC-HRMS for $C_{16}H_{26}N_2O_2$ $[M-H]^-$: Calculated: 277.1922, Found: 277.1910.

**Synthesis of cholesteryl ester probe, Probe 5 (CP-DA)**

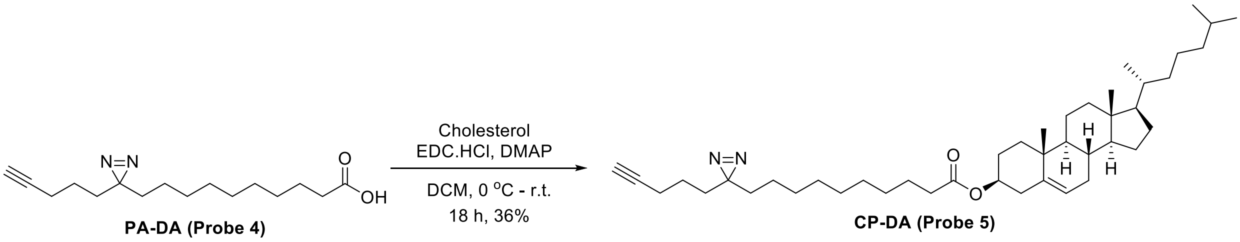


**((3*S*,8*S*,9*S*,10*R*,13*R*,14*S*,17*R*)-10,13-Dimethyl-17-((*R*)-6-methylheptan-2-yl)-2,3,4,7,8,9,10,11,12,13,14,15,16,17-tetradecahydro-1*H*-cyclopenta[a]phenanthren-3-yl 10-(3-(pent-4-yn-1-yl)-3*H*-diazirin-3-yl) decanoate, CP-DA (Probe 5)**

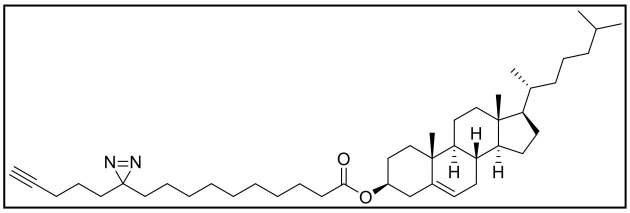


To an ice-cold solution of **PA-DA** (**Probe 4**, 0.028 g, 0.101 mmol) and cholesterol (0.042 g, 0.11 mmol) in dry DCM (5 mL), under nitrogen atmosphere, EDC.HCl (0.023 g, 0.12 mmol) and DMAP (1.2 mg, 0.01 mmol) were added. The reaction mixture was then warmed to room temperature and stirring continued for 16 h. The progress of reaction was monitored using TLC. The reaction solution was concentrated and then purified using column chromatography, silica 100-200 mesh, without doing a workup. The product eluted at 2% ethyl acetate-pet ether and upon concentration and drying the desired product was obtained as white solid (25 mg, 36%):$^{1}$H NMR (400 MHz, $CDCl_3$): $\delta$ 5.37 (d, $J$ = 4.0 Hz, 1H), 4.65-4.57 (m, 1H), 2.30 (d, $J$ = 7.5 Hz, 2H), 2.26 (t, $J$ = 7.5 Hz, 2H), 2.16 (td, $J$ = 6.9, 2.6 Hz, 2H), 2.02-1.95 (m, 2H), 1.94 (t, $J$ = 2.6 Hz, 1H), 1.87-1.78 (m, 3H), 1.61-1.58 (m, 2H), 1.56-1.27 (m, 23H),

1.23-0.94 (m, 21H), 0.91 (d, $J$ = 6.5 Hz, 3H), 0.87 (d, $J$ = 1.6 Hz, 3H), 0.85 (d, $J$ = 1.6 Hz, 3H), 0.67 (s, 3H); $^{13}$C NMR (100 MHz, $CDCl_3$): $\delta$ 173.4, 139.9, 122.7, 83.6, 73.8, 69.0, 56.8, 56.3, 50.2, 42.5, 39.9, 39.7, 38.3, 37.1, 36.7, 36.3, 35.9, 34.8, 33.0, 32.1, 32.0, 29.4, 29.3, 29.2, 28.6, 28.4, 28.2, 28.0, 25.2, 24.4, 24.0, 23.0, 22.9, 22.7, 21.2, 19.5, 18.9, 18.1, 12.0. FT-IR ($\nu_{max}$, cm$^{-1}$): 3310 (alkyne C-H), 1733 (ester C=O), 1582 (N=N); LC-HRMS for $C_{43}H_{70}N_2O_2$ $[M+NH_4]^+$: Calculated: 664.5781, Found: 664.5761.

**Synthesis of cholesterol probe, Probe 6 (Ch-DA)**

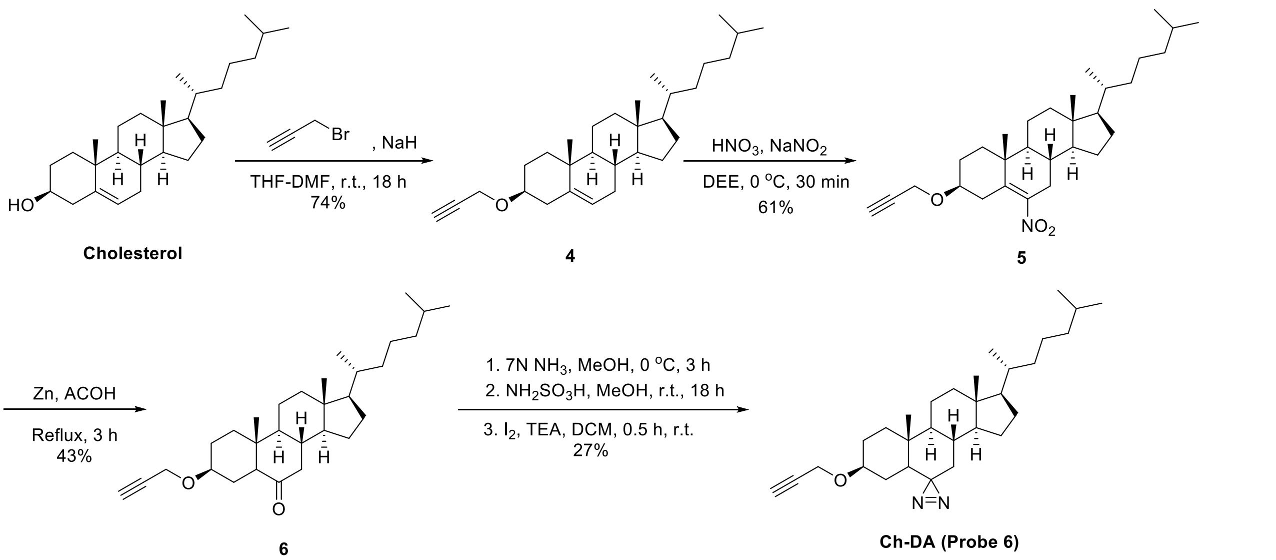


**(3*S*,8*S*,9*S*,10*R*,13*R*,14*S*,17*R*)-10,13-Dimethyl-17-((*R*)-6-methylheptan-2-yl)-3-(prop-2-yn-1-yloxy)-2,3,4,7,8,9,10,11,12,13,14,15,16,17-tetradecahydro-1*H*-cyclopenta[*a*]phenanthrene, 4**

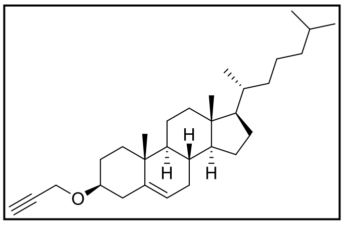

Compound **4** was synthesized using a reported protocol with slight modification[7]. Briefly, NaH (0.124 g, 3.10 mmol, 60% dispersion in oil) in a dry THF (5 mL) and DMF (1.5 mL) was stirred for 15 min under nitrogen atmosphere. Cholesterol (0.4 g, 1.03 mmol) was added to the solution and stirred at room temperature for 30 min. Propargyl bromide (0.293 mL, 1.55 mmol, 80% in toluene) was then added dropwise to the reaction mixture and stirring continued at room temperature for 16 h done. The reaction was quenched using water (100 mL) and extracted using ethyl acetate (3x25 mL). The combined organic layer was again washed with water, brine and dried over anhydrous sodium sulphate and then filtered. The filtrate was concentrated and the residue was purified using column chromatography, silica gel 100-200 mesh. The product eluted at 3% ethyl acetate-pet ether and upon concentration and drying the desired product was obtained as white waxy solid (0.326 g, 74%): $^1$H NMR (400 MHz, $CDCl_3$): $\delta$ 5.37-5.36 (m, 1H), 4.19 (d, $J$ = 2.4 Hz, 2H), 3.42-3.34 (m, 1H), 2.41-2.36 (m, 2H), 2.25-2.18 (m,

1H), 2.03-1.78 (m, 5H), 1.53-1.27 (m, 11H), 1.22-0.94 (m, 13H), 0.91 (d, *J* = 6.5 Hz, 3H), 0.87 (d, *J* = 1.8 Hz, 3H), 0.85 (d, *J* = 1.8 Hz, 3H), 0.67 (s, 3H).

**(3*S*,8*S*,9*S*,10*R*,13*R*,14*S*,17*R*)-10,13-Dimethyl-17-((*R*)-6-methylheptan-2-yl)-6-nitro-3-(prop-2-yn-1-yloxy)-2,3,4,7,8,9,10,11,12,13,14,15,16,17-tetradecahydro-1*H*-cyclopenta[*a*]phenanthrene, 5**

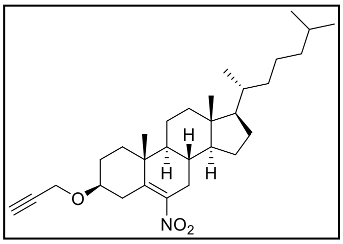


To an ice-cold solution of **4** (0.445 g, 1.05 mmol) in diethyl ether (15mL), concentrated nitric acid ($HNO_3$, 15 mL) was added dropwise using a dropping funnel. The solution was stirred in ice bath for 10 min. Sodium nitrite ($NaNO_2$, 0.13 mg, 1.88 mmol) was added and the reaction mixture was stirred at 0 °C for additional 10-15 min. The reaction was complete with consumption of **1**, TLC monitored. The reaction mixture was then quenched using ice-cold water and the aqueous layer was extracted with ethyl acetate (3x25 mL). The combined organic layer was washed with brine, dried over anhydrous sodium sulphate and then filtered. The filtrate was concentrated and the residue was purified using column chromatography, silica gel 100-200 mesh. The product eluted at 8% ethyl acetate-pet ether and upon concentration and drying the desired product was obtained as pale yellow waxy solid (0.3 g, 61%): $^1$H NMR (400 MHz, $CDCl_3$): $\delta$ 4.22-4.13 (m, 2H), 3.49-3.41 (m, 1H), 2.91-2.86 (m, 1H), 2.56-2.50 (m, 1H), 2.41 (t, *J* = 2.4 Hz, 1H), 2.15-1.95 (m, 5H), 1.90-1.82 (m, 1H), 1.67-1.58 (m, 2H), 1.53-1.28 (m, 8H), 1.23-0.96 (m, 14H), 0.91 (d, *J* = 6.5 Hz, 3H), 0.87 (d, *J* = 1.8 Hz, 3H), 0.85 (d, *J* = 1.8 Hz, 3H), 0.68 (s, 3H); $^{13}$C NMR (100 MHz, $CDCl_3$): $\delta$ 146.5, 138.4, 80.0, 74.4, 56.3, 56.1, 55.8, 49.2, 42.4, 39.6, 39.5, 38.2, 36.7, 36.3, 35.9, 33.5, 31.9, 31.8, 28.2, 28.1, 27.8, 24.3, 23.9, 23.0, 22.7, 21.1, 19.7, 18.8, 11.9; FT-IR ($\nu_{max}$, cm$^{-1}$): 3306 (alkyne C-H), 1518 (-$NO_2$), 1358 (-$NO_2$), 1084 (ether C-O-C); LC-HRMS for $C_{30}H_{47}NO_3$ [M+ $NH_4$]$^+$: Calculated: 487.3900, Found: 487.3869.

**(3*S*,8*S*,9*S*,10*R*,13*R*,14*S*,17*R*)-10,13-Dimethyl-17-((*R*)-6-methylheptan-2-yl)-3-(prop-2-yn-1-yloxy) hexadecahydro-6*H*-cyclopenta[*a*]phenanthren-6-one, 6**

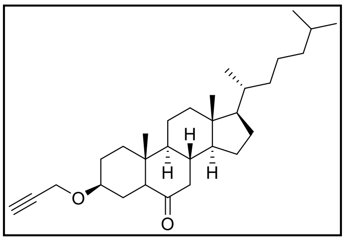


To a solution of **5** (0.6 g, 1.24 mmol) in acetic acid (10 mL) and water (1.5 mL), zinc powder (0.232 g, 3.72 mmol) was added in portions and then heated to reflux for 3 h. The reaction mixture was cooled, diluted with ethyl acetate and filtered over celite to remove zinc residue. The organic layer was washed with water, brine, dried over anhydrous sodium sulphate and then filtered. The filtrate was concentrated and the residue was purified using column chromatography, silica gel 100-200 mesh. The product eluted at 5% ethyl acetate-pet ether and upon concentration and drying the desired product was obtained as off white waxy solid (0.242

mg, 43%): $^1$H NMR (400 MHz, $CDCl_3$): $\delta$ 4.20 (qd, $J$ = 7.2, 2.4 Hz, 2H), 3.50-3.42 (m, 1H), 2.39 (t, $J$ = 2.3 Hz, 1H), 2.32 (dd, $J$ = 13.0, 4.4 Hz, 1H), 2.17 (dd, $J$ = 12.5, 2.5 Hz, 1H), 2.06-1.97 (m, 2H), 1.97-1.74 (m, 5H), 1.63-1.59 (m, 1H), 1.55-1.46 (m, 2H), 1.45-1.31 (m, 6H), 1.28-0.95 (m, 12H), 0.91 (d, $J$ = 6.5 Hz, 3H), 0.87 (d, $J$ = 1.7 Hz, 3H), 0.85 (d, $J$ = 1.6 Hz, 3H), 0.74 (s, 3H), 0.68 (s, 3H); $^{13}$C NMR (100 MHz, $CDCl_3$): $\delta$ 211.0, 80.4, 76.7, 74.0, 56.9, 56.8, 56.3, 55.1, 54.1, 46.9, 43.1, 41.3, 39.7, 39.6, 38.0, 36.8, 36.2, 35.8, 28.2, 28.1, 27.8, 26.0, 24.1, 23.9, 22.9, 22.7, 21.6, 18.8, 13.2, 12.2; FT-IR ($\nu_{max}$, cm$^{-1}$): 3308 (alkyne C-H), 1708 (C=O), 1085 (ether C-O-C); LC-HRMS for $C_{30}H_{48}O_2$ [M+ $NH_4$]$^+$: Calculated: 458.3998, Found: 458.3973.

**(3*S*,8*S*,9*S*,10*R*,13*R*,14*S*,17*R*)-10,13-Dimethyl-17-((*R*)-6-methylheptan-2-yl)-3-(prop-2-yn-1-yloxy)-1,2,3,4,5,7,8,9,10,11,12,13,14,15,16,17-hexadecahydrospiro[cyclopenta[*a*]phenanthrene-6,3'-diazirine], Ch-DA (Probe 6)**

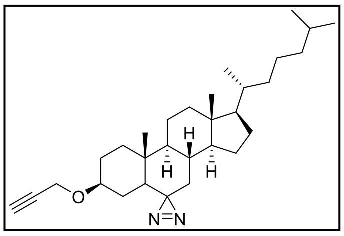


In a sealed tube under nitrogen atmosphere, compound **6** (0.143 g, 0.32 mmol) was dissolved in dry dichloromethane (2 mL) and cooled using an ice bath. 7 N ammonia solution in methanol (10 mL) was added to the reaction mixture and the reaction was stirred at 0 °C for 3 h. $NH_2OSO_3H$ (0.042 mg, 0.37 mmol) was added at 0 °C and the reaction was warmed to room temperature and stirring continued for 16 h. Ammonia gas was evaporated from the reaction mixture using nitrogen flow and then concentrated. The residue was dissolved in dry DCM (10 mL) and sequentially, iodine (0.118 g, 0.47 mmol) and triethylamine (0.067 mL, 0.49 mmol) were added. The reaction mixture was stirred at room temperature for 30 min and then concentrated. The residue was purified using column chromatography, silica gel 100-200 mesh, without workup. The product eluted at 2% ethyl acetate-pet ether and upon concentration and drying the desired product was obtained as white waxy solid (0.04 g, 22%): $^1$H NMR (400 MHz, $CDCl_3$): $\delta$ 4.13-4.04 (m, 2H), 3.37-3.30 (m, 1H), 2.37 (t, $J$ = 2.4 Hz, 1H), 2.02 (dt, $J$ = 12.7, 3.2 Hz, 1H), 1.84-1.72 (m, 4H), 1.68 (dd, $J$ = 13.4, 3.2 Hz, 1H), 1.54-0.93 (m, 23H), 0.90 (d, $J$ = 6.5 Hz, 3H), 0.86 (d, $J$ = 1.6 Hz, 3H), 0.85 (d, $J$ = 1.6 Hz, 3H), 0.83-0.76 (m, 1H), 0.69 (s, 3H), 0.44-0.34 (m, 2H); $^{13}$C NMR (100 MHz, $CDCl_3$): $\delta$ 80.3, 77.3, 74.0, 56.3, 56.2, 55.1, 53.9, 45.3, 42.8, 39.9, 39.6, 37.9, 37.5, 36.4, 36.3, 35.9, 34.0, 29.6, 28.2, 27.5, 24.0, 23.9, 23.0, 22.7, 21.3, 18.8, 13.1, 12.2; FT-IR ($\nu_{max}$, cm$^{-1}$): 3308 (alkyne C-H), 1582 (N=N), 1089 (ether C-O-C); LC-HRMS for $C_{30}H_{48}N_2O$ [M+ $NH_4$]$^+$: Calculated: 470.4110, Found: 470.4072.

### CHARACTERIZATION SPECTRA OF PROBES

[1]H NMR of **Probe 1**

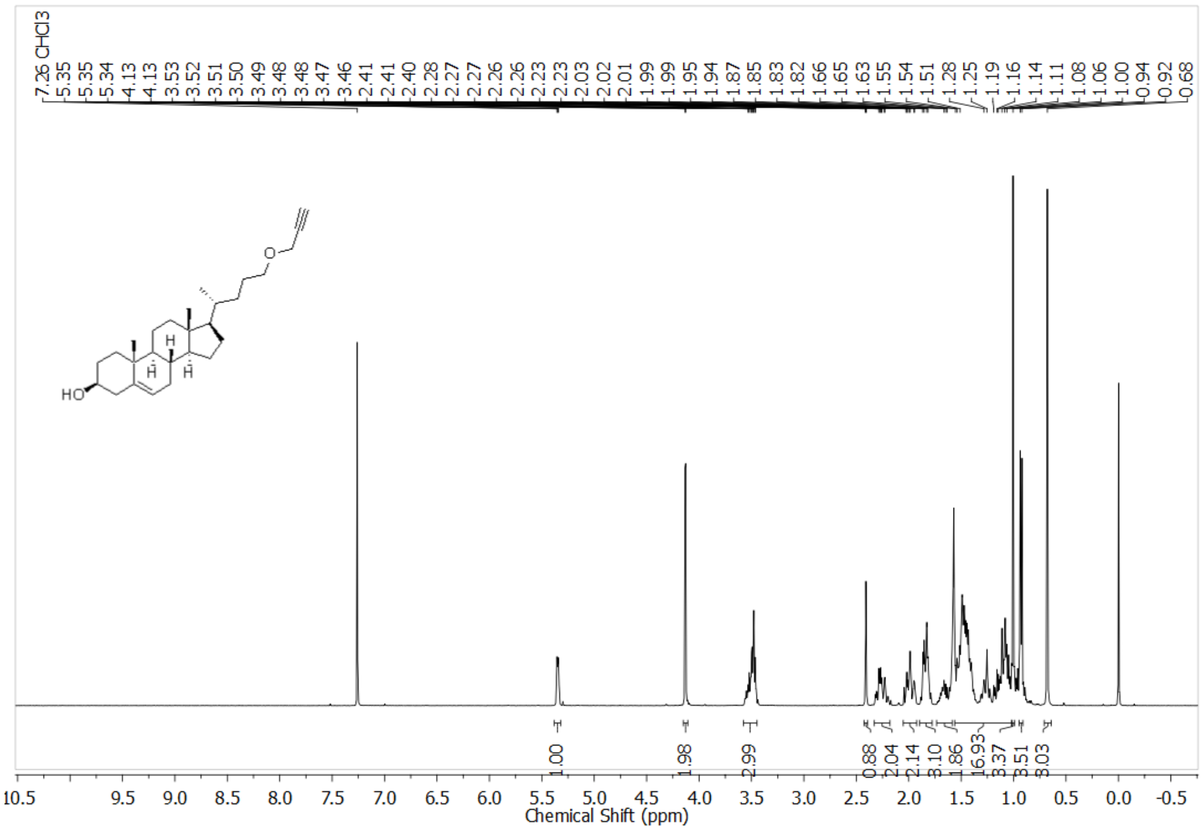


[13]C NMR of **Probe 1**

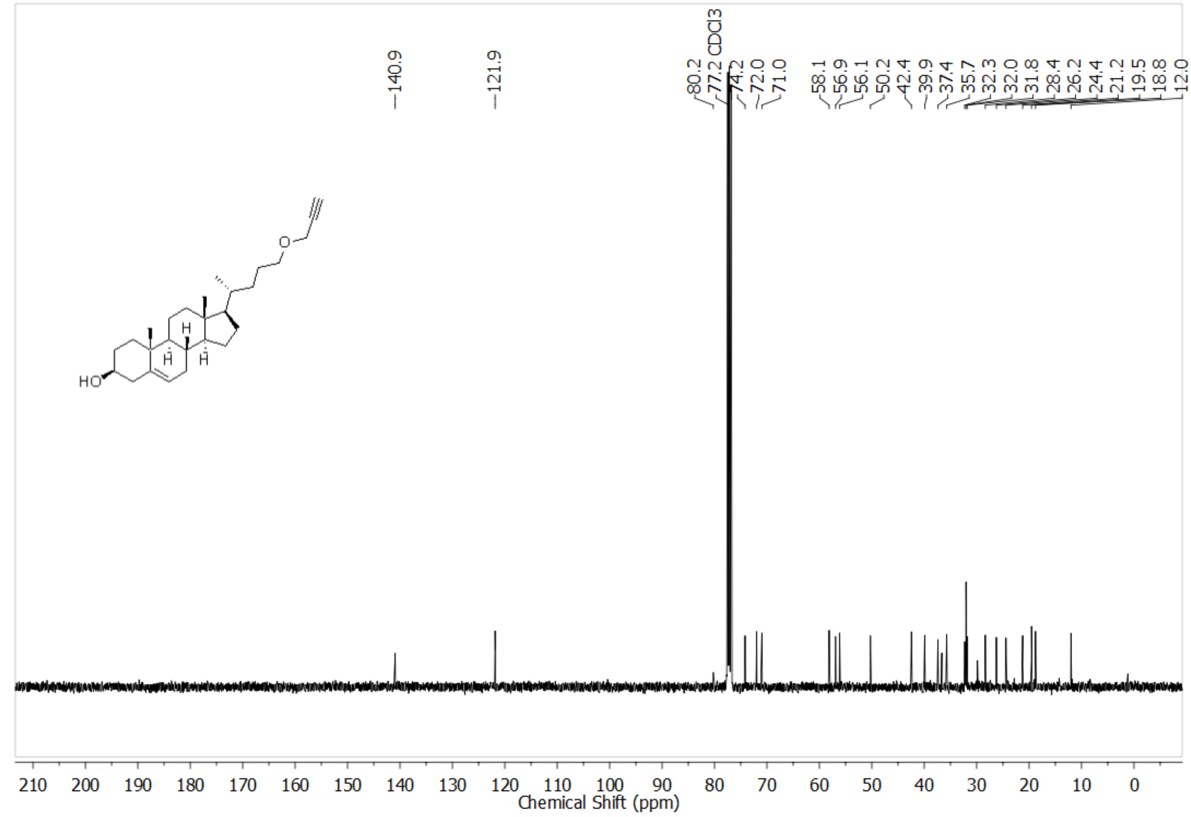

FTIR spectra of **Probe 1**

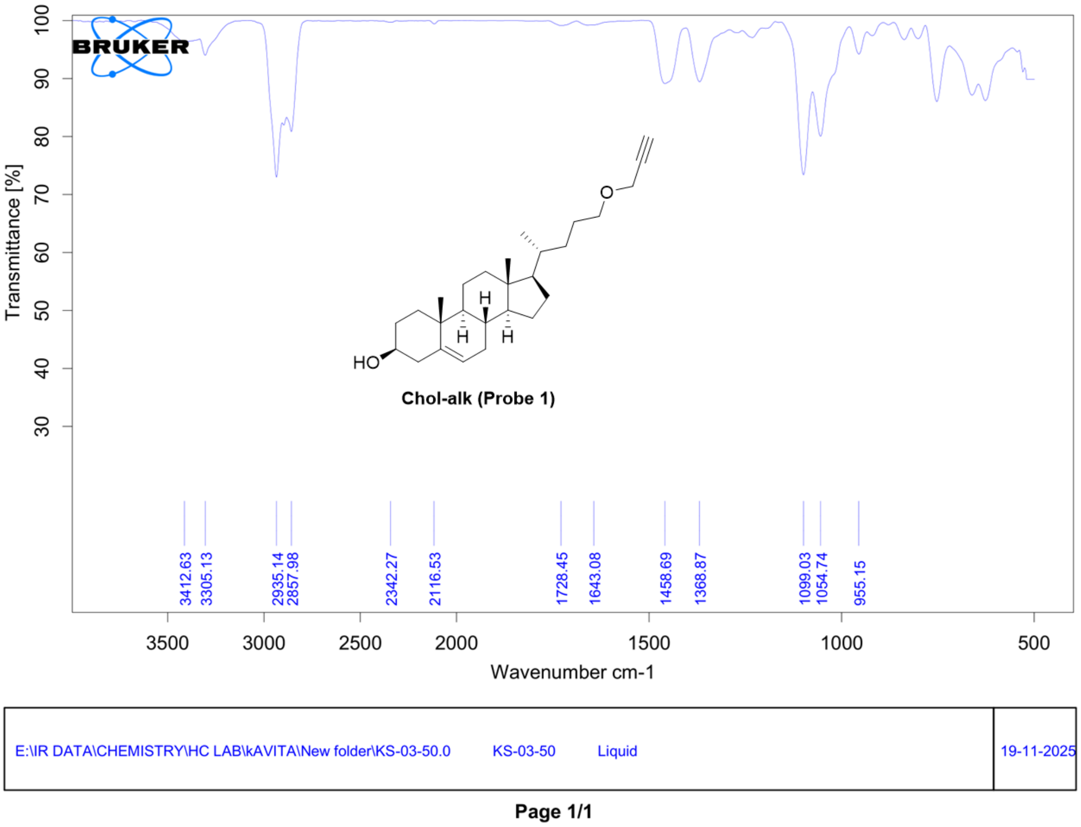


HRMS of **Probe 1**

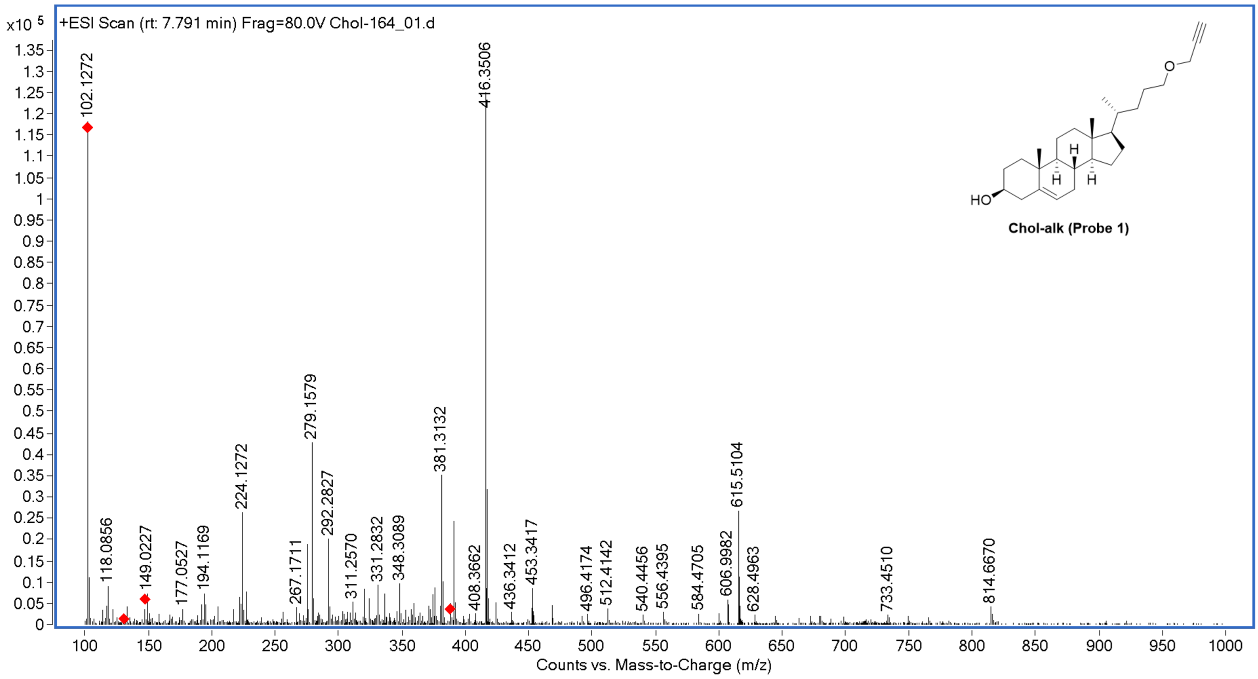

$^{1}H$ NMR of **Probe 2**

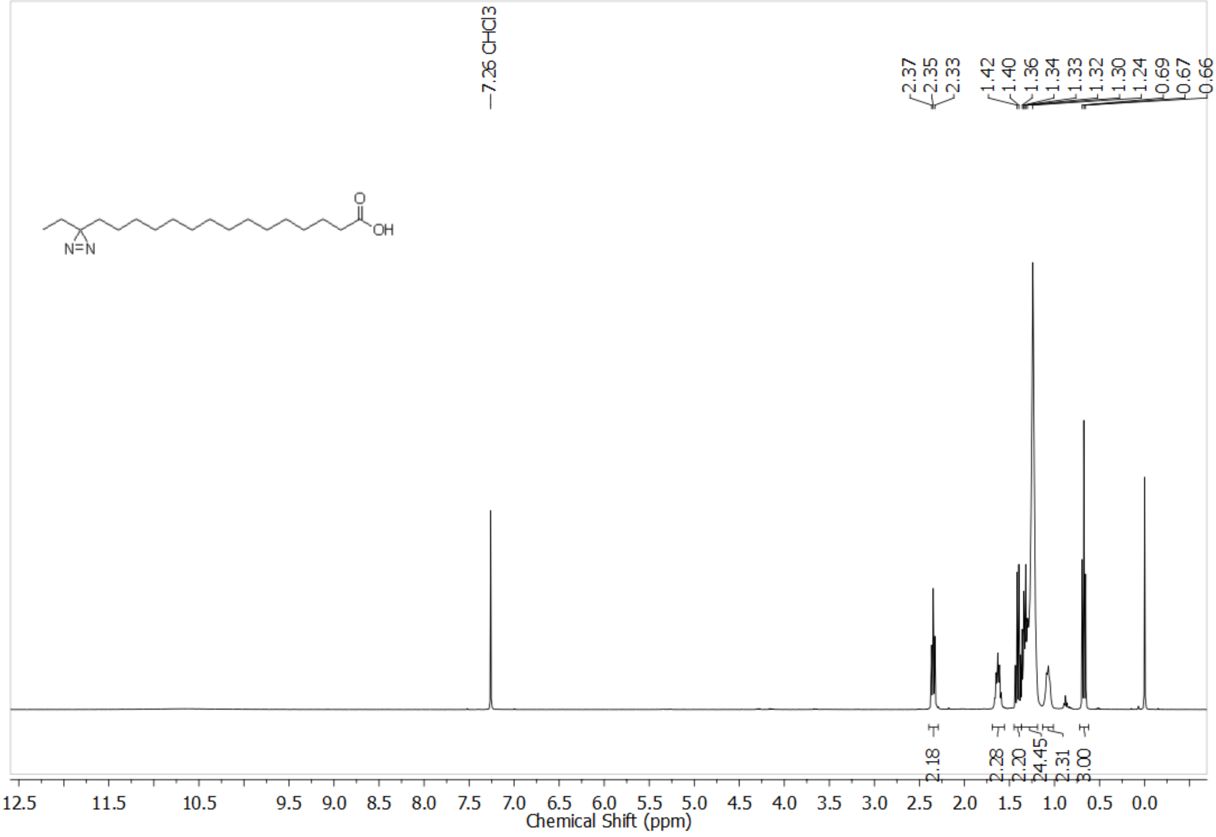


HRMS of **Probe 2**

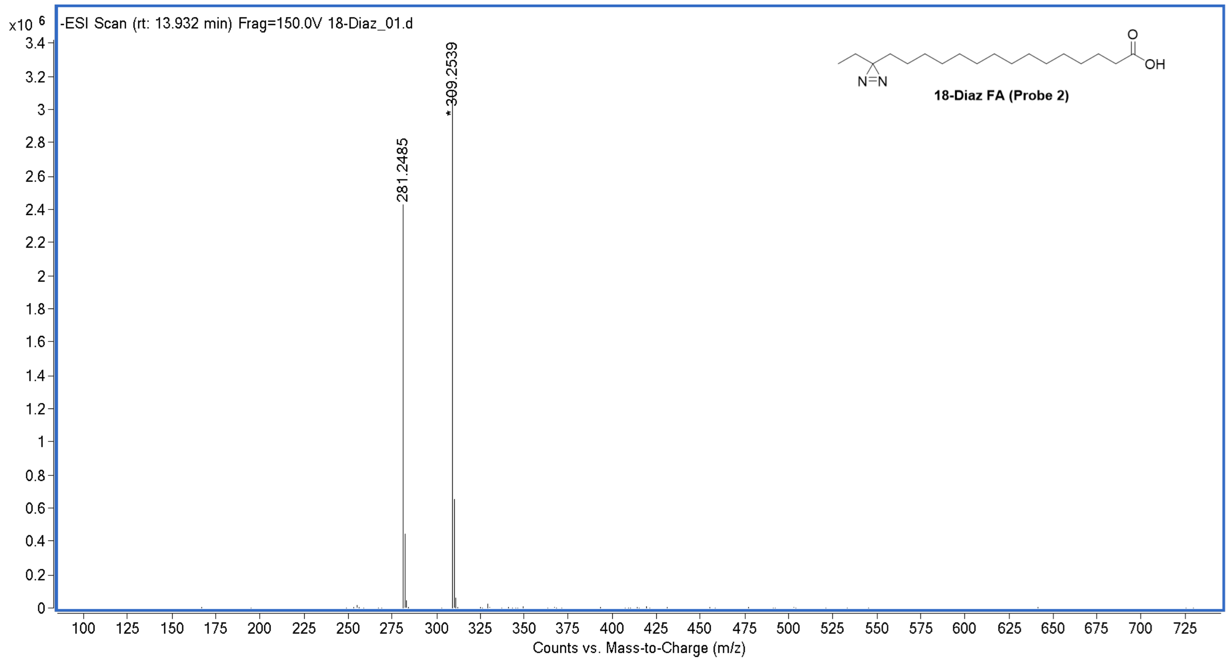

$^{1}H$ NMR of **Probe 3**

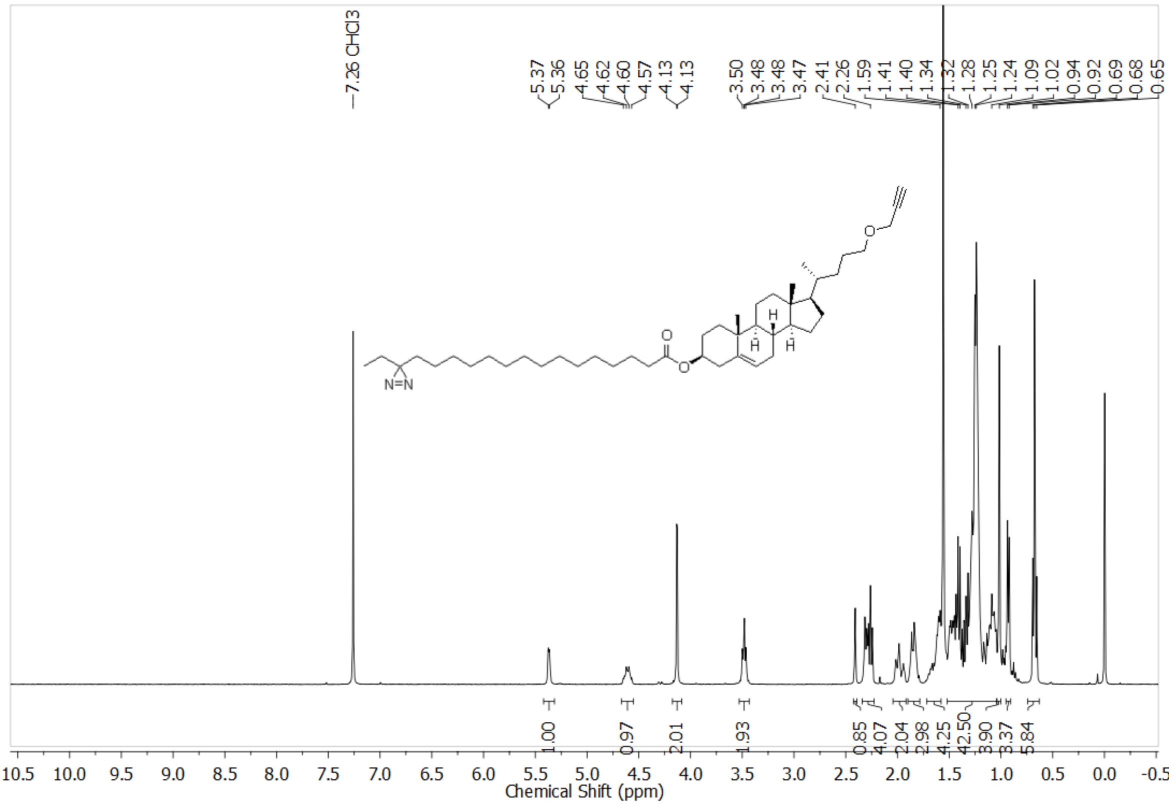

$^{13}C$ NMR of **Probe 3**

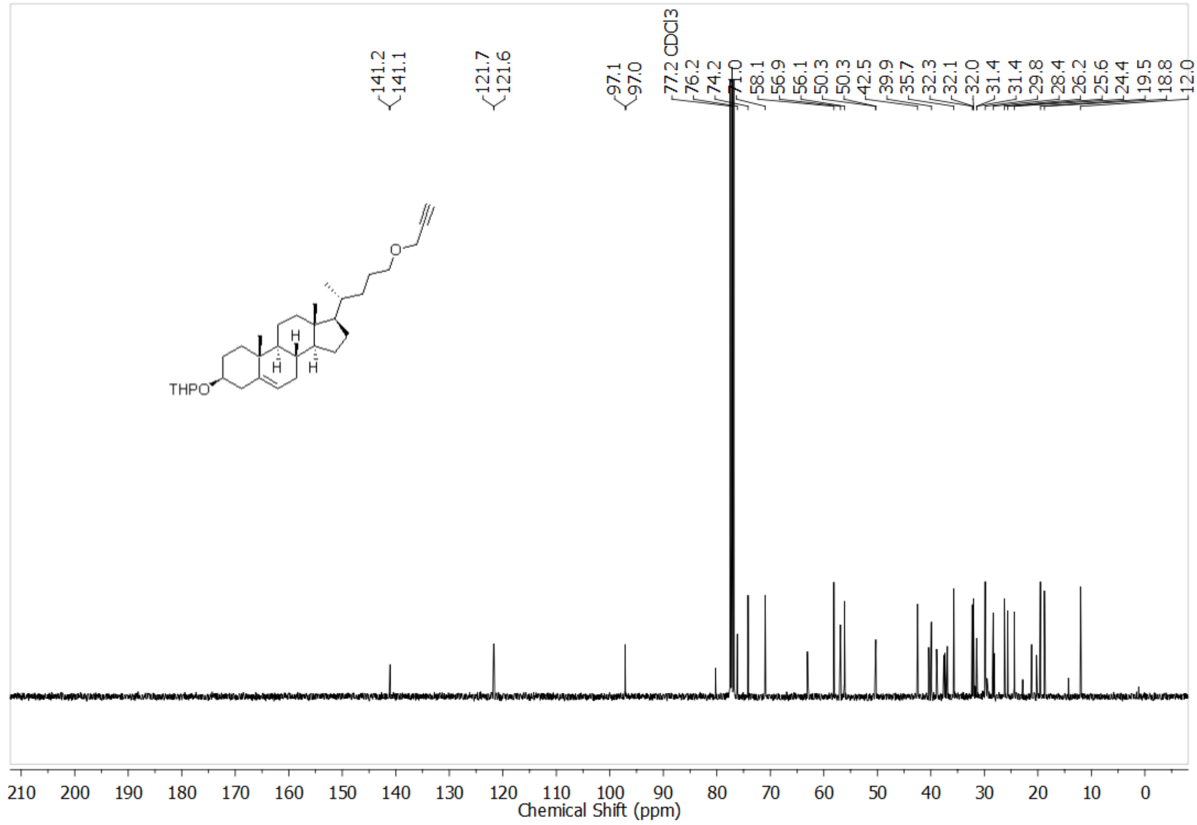

FTIR spectra of **Probe 3**

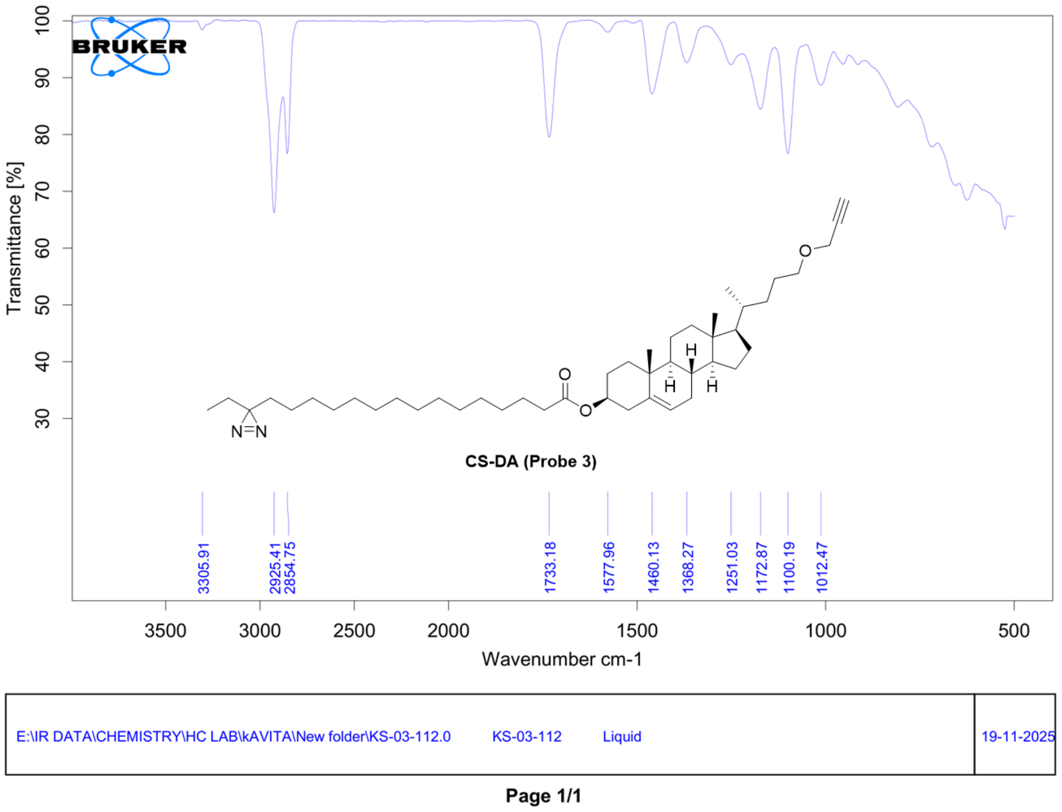


HRMS of **Probe 3**

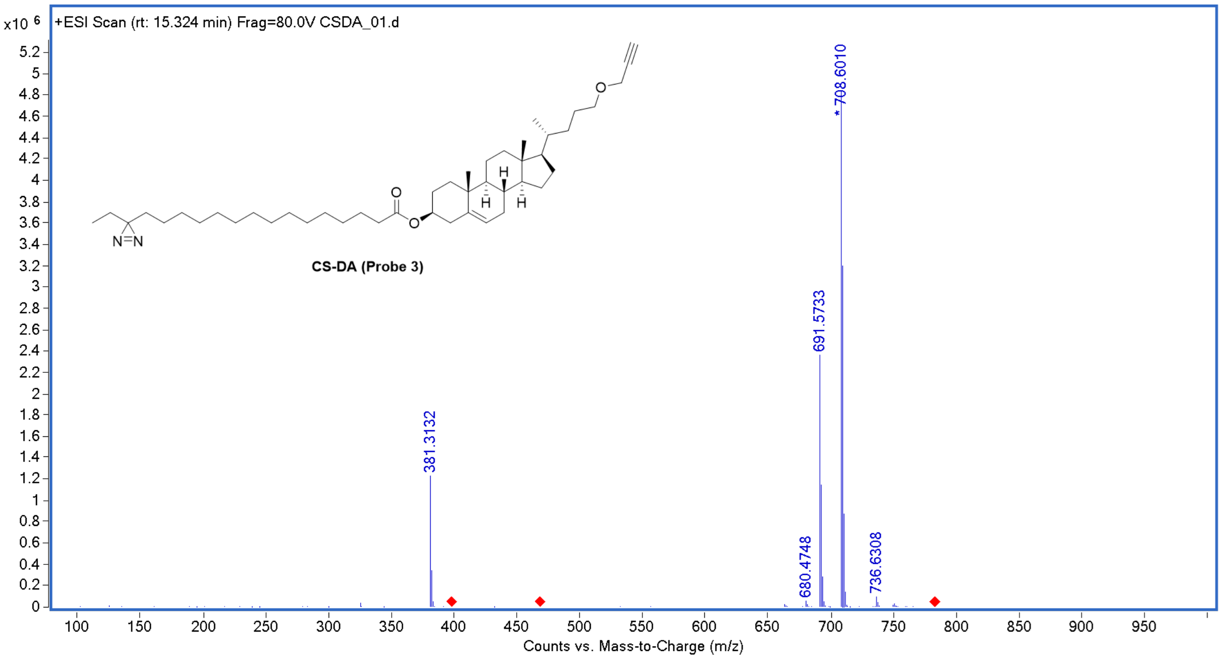

[1]H NMR of **Probe 4**

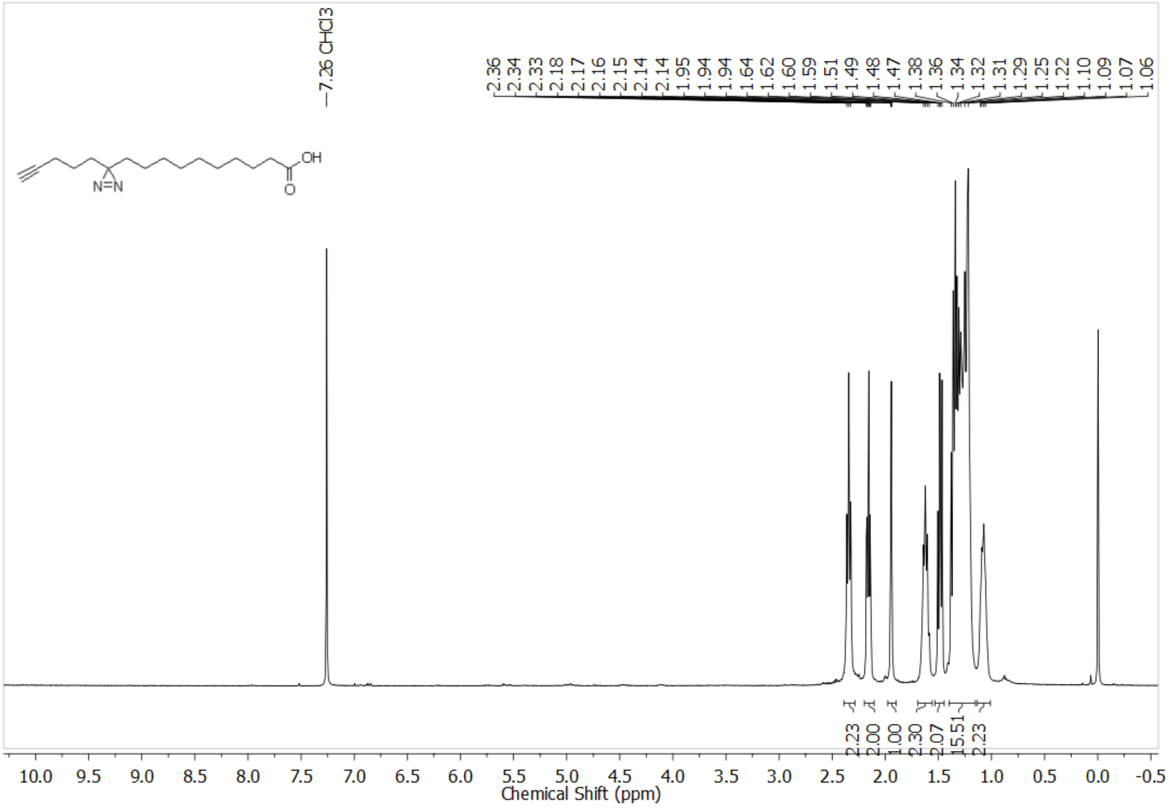


HRMS of **Probe 4**

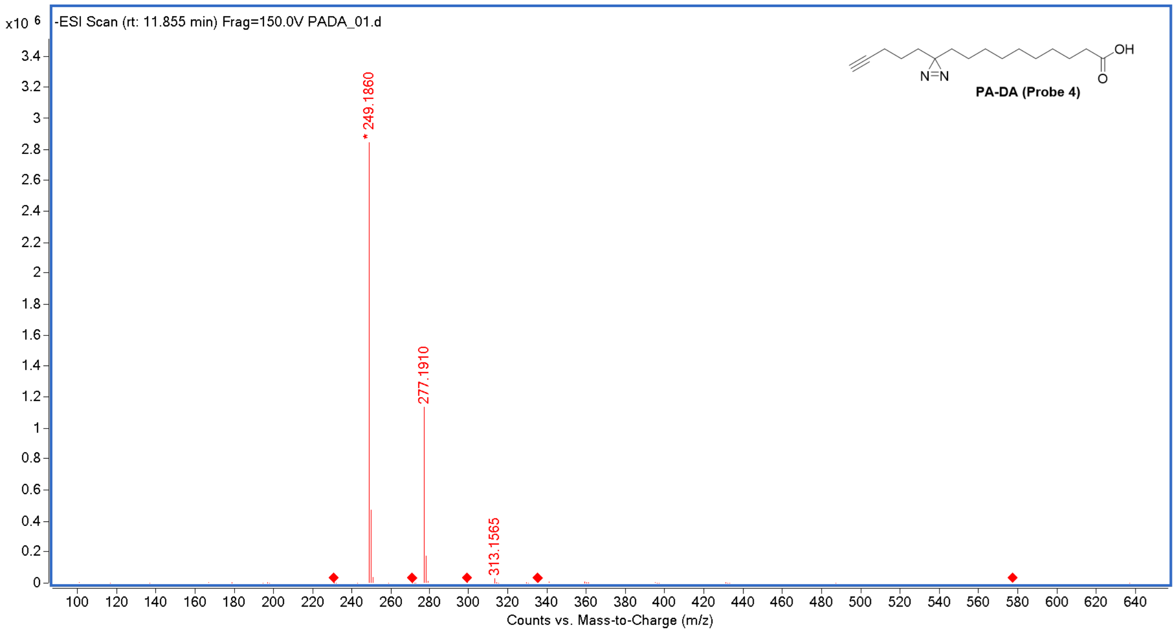

$^{1}$H NMR of **Probe 5**

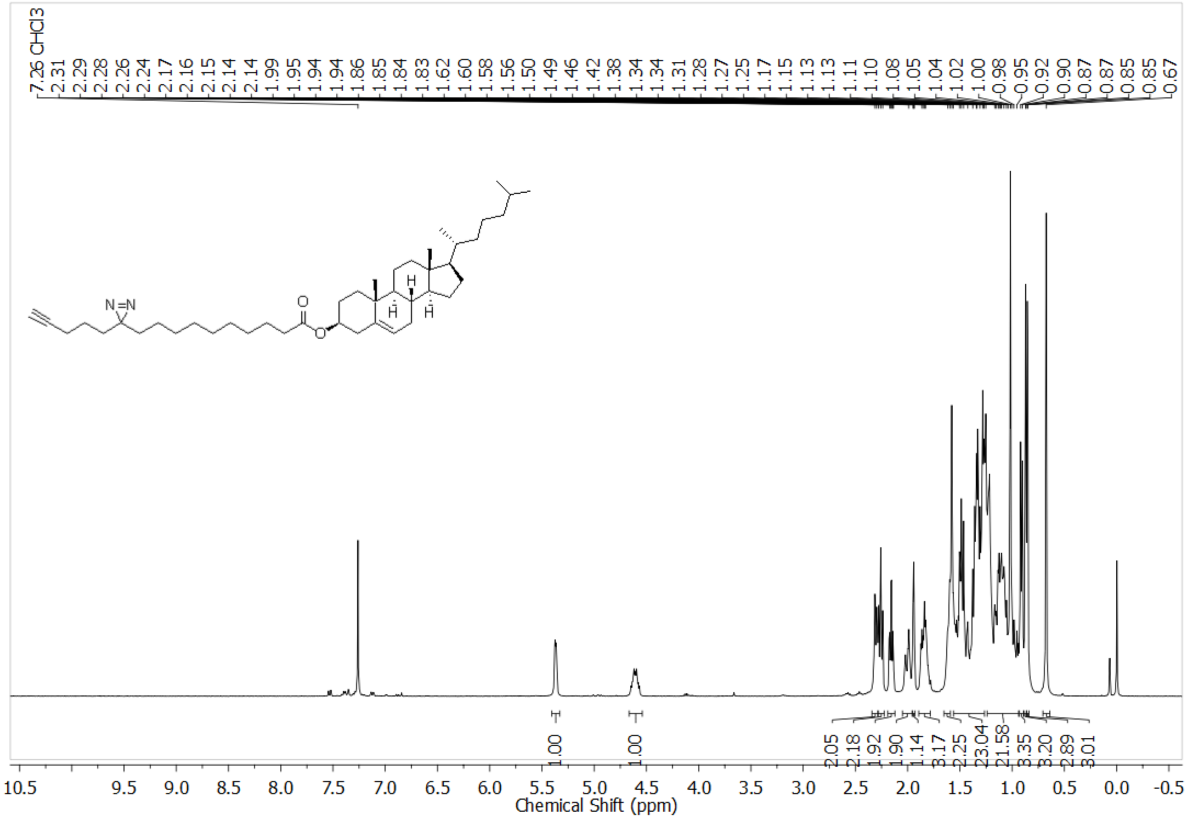


$^{13}$C NMR of **Probe 5**

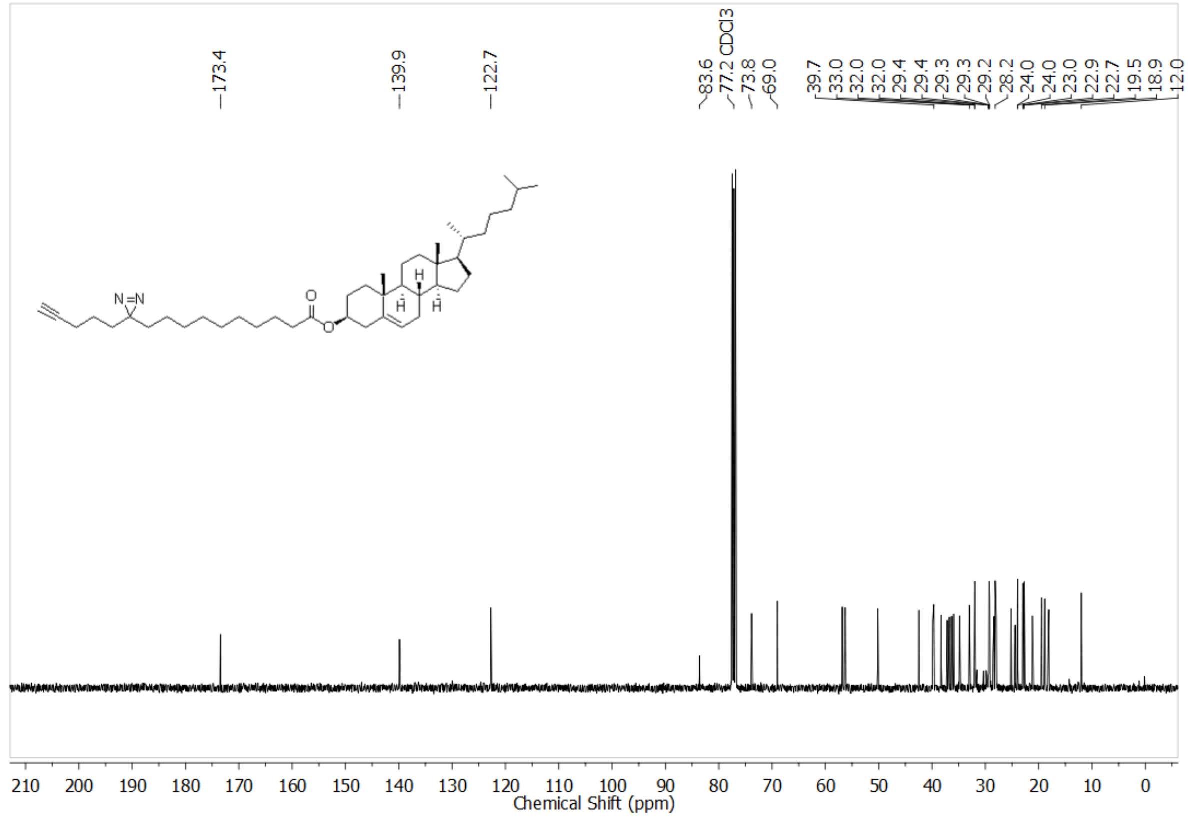

FTIR spectra of **Probe 5**

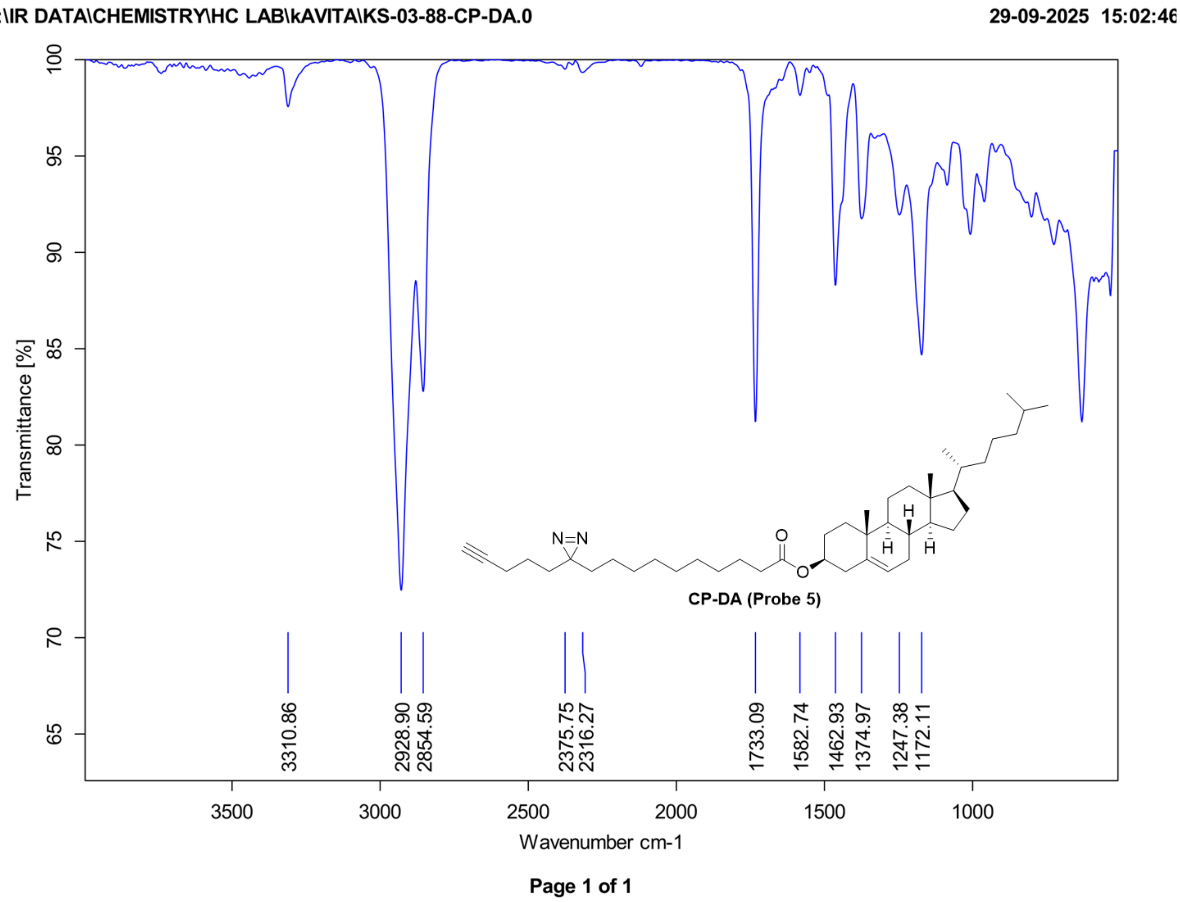


HRMS of **Probe 5**

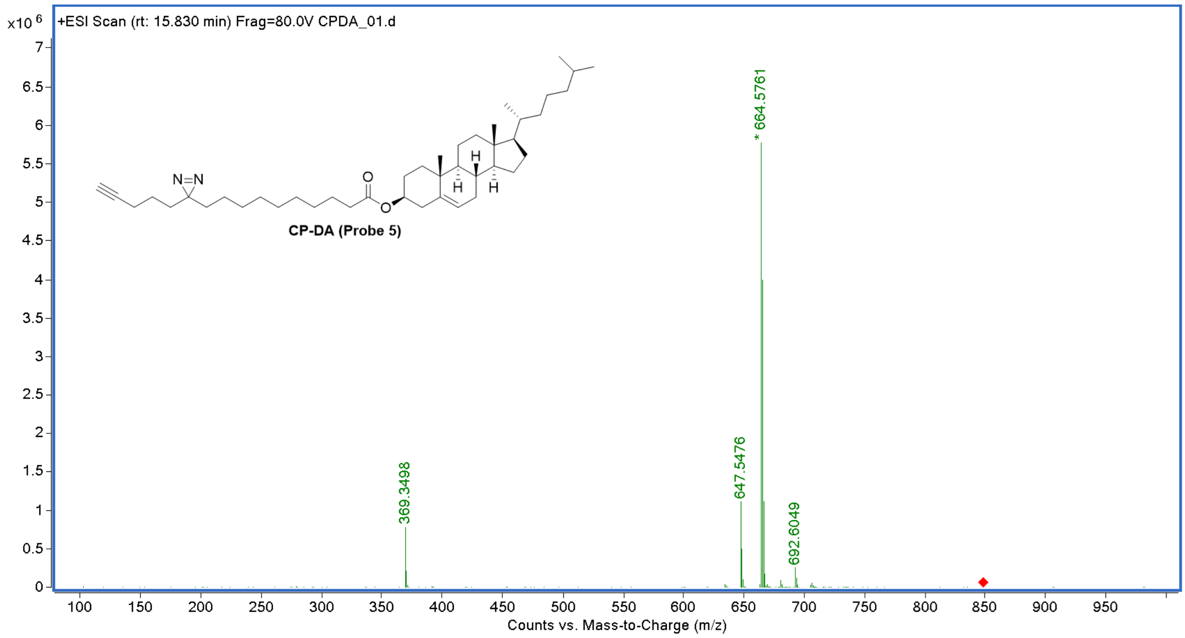

$^{1}$H NMR of **Probe 6**

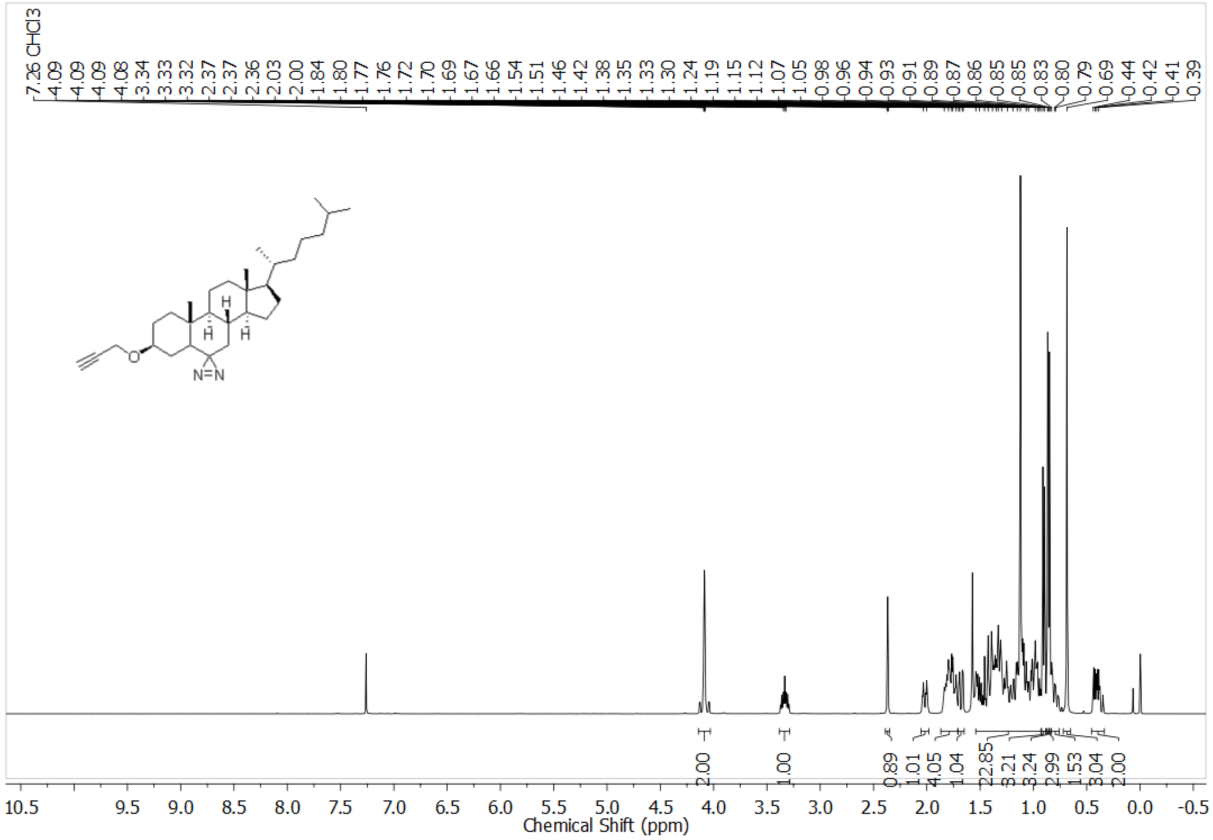

$^{13}$C NMR of **Probe 6**

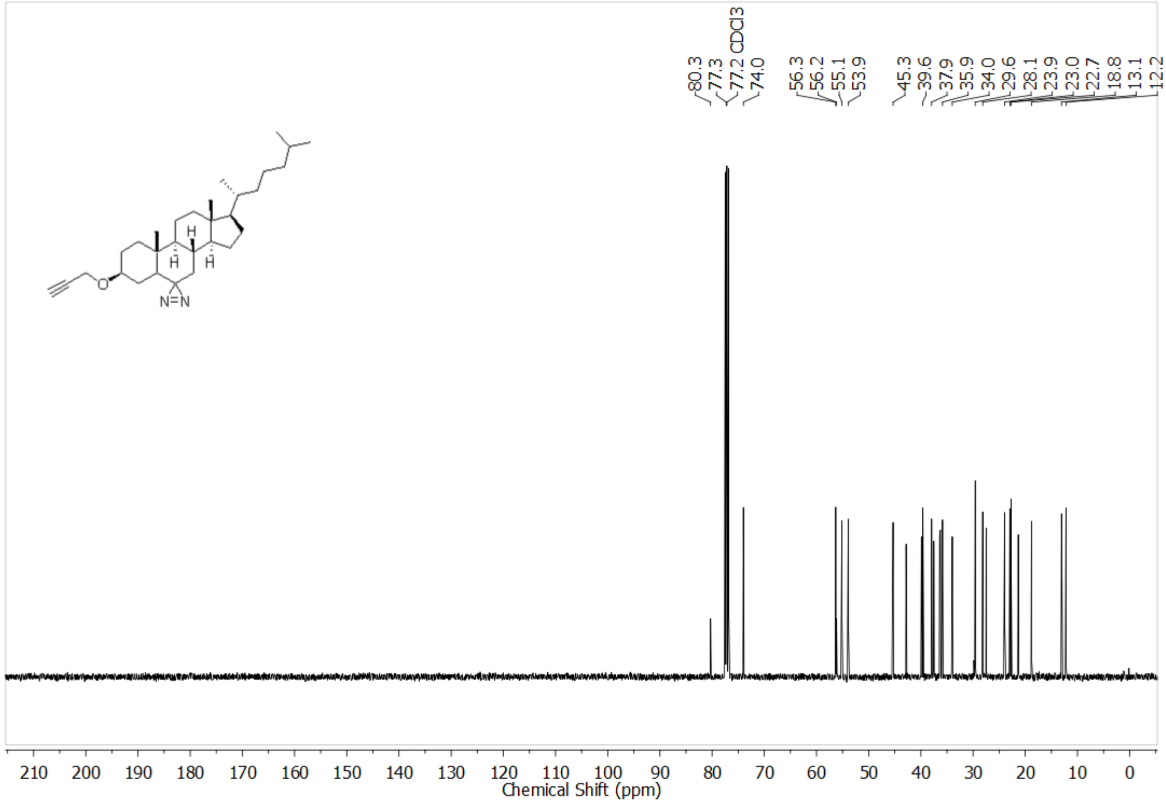

FTIR spectra of **Probe 6**

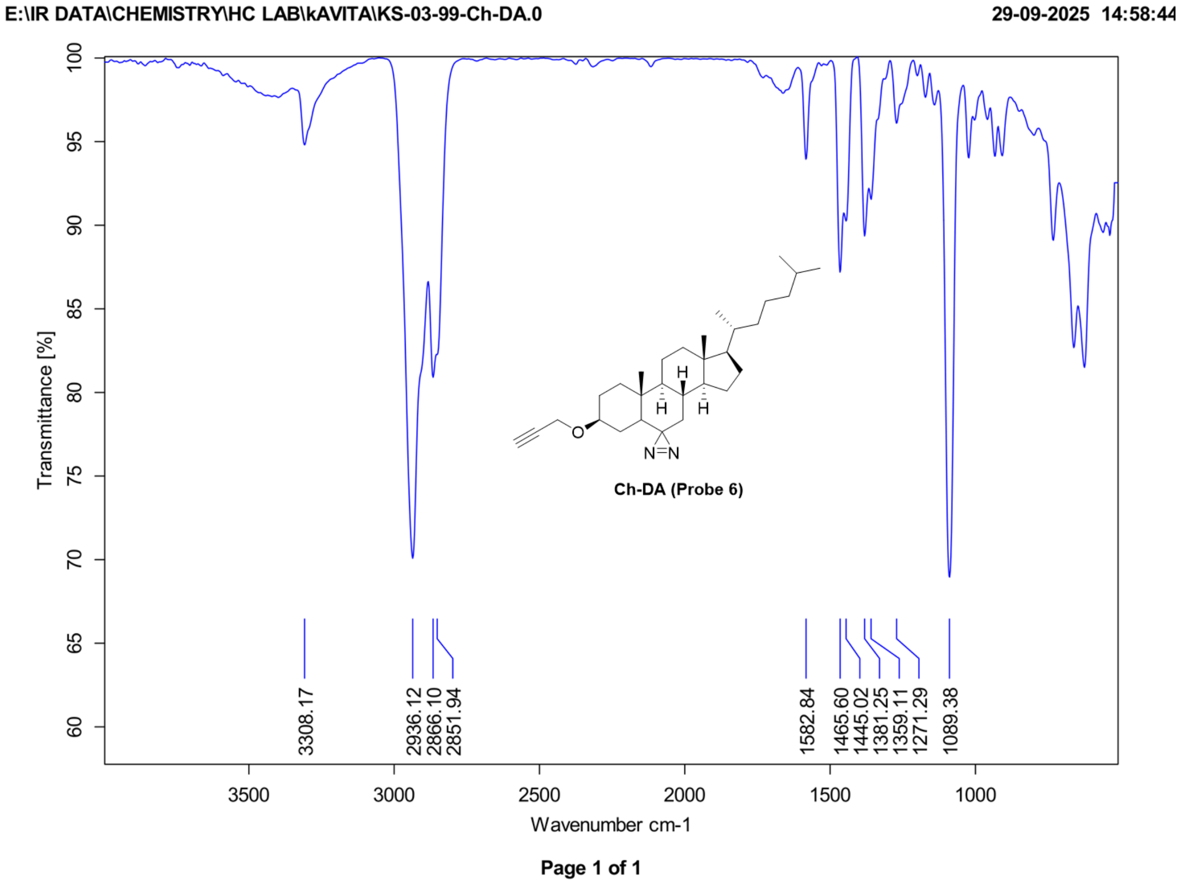


HRMS of **Probe 6**

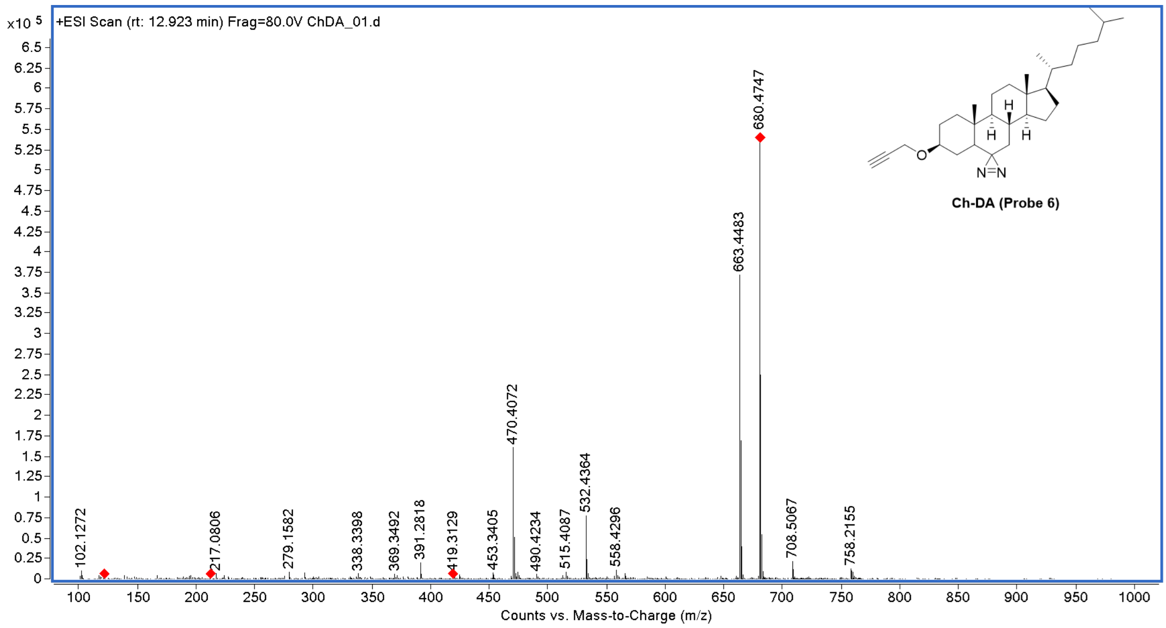

### CHARACTERIZATION SPECTRA OF INTERMEDIATES

$^{1}$H NMR of **1**

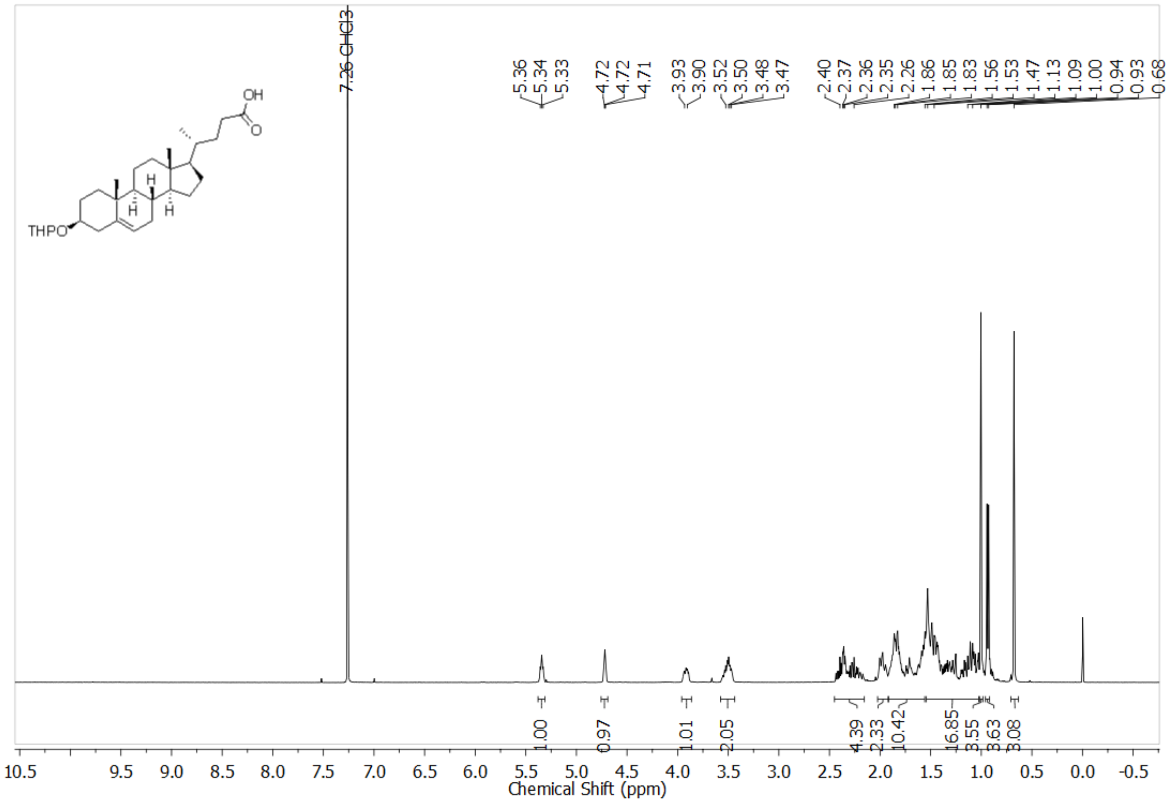


$^{1}$H NMR of **2**

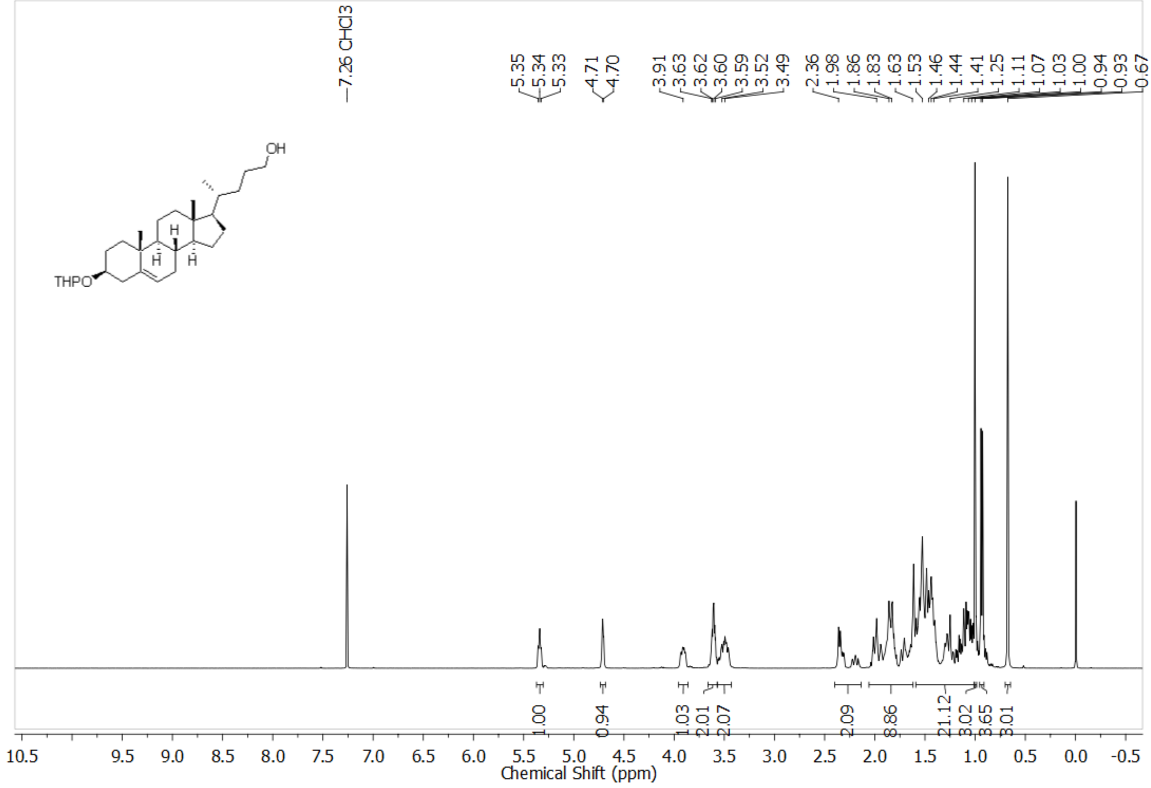

$^{1}$H NMR of **3**

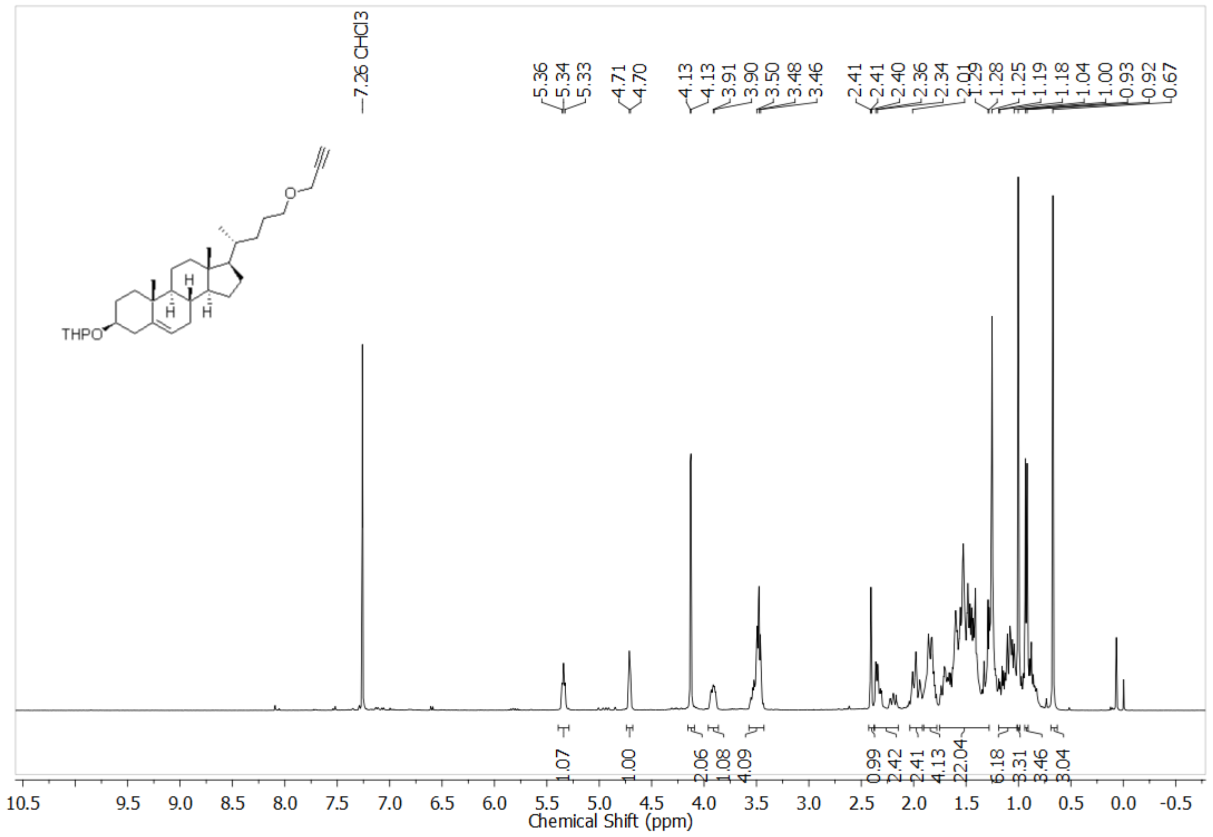


$^{13}$C NMR of **3**

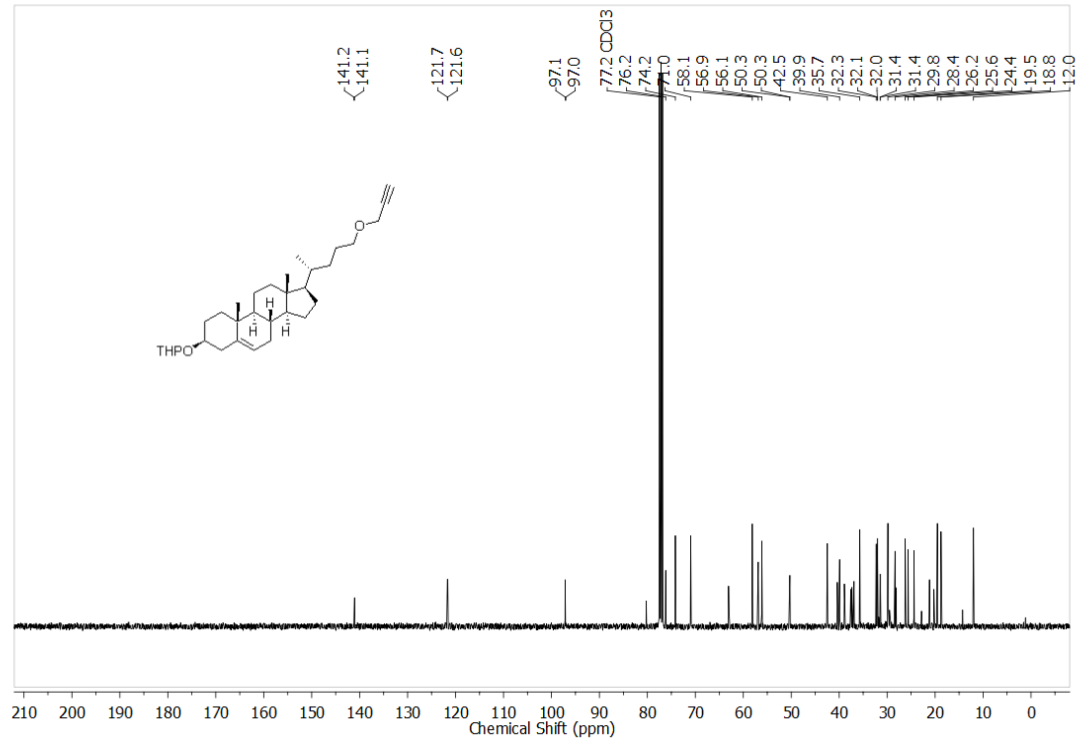

FTIR spectra of **3**

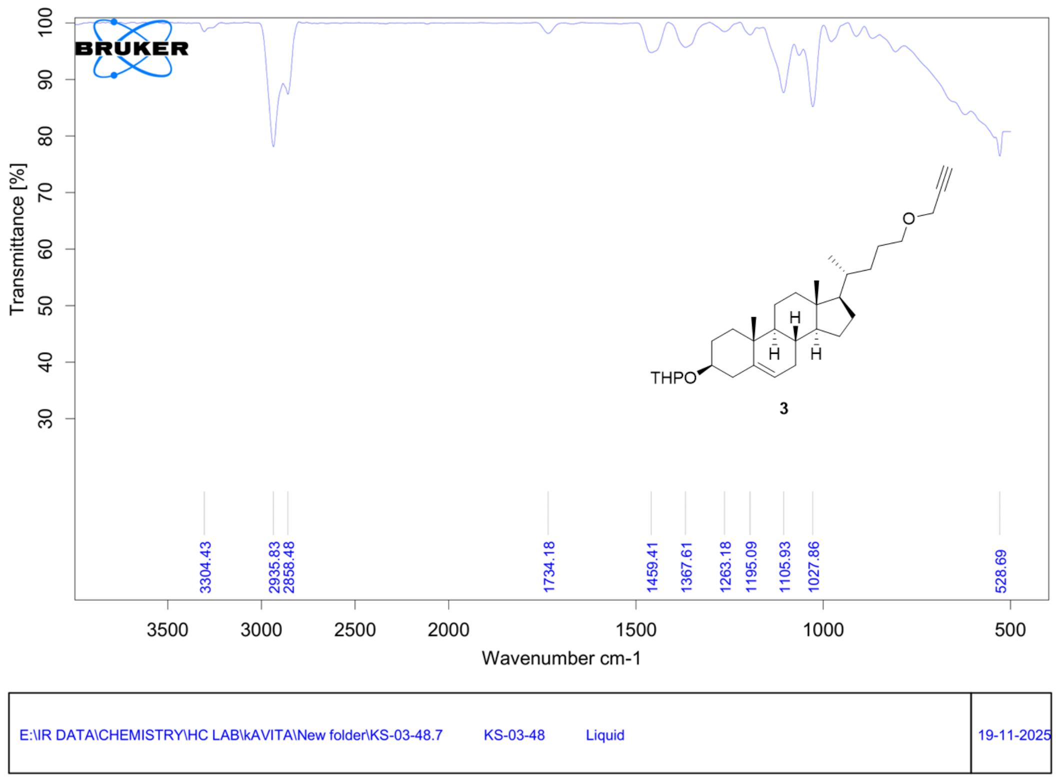


[1]H NMR of **4**

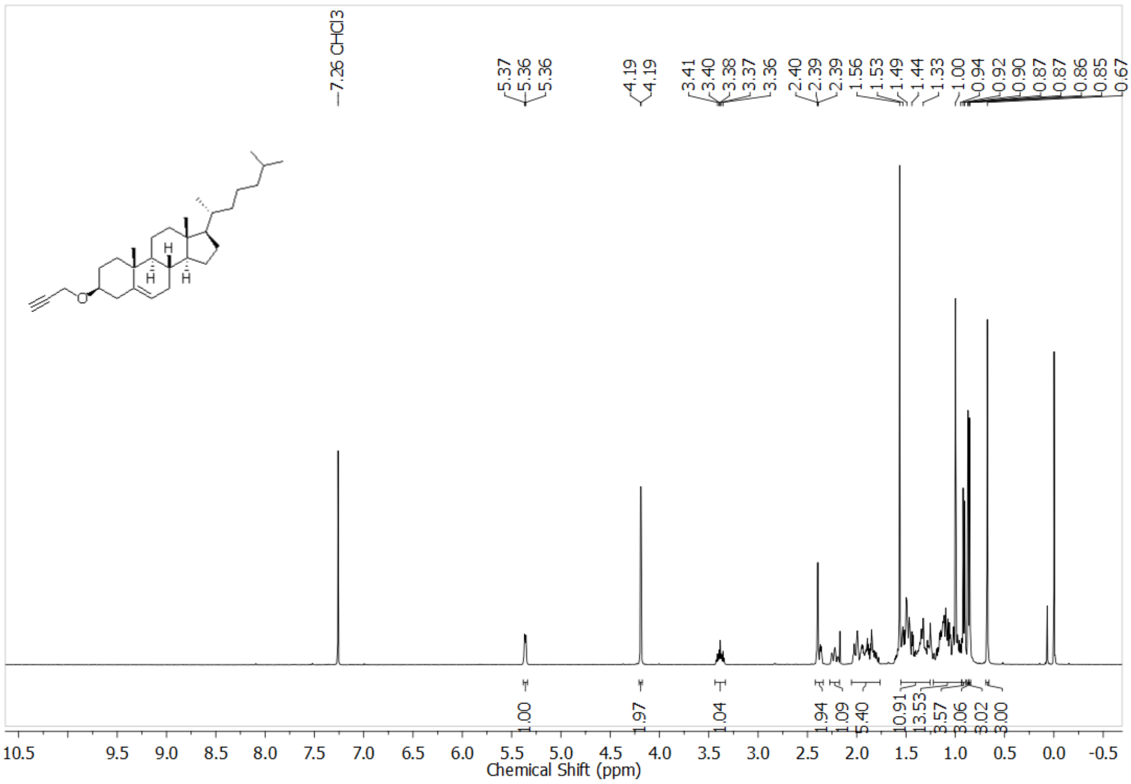

$^{1}$H NMR of **5**

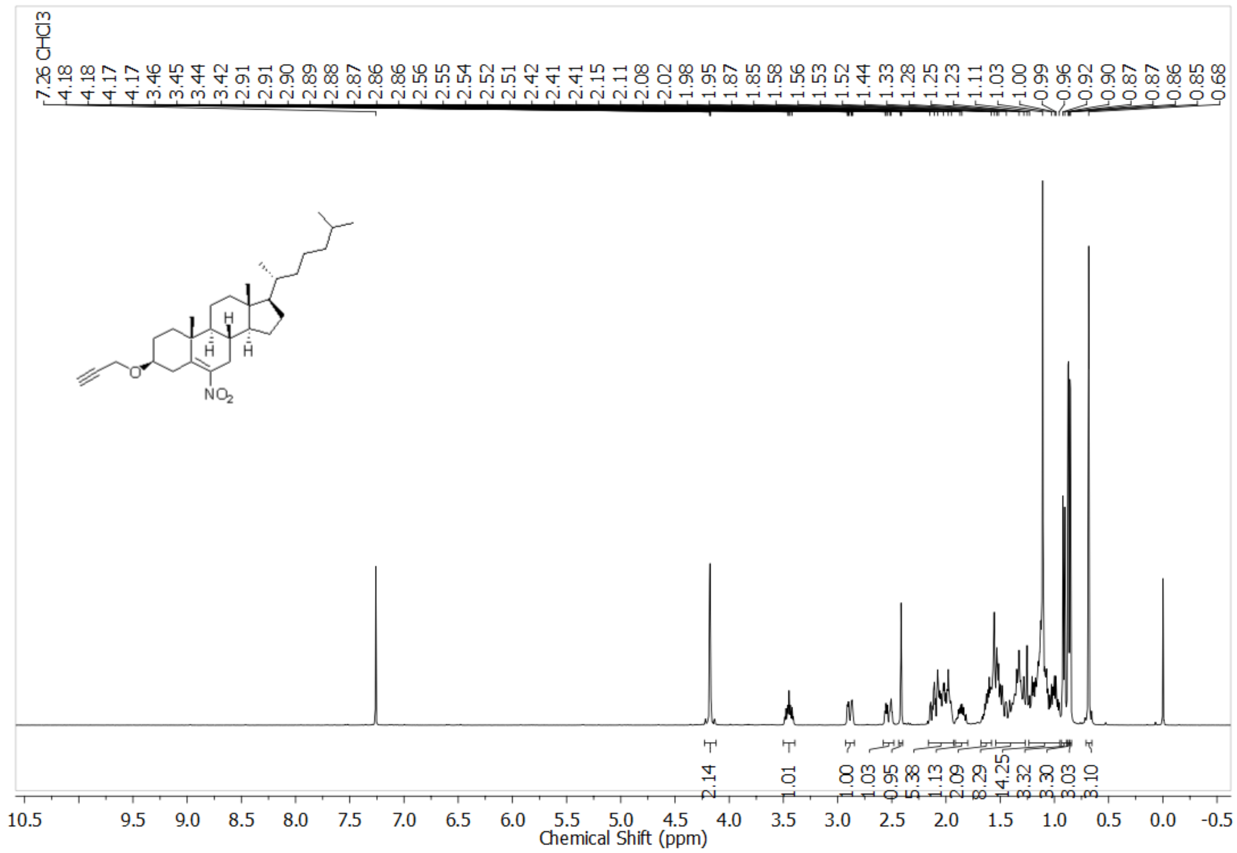

$^{13}$C NMR of **5**

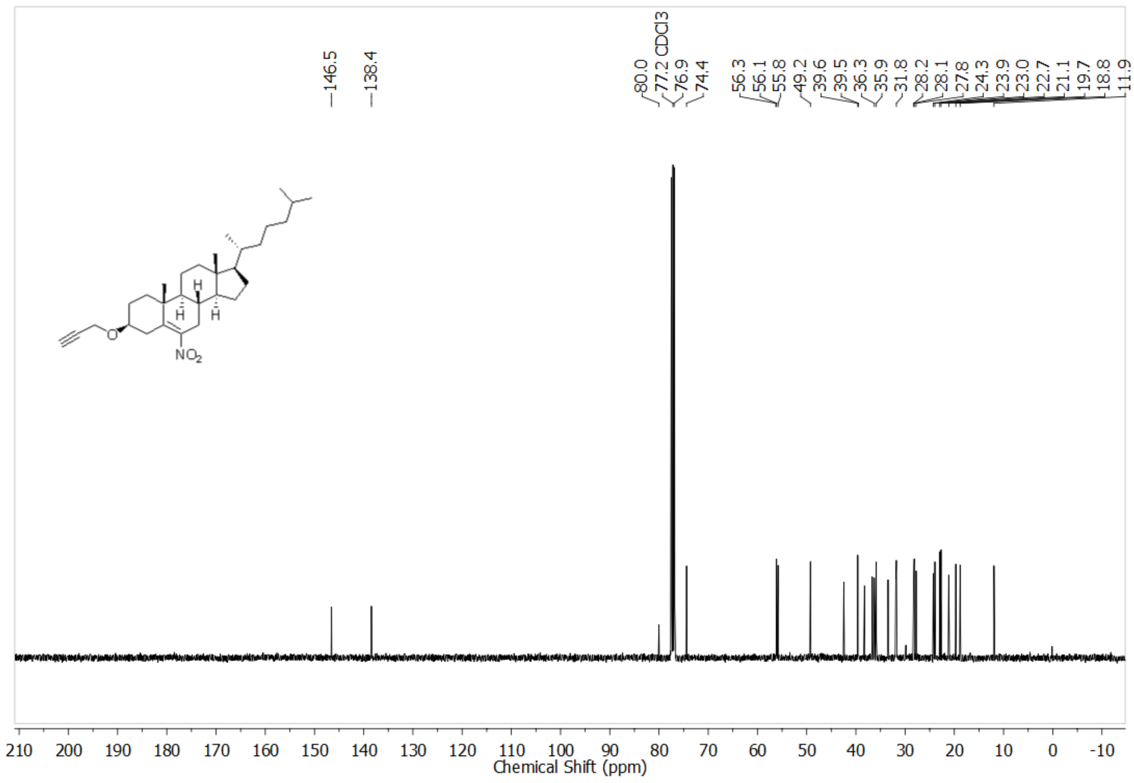

FTIR spectra of **5**

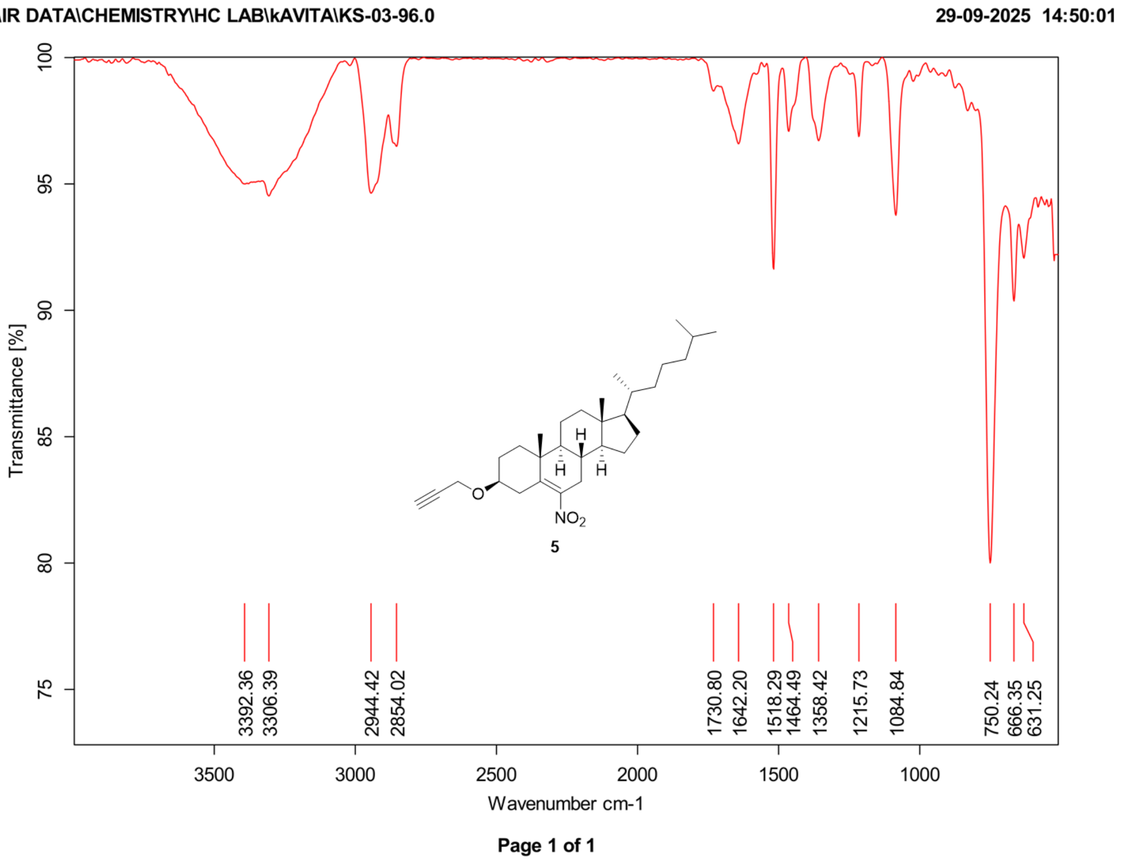


HRMS of **5**

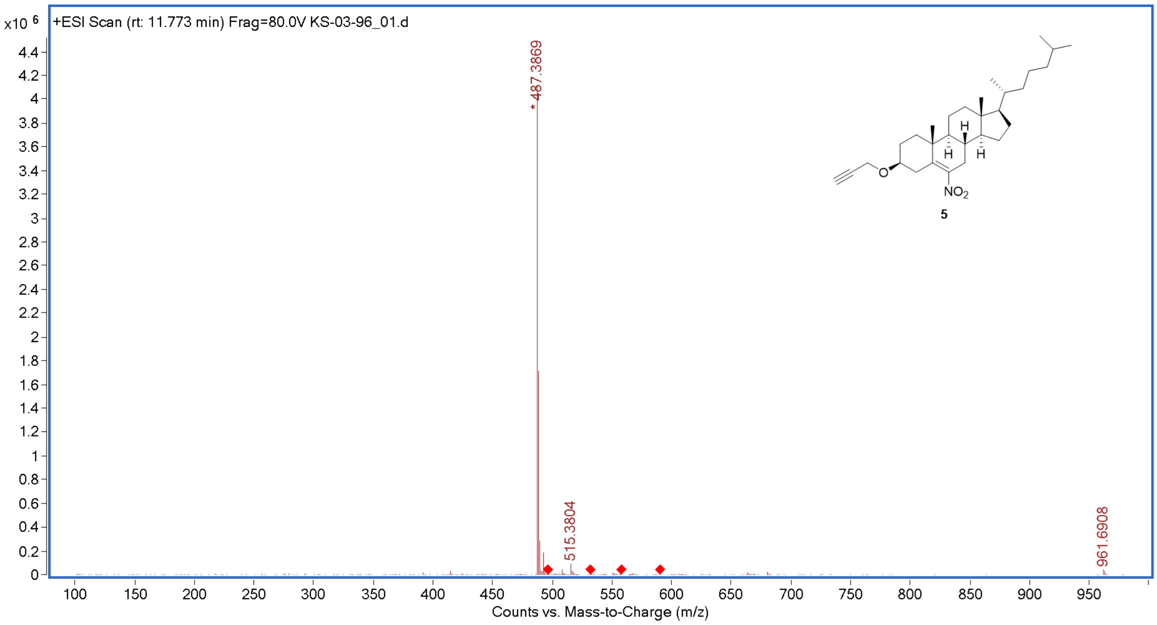

<sup></sup>[1]H NMR of **6**

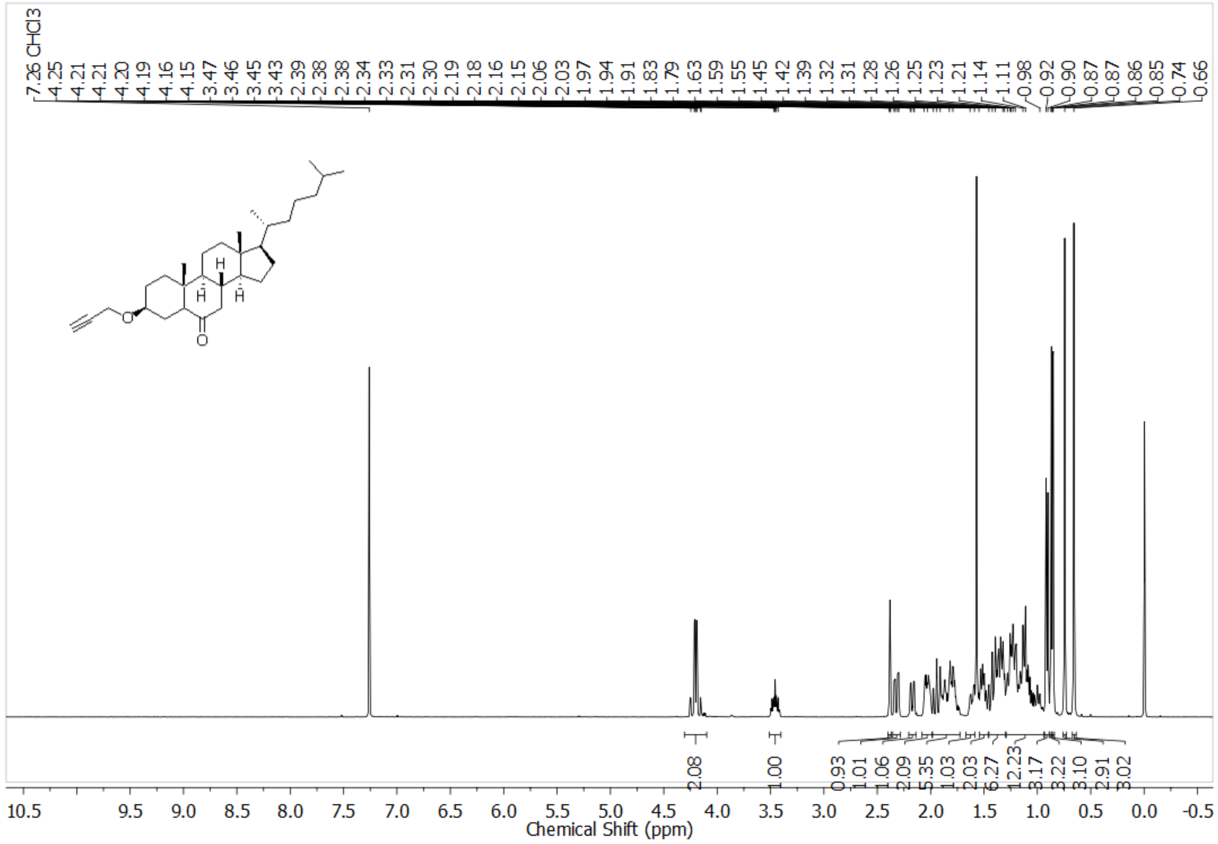

[13]C NMR of **6**

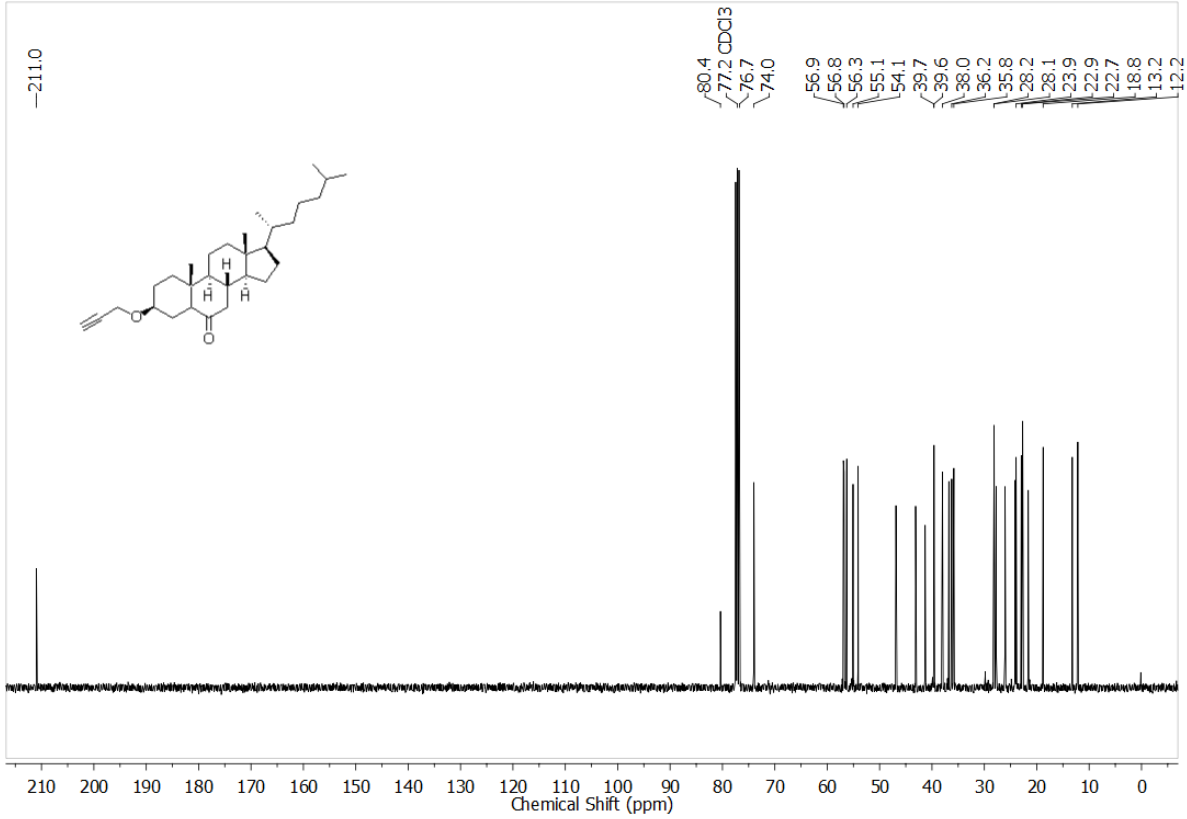

FTIR spectra of **6**

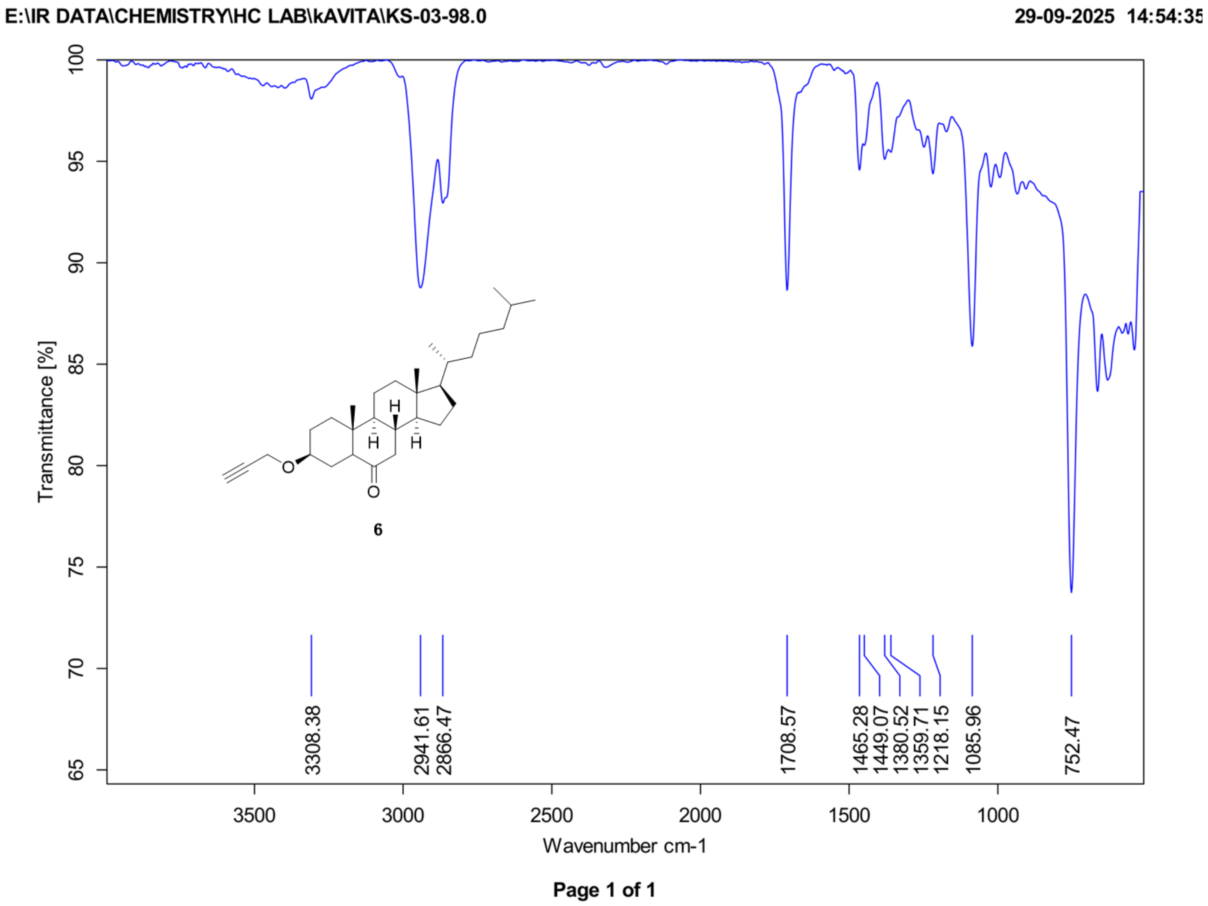


HRMS of **6**

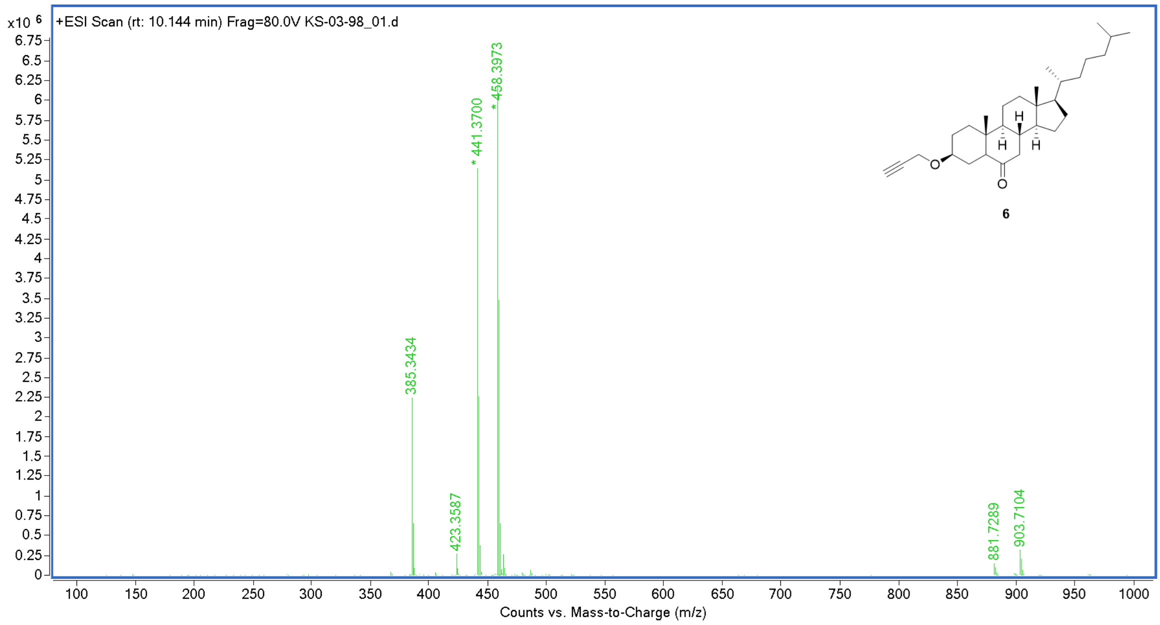

$^{1}$H NMR of **7**

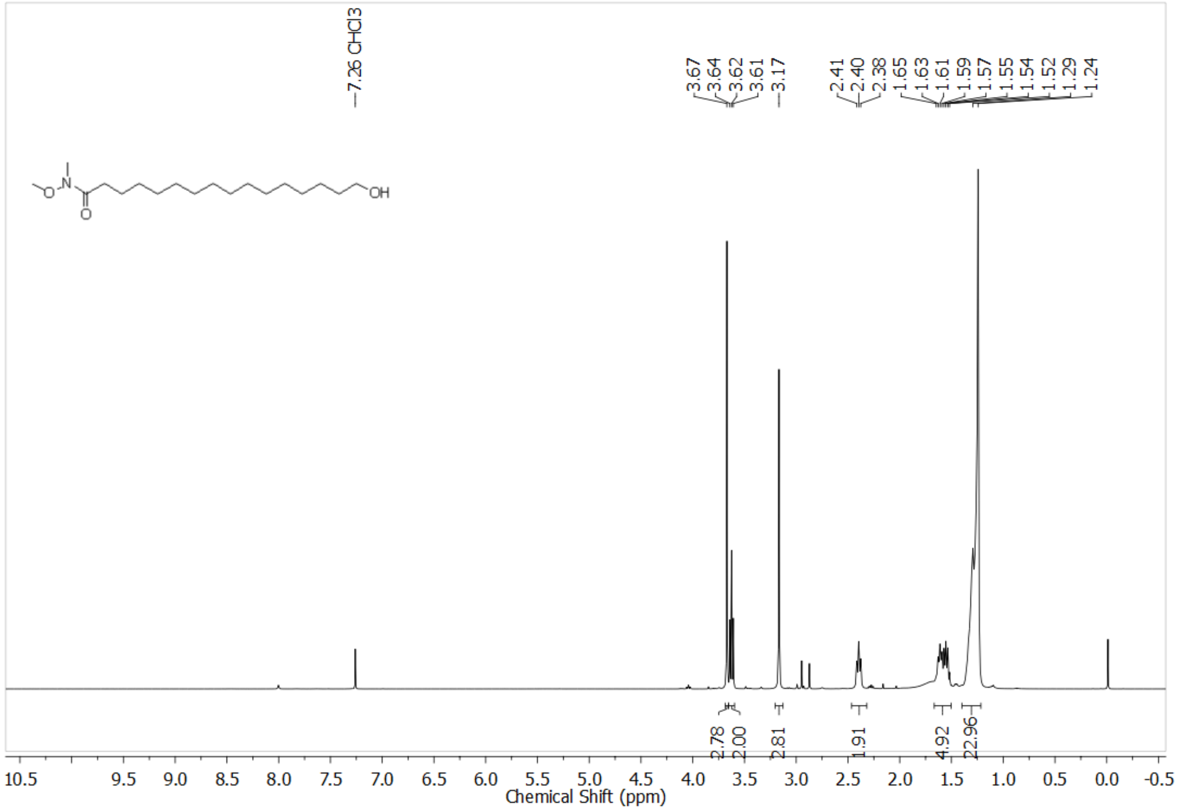


$^{1}$H NMR of **8**

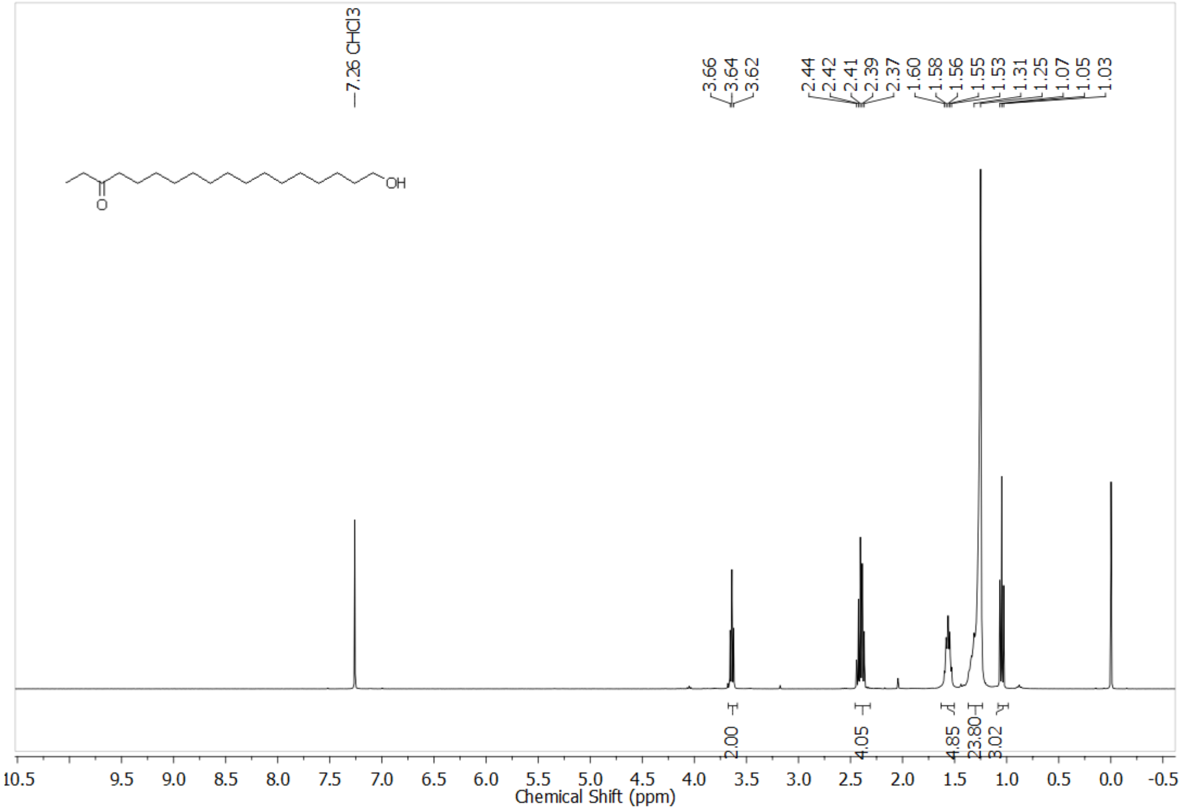

$^1$H NMR of **9**

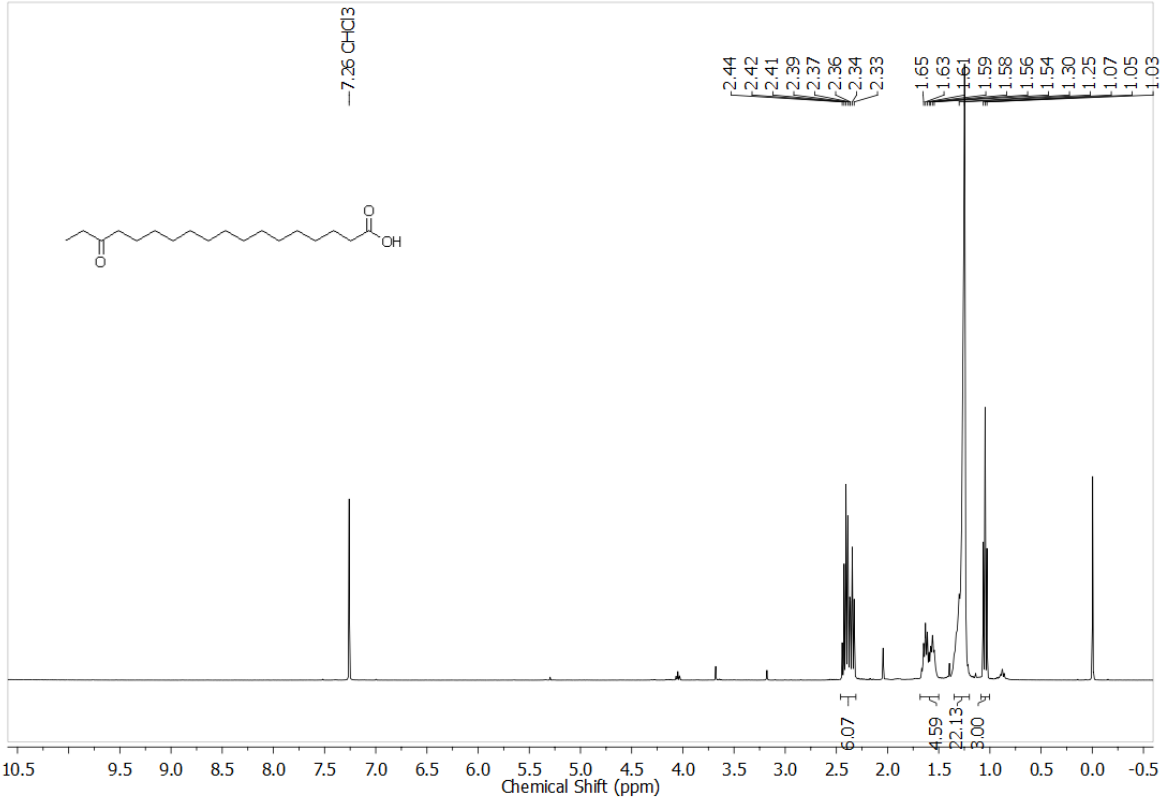


$^1$H NMR of **10**

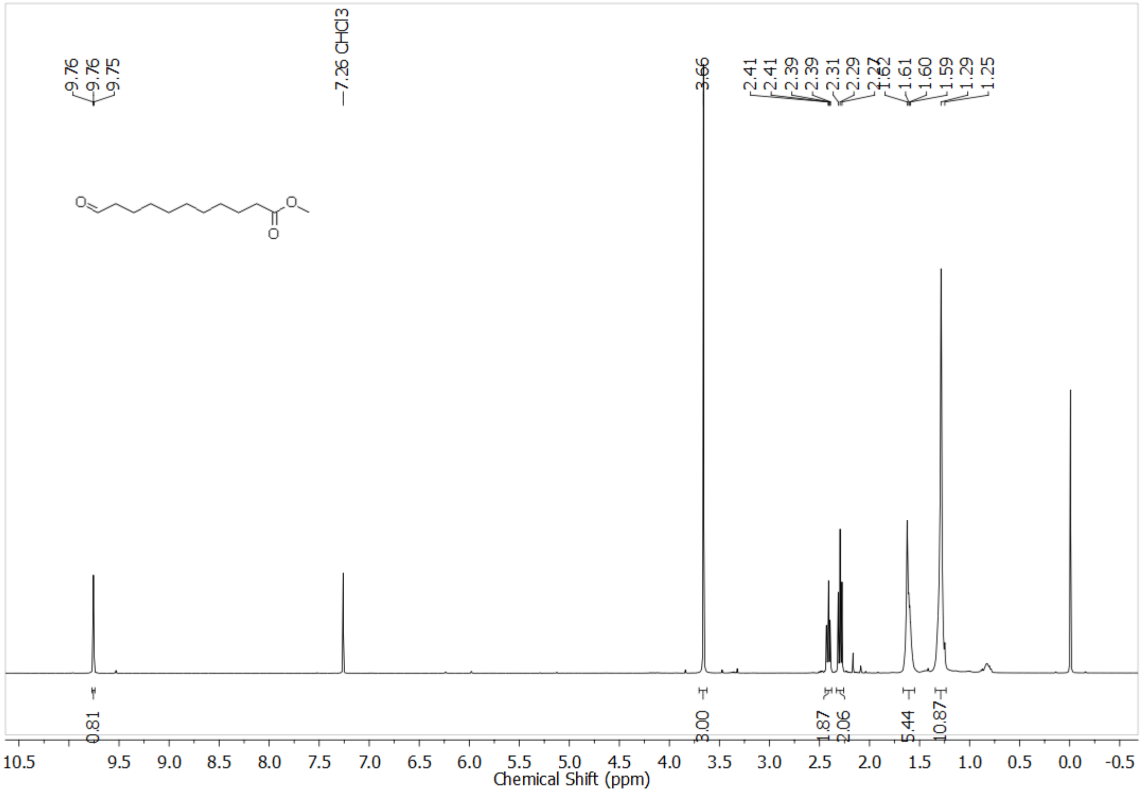

[1]H NMR of **11**

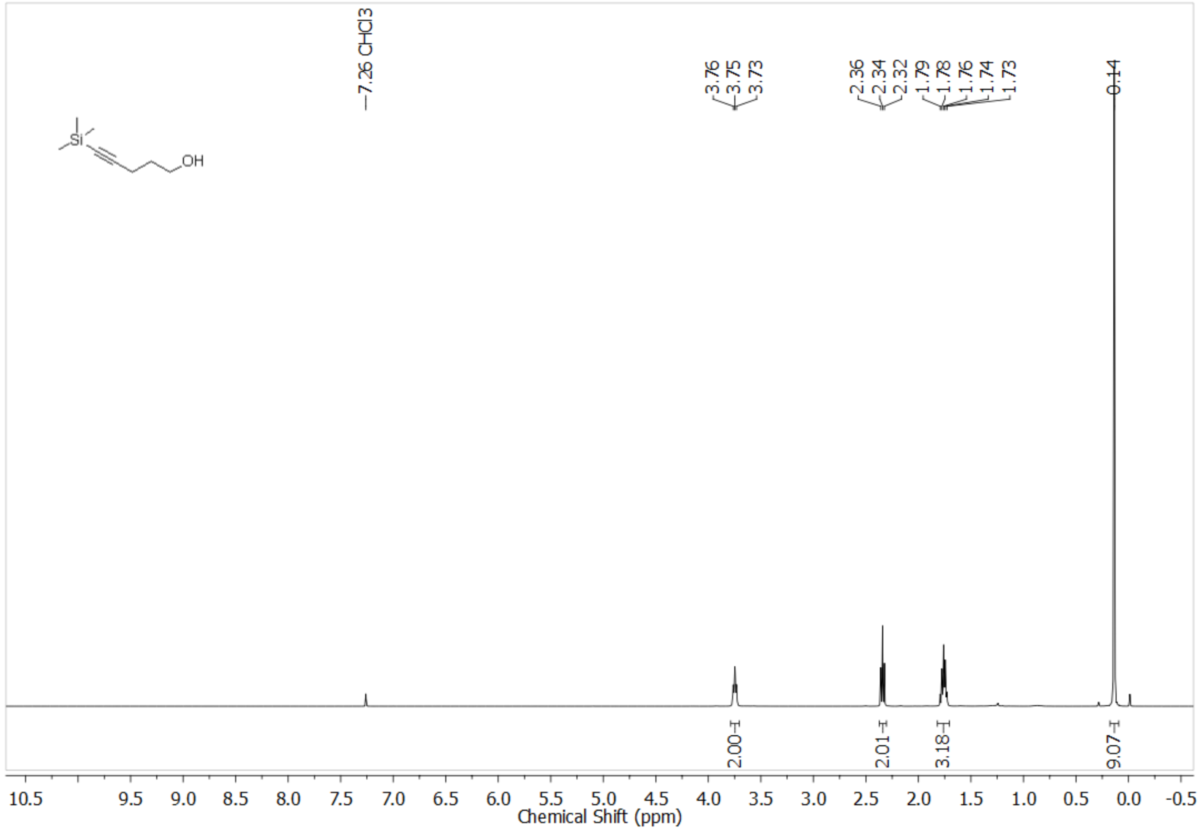


[1]H NMR of **12**

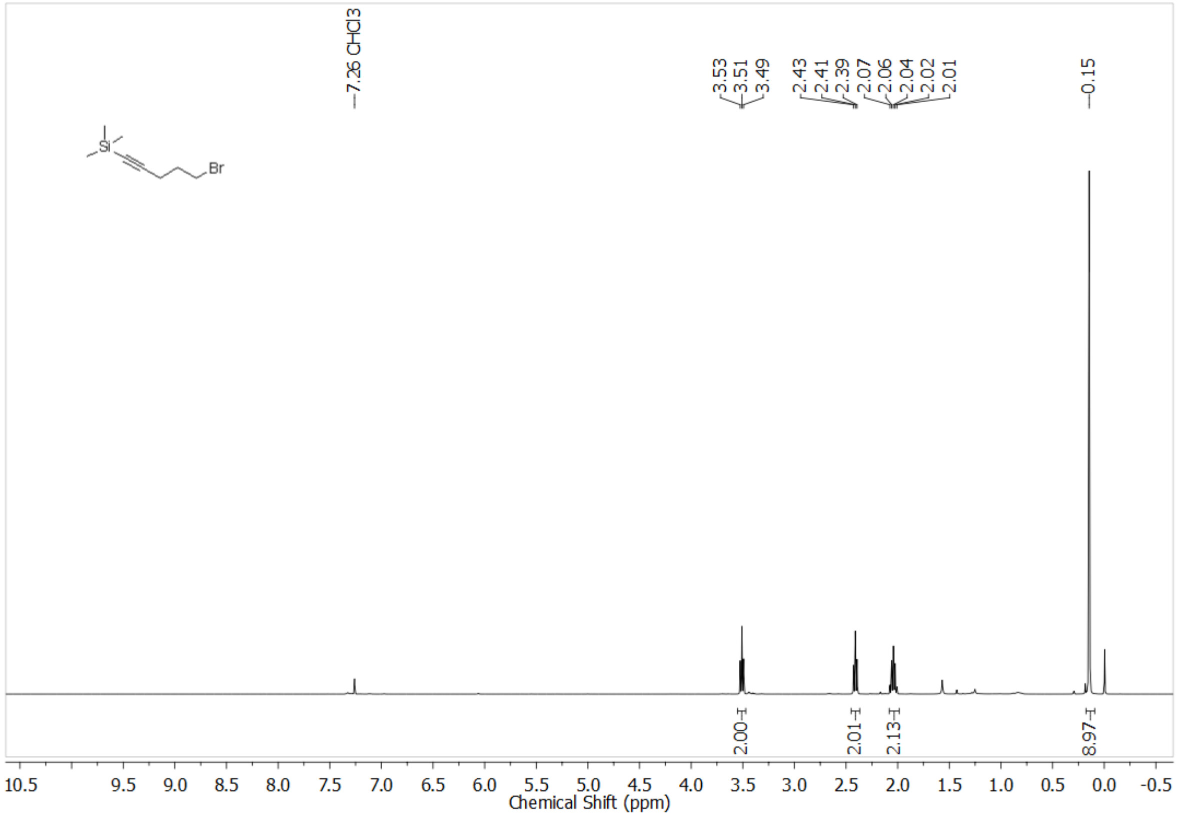

$^{1}$H NMR of **14**

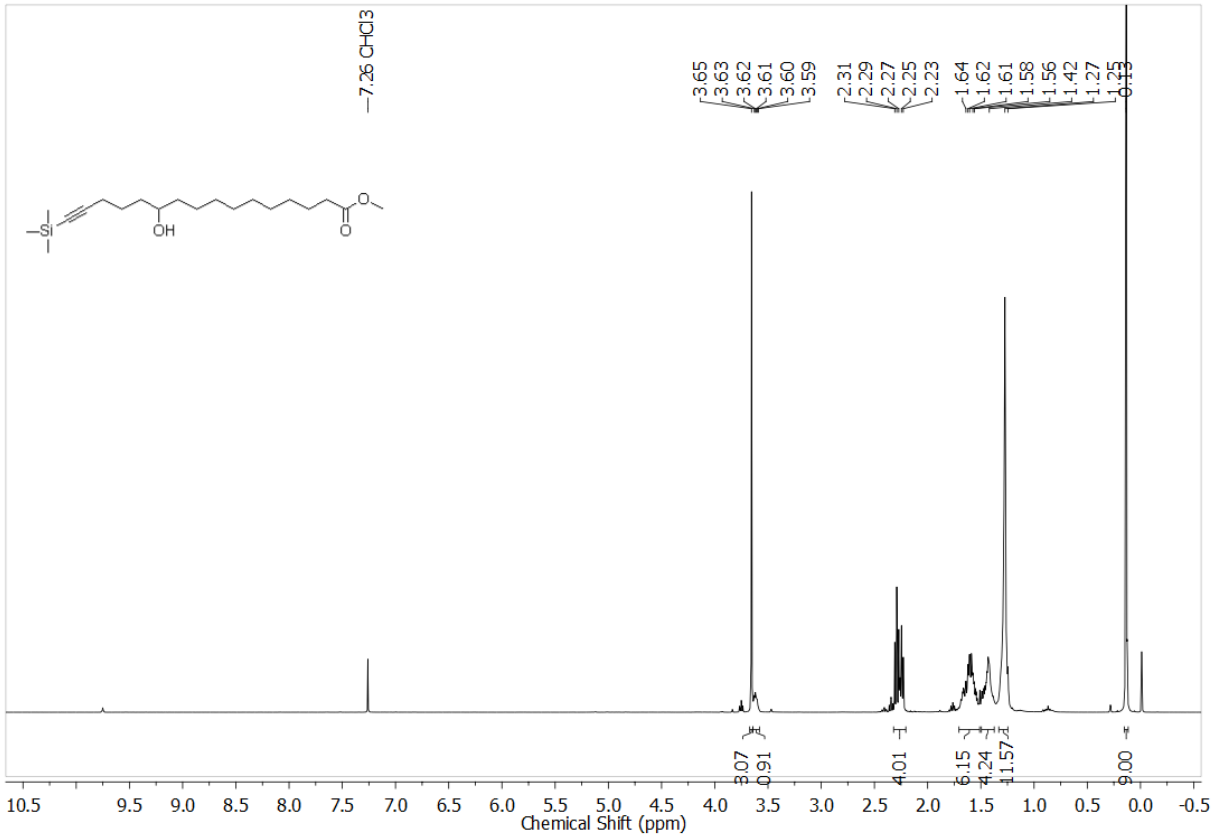


$^{1}$H NMR of **15**

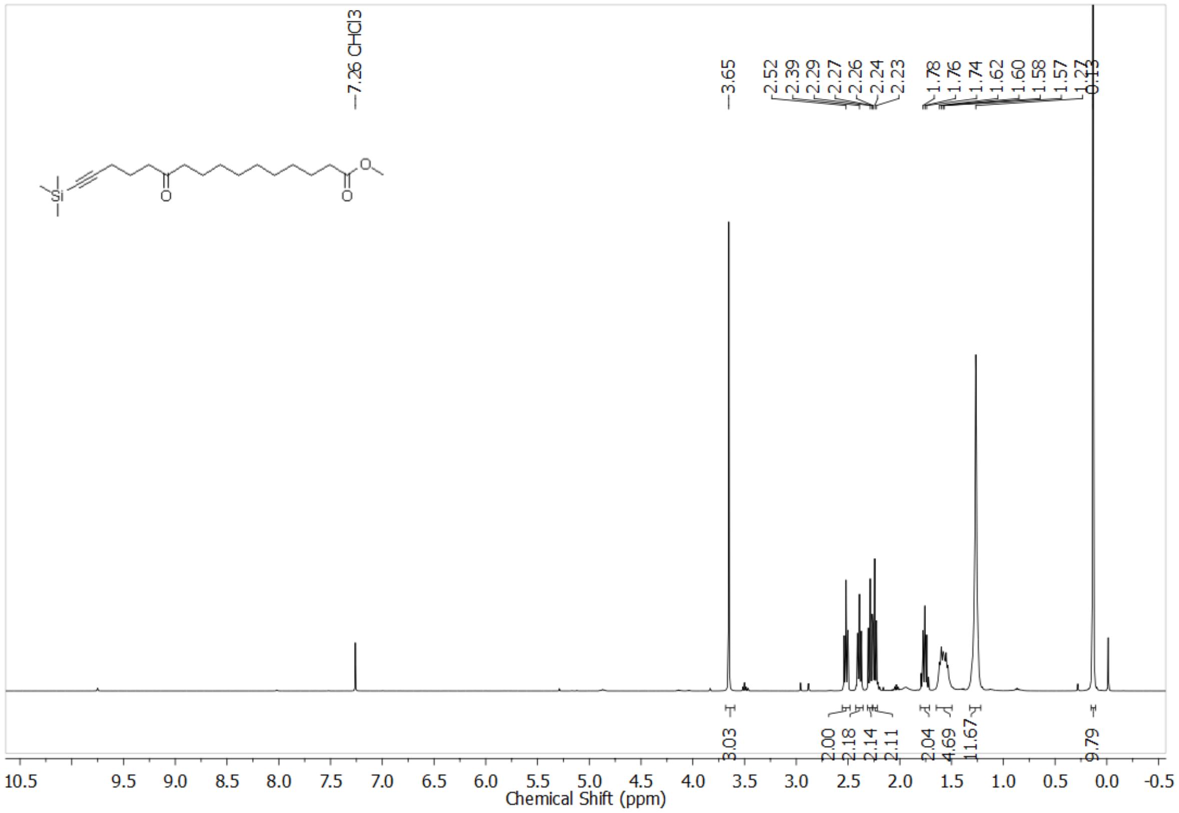

[1]H NMR of **16**

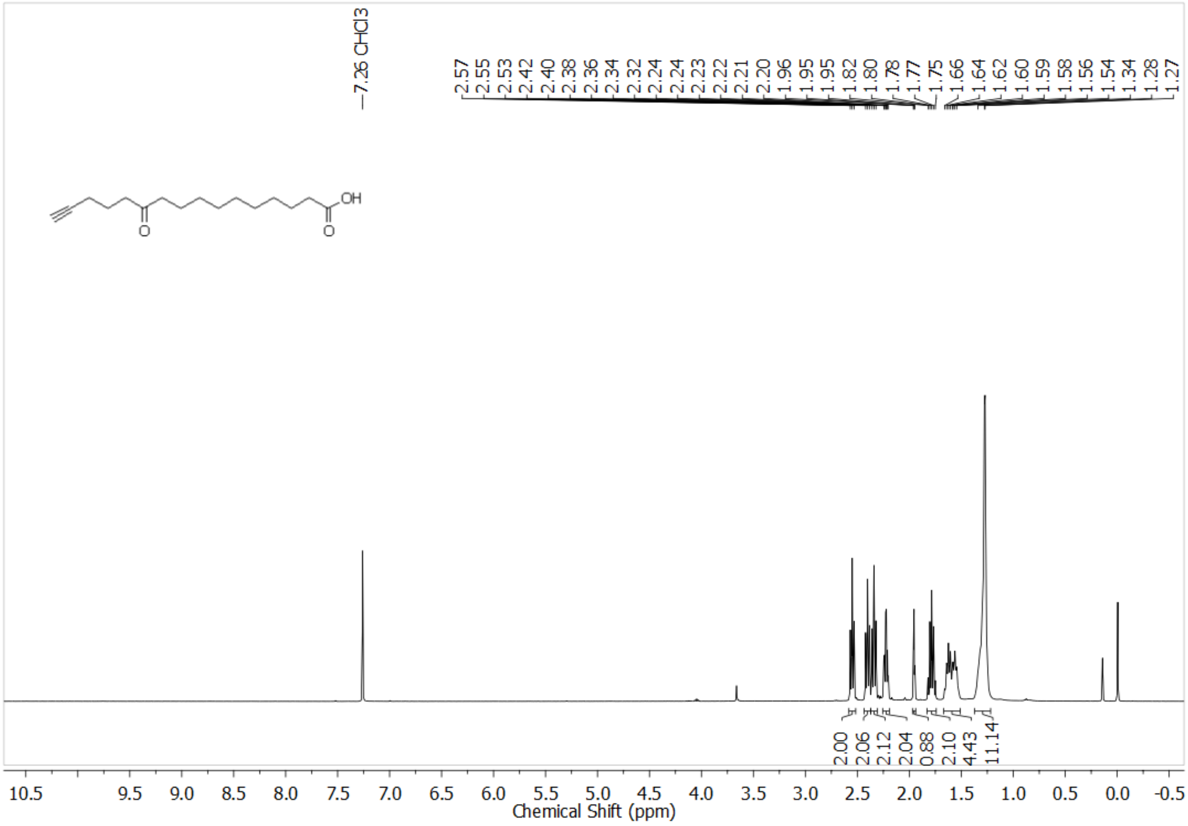
7.26 CHCl3
2.57
2.55
2.53
2.42
2.40
2.38
2.36
2.34
2.32
2.24
2.24
2.23
2.22
2.21
2.20
1.96
1.95
1.95
1.82
1.80
1.78
1.77
1.75
1.66
1.64
1.62
1.60
1.59
1.58
1.56
1.54
1.34
1.28
1.27
OH
O
O
2.00
2.06
2.12
2.04
0.88
2.10
4.43
11.14
10.5
9.5
9.0
8.5
8.0
7.5
7.0
6.5
6.0
5.5
5.0
4.5
4.0
3.5
3.0
2.5
2.0
1.5
1.0
0.5
0.0
-0.5
Chemical Shift (ppm)
