## Supplementary material for "Orthogonal Chemical Proteomic Strategies Reveal the Cholesteryl Ester Interactome in Mammalian Cells": SI Material

### INDEX

| Content | Page No: |
| --- | --- |
| Supplementary Schemes 1 – 2 | 3 – 5 |
| Supplementary Figures 1 – 14 | 9 – 19 |
| Loading control gels of main figures | 20 – 21 |
| References | 22 |

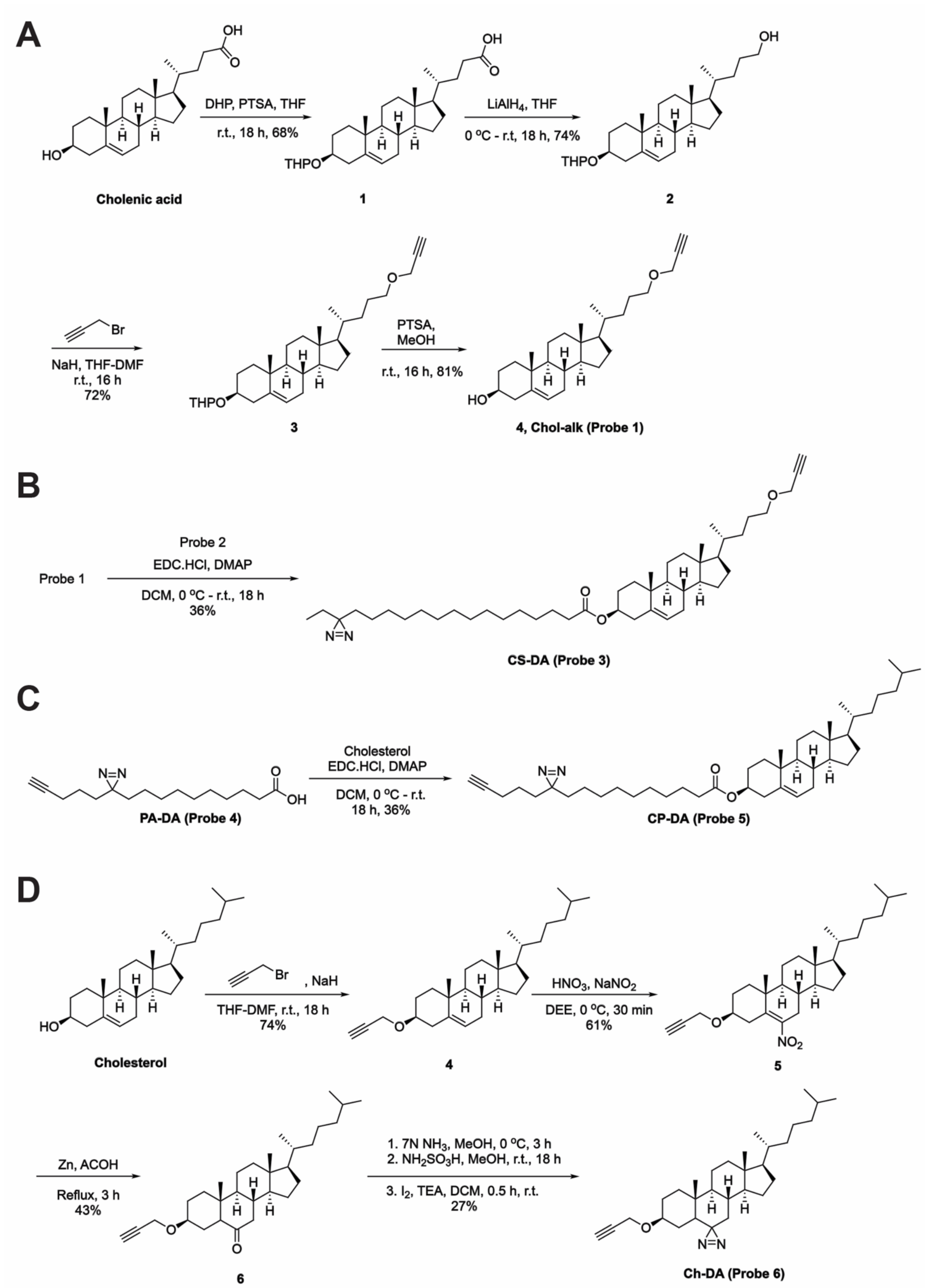


**Scheme 1.** Synthetic schemes of the various new probes reported in this study: (A) Chol-alk (Probe 1), (B) CS-DA (Probe 3), (C) CP-DA (Probe 5) and (D) Ch-DA (Probe 6). Complete details of all the synthesis and analytical characterization of the various intermediates can be found in the ***Supplementary Synthetic Note***.

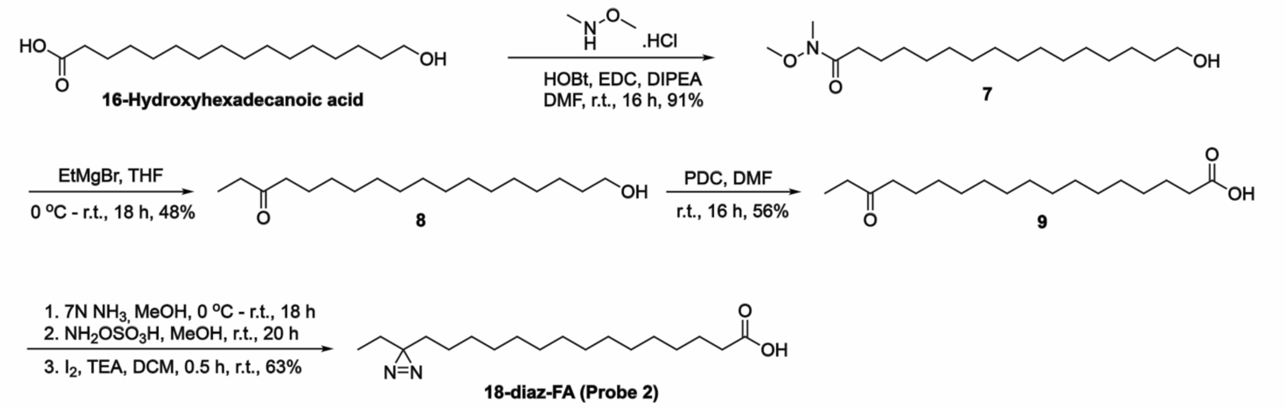


**Supplementary Scheme 2.** Synthesis of 18-diaz-FA (*Probe 2*) as per a reported protocol[1]. Complete details of all the synthesis and analytical characterization of the various intermediates can be found in the ***Supplementary Synthetic Note***.

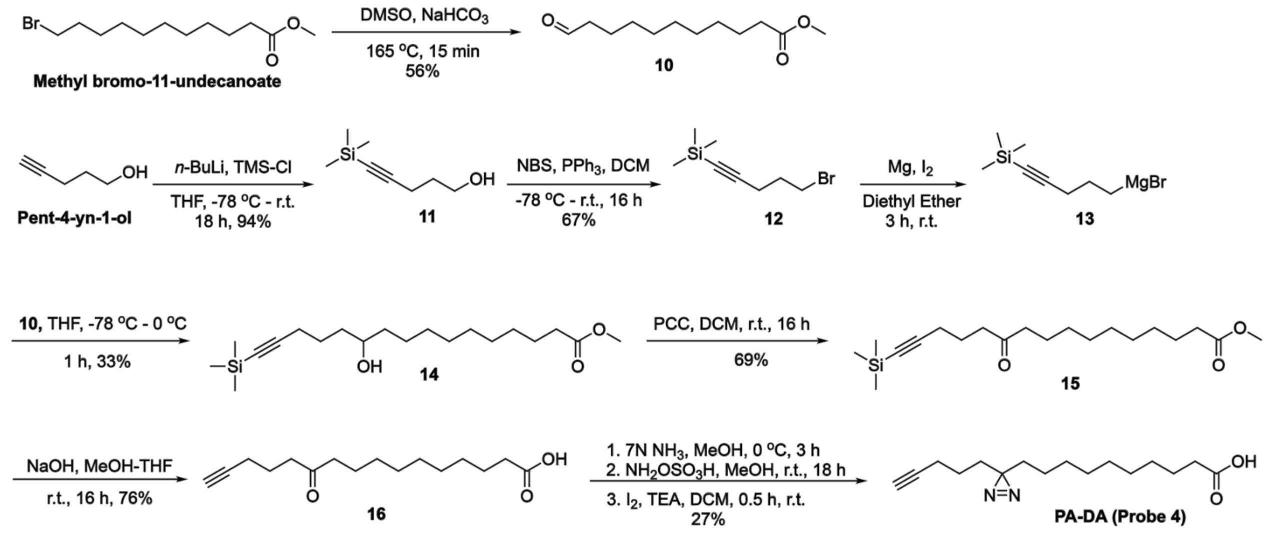


**Supplementary Scheme 3.** Synthesis of PA-DA (*Probe 4*) as per a reported protocol[2,3]. Complete details of all the synthesis and analytical characterization of the various intermediates can be found in the ***Supplementary Synthetic Note***.

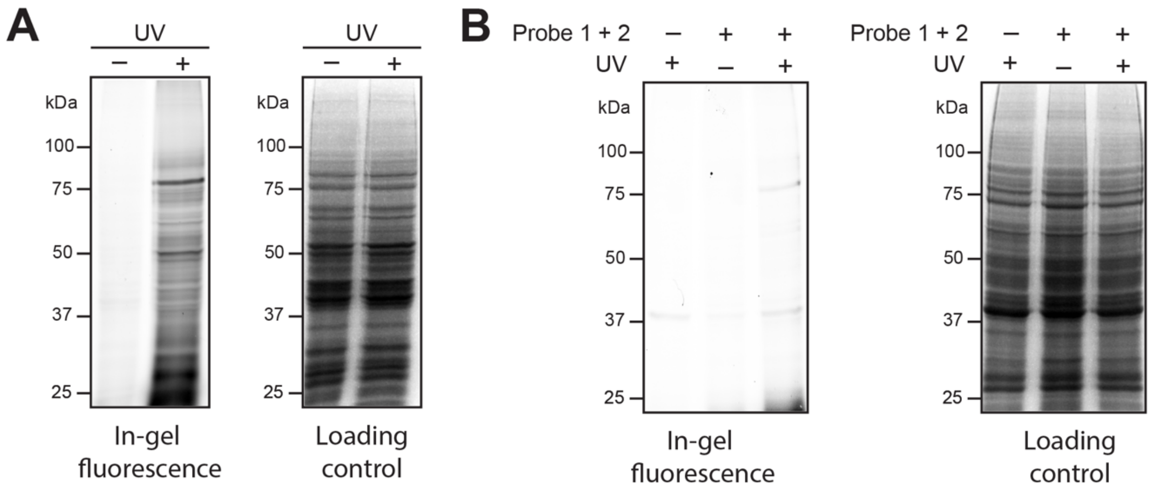


**Supplementary Figure 1.** A representative fluorescence gel from a gel based chemical proteomics experiment depicting: (**A**) UV-dependent photocrosslinking (protein labeling) in the metabolic labeling experiment when both probe 1 and probe 2 (200 μM each, 4 h) are added in RAW264.7 cells. (**B**) Negligible in-gel fluorescence observed in UV treated samples when both probe 1 and probe 2 (200 μM each, 4 h) are added to Neuro2A cells due to low intrinsic ACAT activity, confirming the requirement for ACAT activity for the metabolic assembly of probe 3 in cells. For (**A, B**): This experiment was done three times with reproducible results each time.

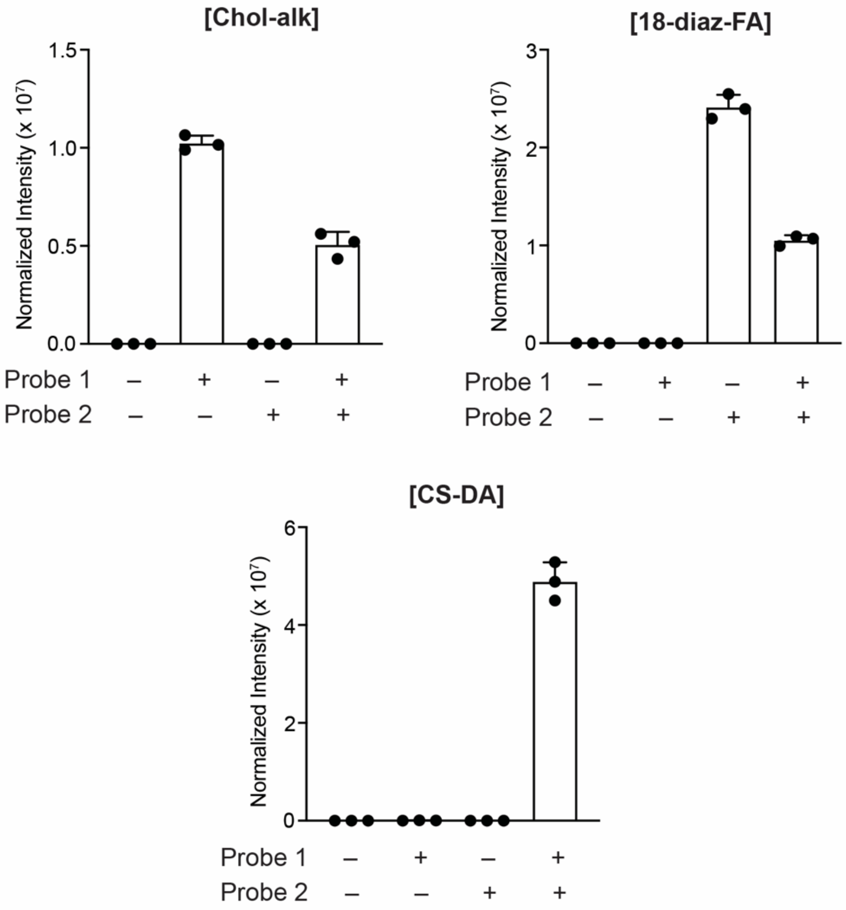


**Supplementary Figure 2.** A semi-quantitative LC-MS based lipidomics experiment showing the uptake and consumption of Chol-alk (Probe 1) and 18-diaz-FA (Probe 2) extracted from treated RAW264.7 cells (200 μM each, 4 h) and the formation of CS-DA (Probe 3) only when both probes are added. This experiment was done three times with reproducible results each time.

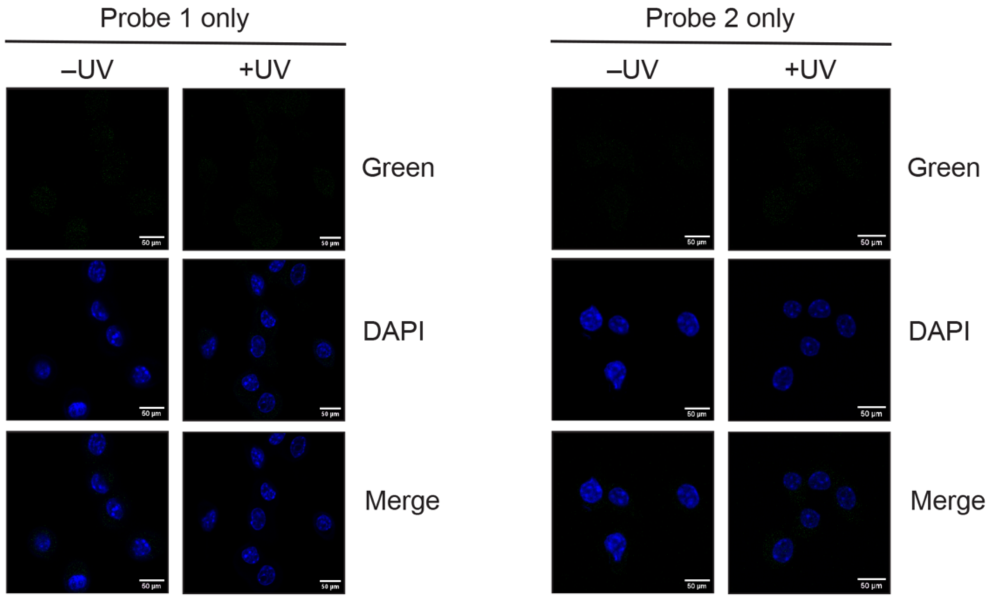


**Supplementary Figure 3.** Representative fluorescence microscopy images showing no cellular fluorescence from Probe 1 and Probe 2 (green channel) when both probes (200 μM, 4 h) are added individually to RAW264.7 cells in both UV-treated samples and the non-UV irradiated controls. DAPI (blue channel) was used in these fluorescence microscopy experiments to mark the nucleus in individual cells. The scale bars on each panel are 50 μm in length. This experiment was done three times with reproducible results each time.

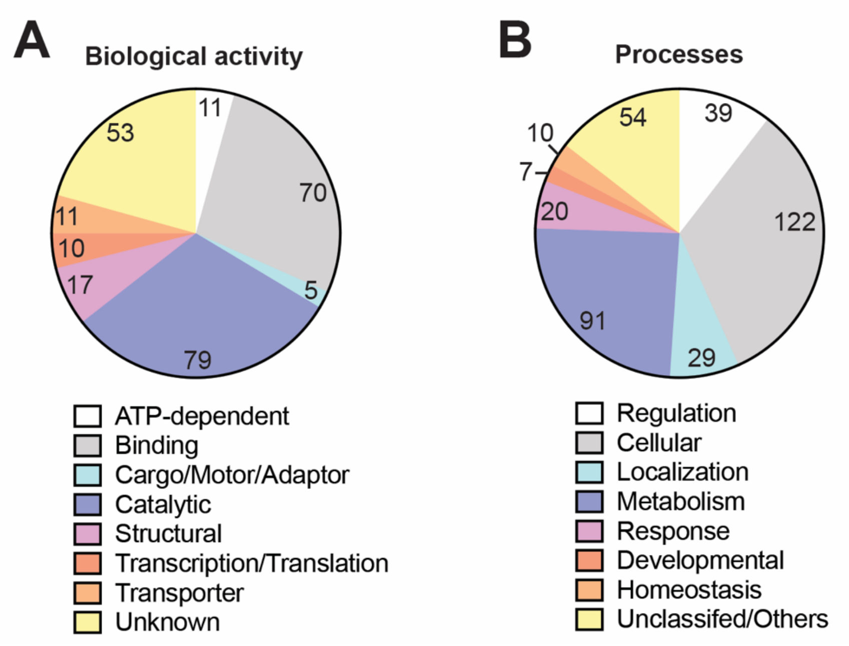


**Supplementary Figure 4.** Categorization of "hits" from the metabolically synthesized Probe 3 into: (**A**) known biological activities; and (**B**) physiological processes they are involved in, based on the Panther database classification[4,5].

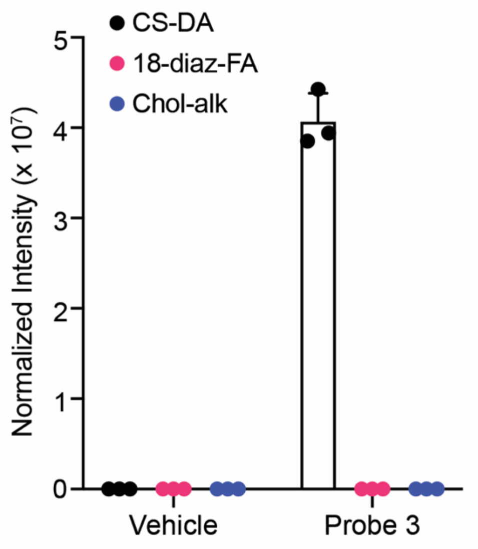


**Supplementary Figure 5.** A semi-quantitative LC-MS based lipidomics experiment showing the uptake CS-DA (Probe 3) extracted from RAW264.7 cells only when treated with Probe 3 (50 μM probe, 4 h treatment) with no signal in vehicle treated controls and no formation of hydrolytic degradation products [Chol-alk (Probe 1) and 18-Diaz-FA (Probe 2)] in Probe 3 treated cells. This experiment was done three times with reproducible results each time.

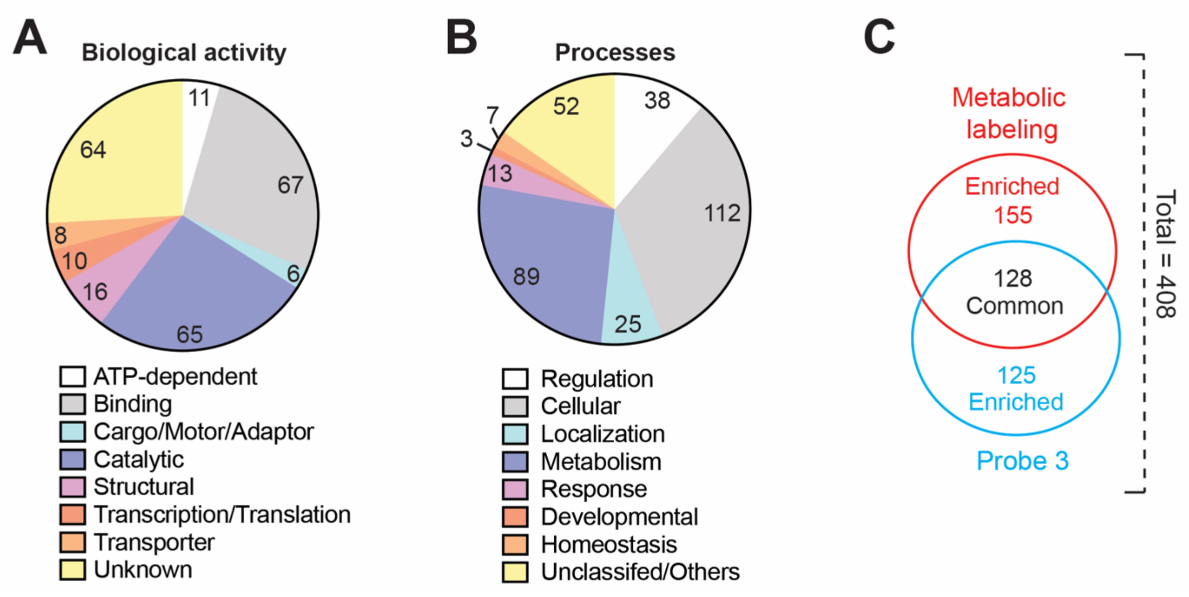


**Supplementary Figure 6. PANTHER Classification and comparison of Probe 3 hits with metabolic labelling.** Categorization of “hits” when Probe 3 was directly fed to RAW264.7 cells into: (**A**) known biological activities; and (**B**) physiological processes they are involved in based on the Panther database classification[4,5]. (**C**) Venn diagram depicting shared and unique protein “hits” of metabolic labeling experiment and Probe 3 feeding experiment respectively.

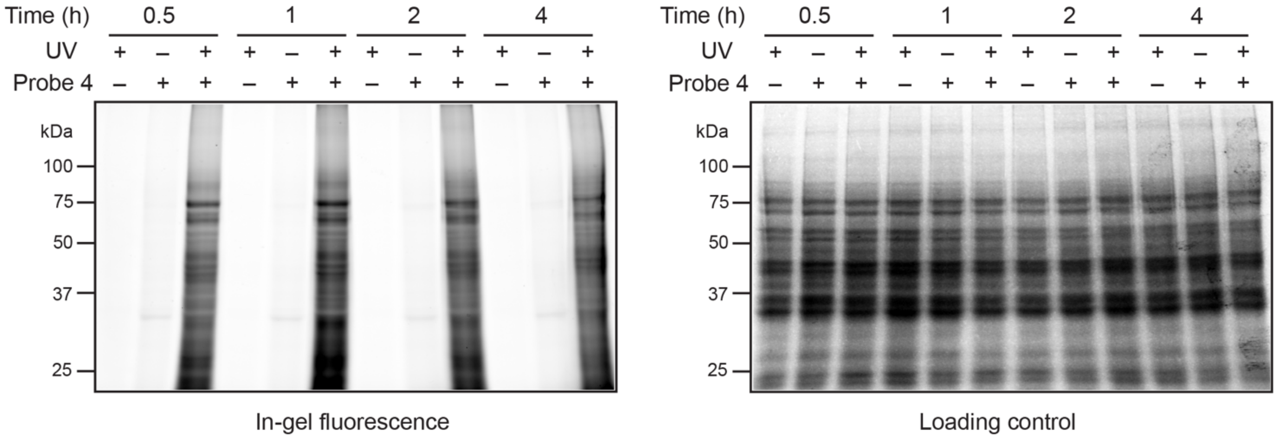


**Supplementary Figure 7.** A representative fluorescence gel from a time-course experiment showing UV-dependent protein labeling when Probe 4 (50 μM) is fed to RAW264.7 cells from 30 min to 4 h, with optimal uptake and probe labeling observed at all timepoints. The Coomassie staining shows equal protein loading for all time points in this experiment. This experiment was done three times with reproducible results each time.

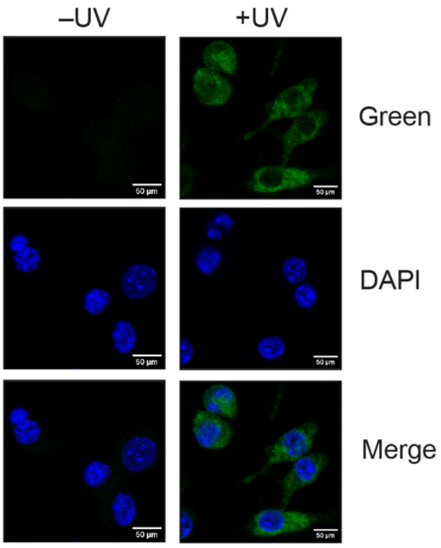


**Supplementary Figure 8.** Representative fluorescence microscopy images showing cellular fluorescence for Probe 4 (green channel) fed to RAW264.7 cells (50 μM, 30 min) only in the presence of UV. In the same experiment, no fluorescence (green channel) was observed in the non-UV irradiated controls. DAPI (blue channel) was used in these fluorescence microscopy experiments to mark the nucleus in individual cells. The scale bars on each panel are 50 μm in length. This experiment was done three times with reproducible results each time.

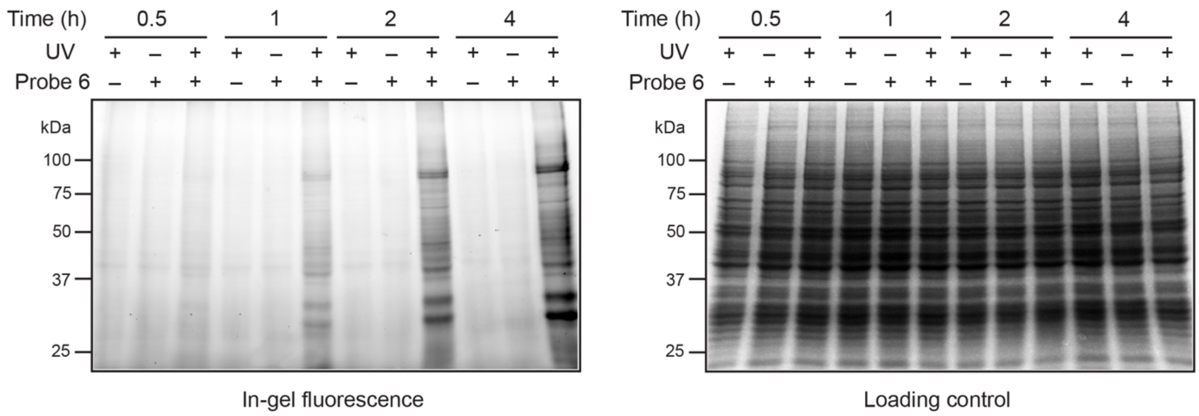


**Supplementary Figure 9.** A representative fluorescence gel from a time-course experiment showing linear time-dependent increase in protein labeling when Probe 6 (50 μM) is fed to RAW264.7 cells from 30 min to 4 h, with maximum labeling observed at 4 h timepoint. The Coomassie staining shows equal protein loading for all time points in this experiment. This experiment was done three times with reproducible results each time.

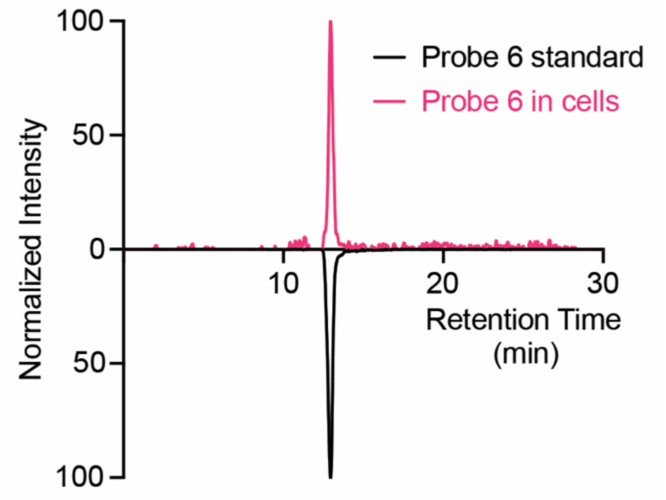


**Supplementary Figure 10.** A LC-MS experiment showing the elution profile of Probe 6 (50 μM, 4 h treatment) extracted from treated RAW264.7 cells co-eluting with the Probe 6 synthesized standard at the same retention time. This experiment was done three times with reproducible results each time.

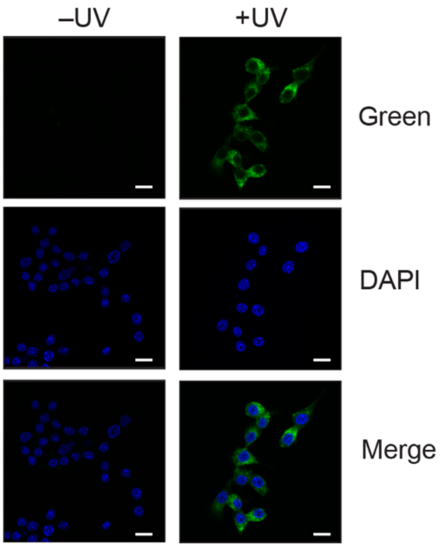


**Supplementary Figure 11.** Representative fluorescence microscopy images showing cellular fluorescence for Probe 6 (green channel) fed to RAW264.7 cells (50 μM, 4 h) only in the presence of UV. In the same experiment, no fluorescence (green channel) was observed in the non-UV irradiated controls. DAPI (blue channel) was used in these fluorescence microscopy experiments to mark the nucleus in individual cells. The scale bars on each panel are 50 μm in length. This experiment was done three times with reproducible results each time.

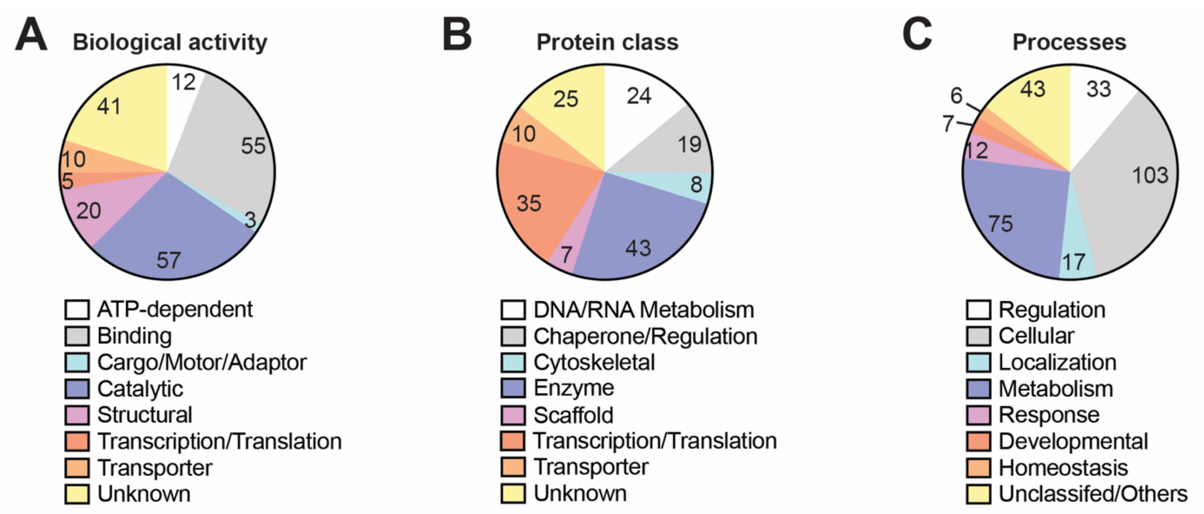


**Supplementary Figure 12.** Categorization of unique “hits” (224 Proteins) from Probe 5 into: (**A**) known biological activities; and (**B**) protein classes, and (**C**) physiological processes they are involved in, based on the Panther database classification[4,5].

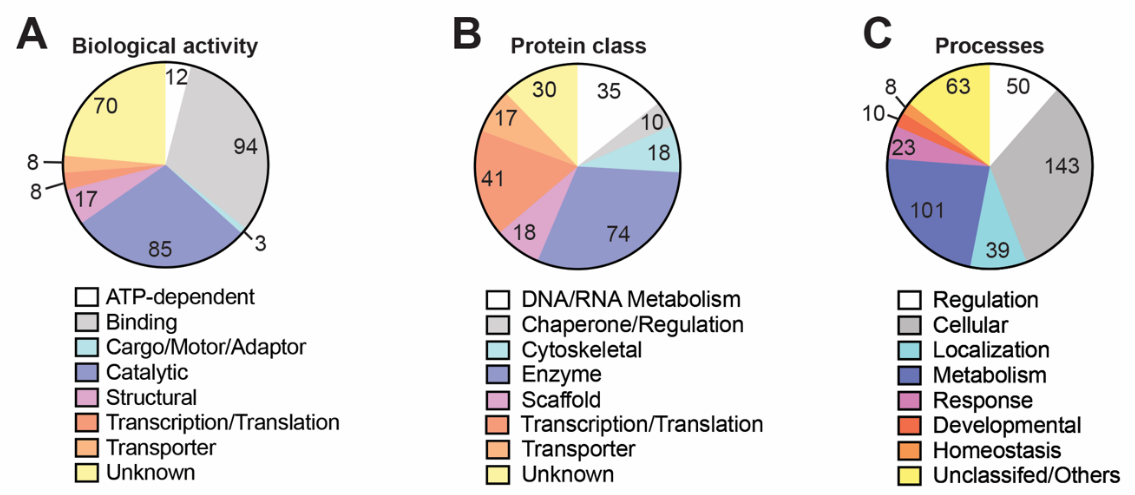


**Supplementary Figure 13.** Categorization of unique "hits" (356 Proteins) from Probe 6 into: (**A**) known biological activities; and (**B**) protein classes, and (**C**) physiological processes they are involved in, based on the Panther database classification[4,5].

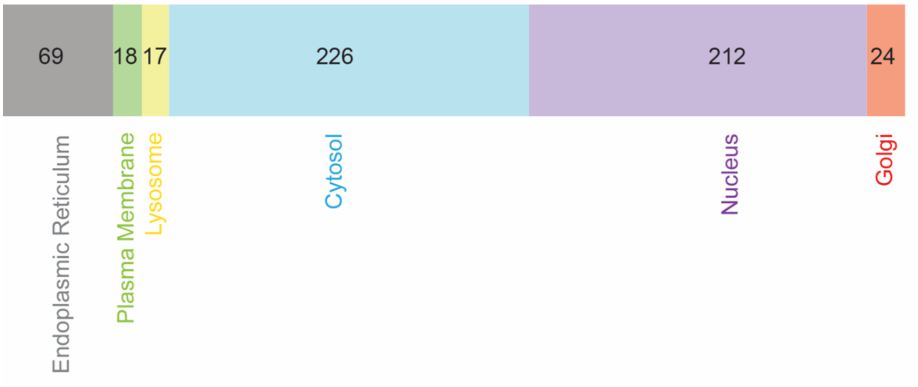


**Supplementary Figure 14.** Subcellular localization of total CE "hits" (495 proteins in total) obtained from all three orthogonal strategies based on the UniProt annotation[6].

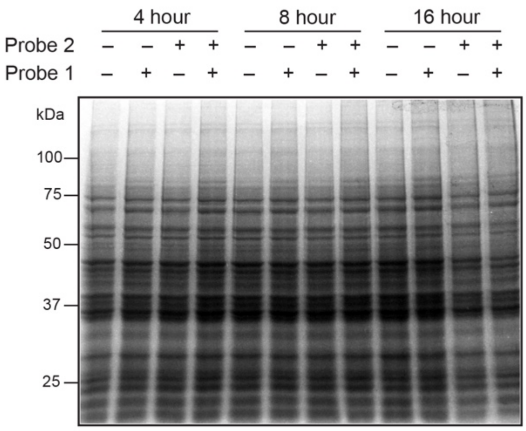


Loading control (Coomassie staining) for **Figure 2B**.

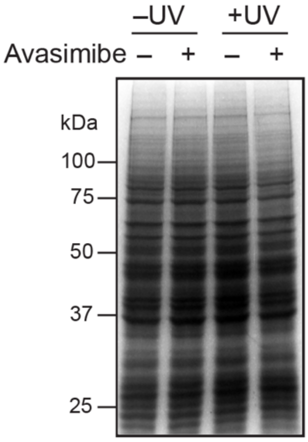


Loading control (Coomassie staining) for **Figure 2C**.

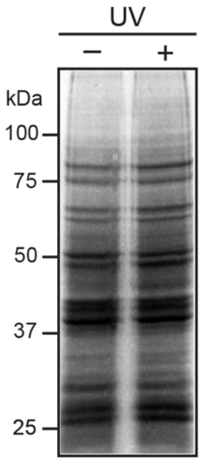


Loading control (Coomassie staining) for **Figure 3A**.

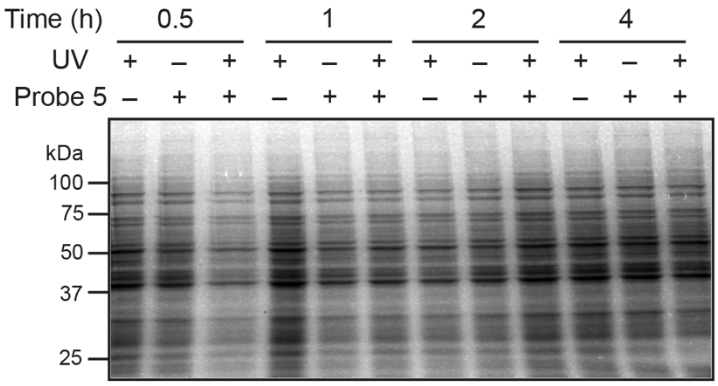


Loading control (Coomassie staining) for **Figure 4B**.

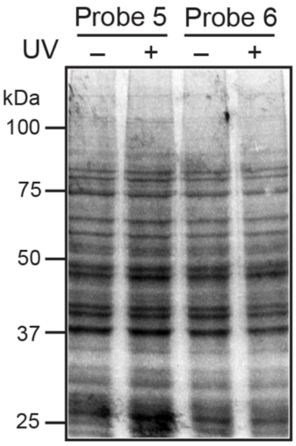


Loading control (Coomassie staining) for **Figure 4E**.
